# Transcriptome analysis provides insights into candidate genetic regulators of birch response to salt stress

**DOI:** 10.64898/2026.10.29.762967

**Authors:** Zhi Li, Qingjing Huang, Liangju Li, Lixiu Yan, Bangmeng Zhou, Jie Cheng

## Abstract

**Background:** Soil salinization is a worldwide matter of concern and can result in a series of agricultural and ecological problems. Previous transcriptome data have shown genetic regulatory mechanisms of salt stress response in *a salt-tolerant species of Betula halophila and a salt-sensitive species of B. platyphylla*, but comparative analysis of transcriptional changes in the the two species remains limited.

**Aims:** We compared gene expression in leaves of *B. halophila* and leaves and roots of *B. platyphylla*.

**Methods:** Differentially expressed genes (DEGs) were identified after 24 h of salt stress, followed by Gene Ontology enrichment and gene regulatory network (GRN) construction. Selected DEGs were examined by quantitative real-time PCR (RT-qPCR).

**Results:** Compared with 0 h, 956 DEGs were found in *B. halophila* leaves (BHLeaf), 2,955 in *B. platyphylla* leaves (BPLeaf), and 3,827 in *B. platyphylla* roots (BPRoot). BHLeaf DEG expression patterns were similar to those of BPRoot. DEGs from BHLeaf and BPRoot were enriched in cell wall organization or biogenesis, whereas hormone biosynthesis and signaling genes were altered mainly in *B. platyphylla*. RT-qPCR showed that selected DEGs were induced by 200 mM NaCl for 24 h in *B. platyphylla*.

**Conclusions:** Our work uncovered genetic mechanisms of salt stress response commonly and uniquely occurring in two birch species and provides resources for breeding salt-tolerant birch.

**Highlights:**

- Salt stress altered the expression of 956, 2,955 and 3,827 genes in the two birch species.
- BHLeaf expression patterns were similar to those of BPRoot.
- The expression changes of nine selected DEGs were examined by RT-qPCR in *B. platyphylla*.

## Introduction

More than 20% of the world’s irrigated lands are threatened by high salinity, which is dramatically increasing every year due to climate change and unsustainable irrigation management (Velasco et al. 2018; Singh 2021). High salinity is one of the environmental stressors that severely affects all aspects of plant growth and development (Xiao and Zhou 2023). To cope with stress problems in salt-affected problematic soils, plants have evolved diverse physiological and molecular mechanisms during the long period of evolution (Zhao et al. 2020b; Rahman et al. 2022; Thinn et al. 2025). Owing to the degree of tolerance to salt stress, land plants can be mainly divided into glycophytes and halophytes (Flowers et al. 2015). For glycophytes, the general limit of external salt concentrations above which plant yield is severely reduced is approximately 86 mM sodium chloride (NaCl) (Chapman 1942), while halophytes are only observed for their yield penalties when the external salt concentrations exceed 200 mM NaCl (Flowers and Colmer 2008; Santos et al. 2016). The halophytic plants which grow in salt-alkalinized habitats are truly salt-tolerant plants, which are provided with the capability of excluding, excreting, and accumulating salts into/from their cells, organelles, and tissues (Shabala 2013; Flowers et al. 2015; Percey et al. 2016). Moreover, external salt concentrations ranged from 40 to 100 mM NaCl severely inhibit photosynthesis in glycophytes (Munns and Tester 2008), while many halophytes are capable of maintaining or improving photosynthesis at these salt concentrations (Stepien and Johnson 2009). According to the salt-tolerant characteristics of halophytes mentioned earlier, it is reasonable for us to select them as ideal models to elucidate the mechanisms that underlie salinity tolerance (Fan 2020).

In the past decades, it has been well document that plants responding to salt stress involve a variety of biological processes regulated by different functional components, mainly including osmoregulation, ionic homeostasis, oxidation-reduction regulation, antioxidant enzyme activation, as well as the biosynthesis of hormone and compatible solutes (Munns and Tester 2008; Velasco et al. 2018; Zhao et al. 2020b; Singh 2021). However, the elucidation of contrasting mechanisms underlying salt stress response between halophytes and glycophytes are far from being completely understood. Therefore, it is necessary for us to screen the salt tolerant gene alleles and transform them for the purpose of obtaining salinity-resistant plants with strong acclimation and adaptation to extreme saline environments (Fang et al. 2021).

Birch trees (*Betula spp.*), widely distributed across Europe and parts of Asia, can thrive in a wide range of habitats from boreal forests to temperate woodlands (Jonczak et al. 2020). The ecological, economic, and evolutionary values of birch trees (Gang et al. 2019), together with their roles in scientific researches, underscore the need for thorough and systematic studies into their growth and developmental properties under normal and stressful conditions. *Betula halophila*, a critically endangered halophytic species, is first discovered in a swamp co-occurred with extremely high salinity located in Xinjiang province in 1956 (Shao et al. 2018b). *B. platyphylla*, a pioneer hardwood tree species, has exceptional cold tolerance and weak salt resistance (Liu et al. 1998; Chen et al. 2021a). In the present study, we compared the expression alterations of genes across leaves and roots of *B. platyphylla* as well as leaves of *B. halophila*. Based on the analysis of differentially expressed genes (DEGs), we proposed the gene regulatory networks (GRNs) to observe how birch seedlings responded to high saline conditions. Furthermore, we randomly selected nine genes from our constructed GRNs and confirmed their expression levels using quantitative real-time polymerase chain reaction (RT-qPCR) assay. Our transcriptome analysis explored the genetic regulatory mechanisms involved in salt stress response and discovered a total of potential genes for enhancing salt tolerance in birch, which could lay the foundation for decoding distinct adaptation mechanisms of glycophytes to saline-alkali lands.

## Materials and methods

### Transcriptome data collection

To understand the transcriptional dynamics of birch seedlings in response to salt stress, we respectively downloaded the transcriptome data of *B. halophila* (Shao et al. 2018a) and *B. platyphylla* (Jia et al. 2022) from Sequence Read Archive (SRA) data on the National Center for Biotechnology Information (NCBI) website. Of which, the leaves from *B. halophila* as well as the leaves and roots from *B. platyphylla* were all subjected to 0 mM NaCl for 0 h and 200 mM NaCl for 24 h. Three biological replicates were conducted. In our analysis, the leaves under 200 mM NaCl of *B. halophila* were designated as BHLeaf. The leaves and roots under 200 mM NaCl of *B. platyphylla* were respectively designated as BPLeaf and BPRoot.

### Transcriptome data analysis

We firstly discarded the adaptor sequences and reads with unknown or low-quality bases from raw reads using fastp v0.23.2 (Chen et al. 2018) to obtain clean reads. Clean reads were then mapped to the genome of *B. pendula subsp. pendula* v.1.2 (Salojärvi et al. 2017) using HISAT2 v2.2.1 (Kim et al. 2019). The transcripts assembly and quantitation were evaluated using StringTie v2.1.3b (Pertea et al. 2016). The gene expression levels were calculated by fragments per kilo-bases per million mapped reads (FPKM) values. We used DESeq2 package (Love et al. 2014) to acquire the differentially expressed genes (DEGs). The criterion of false discovery rate (FDR)-corrected P value of 0.05 and an expression value of log |fold change|>1 was used for judging the significance of expression differences (Benjamini and Yekutieli 2001). Classical Venn diagram was constructed using EVenn online software (Chen et al. 2021b). Heatmaps of gene expression patterns were carried out by pheatmap package v1.0.12 (https://cran.r-project.org/package=pheatmap). Transcription factors (TFs) were annotated and classified by the iTAK online software (Zheng et al. 2016).

### The construction of gene regulatory networks (GRNs)

In our work, genes differentially expressed at one or more tissues from two species were regarded as a gene set. Based on the Short Time-series Expression Miner (STEM) clustering method, clusters were divided by STEM software v1.3.13 (Ernst and Bar-Joseph 2006) with default parameters. Further, we employed the GEne Network Inference with Ensemble of trees (GENIE3) package to construct the potential GRNs related to salt stress. After that, the GRNs were visualized using Cytoscape v.3.10.2 (Shannon et al. 2003).

### Gene ontology (GO) enrichment analysis of DEGs

Through homolog searches, we conducted the functional annotation of genes in birch genome. The BLASTP v2.7.1 program was used for searching the Uniprot/SwissProt Release 2021_01 (10-Feb-2021) database with parameters including-evalue 1e-5,-num_alignments 1, and - outfmt 6. The BLASTP hits with a percentage of identical matches lower than 60% and an alignment length shorter than 70 bp were filtered out. The unique functional annotation of these filtered BLASTP results was regarded as the non-redundant functional annotation database. GO enrichment analysis was carried out using the clusterProfiler package v3.18.0 (Yu et al. 2012).

### The prediction of homologous genes related to salt stress in birch

Genome files from *Arabidopsis thaliana* (TAIR10) were downloaded from the Phytozome v13 online website (Goodstein et al. 2012). Genome files of *B. pendula* v1.2 scaffolds (Salojärvi et al. 2017) were retrieved from the CoGe comparative genomics platform (https://genomevolution.org/CoGe). After that, we performed the OrthoFinder v2.2.7 software (Emms and Kelly 2015) to identify homology relationships between birch and *Arabidopsis* proteins. In combination with the information of *Arabidopsis* genes related to salt stress signaling, response, and tolerance reported in public articles, we predicted the birch candidate genes involved in salt stress response.

### Expression patterns of candidate genes related to salt stress response using RT-qPCR assay

Micro-propagated plants of *B. platyphylla* were grown on 6‰ agar-solidified woody plant medium (WPM) with 3.0% sucrose under the pH=5.8. The roots of seedlings were created by WPM medium with the supplement of 1.0 mg/L naphthalene acetic acid (NAA) for 30 d. The culture room parameters were set as follows: 25°C, 16 h of light per day, and light density of approximately 5000 lux. After that, the rooted seedlings were cultivated in the substrates mixed with soils, vermiculites, and perlites for two months under the standard greenhouse conditions. Young seedlings were subjected to 200 mM NaCl solutions for 0 and 24 h. At every time-point, roots and the third leaves close to the apical buds were respectively harvested. Six biological replicates were conducted.

RNA extraction and reverse transcription assays were completed according to the protocols described previously (Bian et al. 2025). We randomly selected nine candidate genes related to salt stress response and examined the expression levels of these genes using quantitative real-time polymerase chain reaction (RT-qPCR) assay. RT-qPCR assay was accomplished by the PrimerScript RT Reagent Kit (Perfect Real Time, Takara) on a LightCycler^®^ 480 II instrument (Roche Diagnostics, Indianapolis, USA). *18S rRNA* (GenBank Accession #: FJ228477.1) was used as a housekeeping gene. The specific primers of genes used for RT-qPCR assay were listed in Table 1. Relative expression levels were calculated using the 2^−ΔΔCt^ method (Schmittgen and Livak 2008). Three replicates were performed for each sample.

**Table 1.**
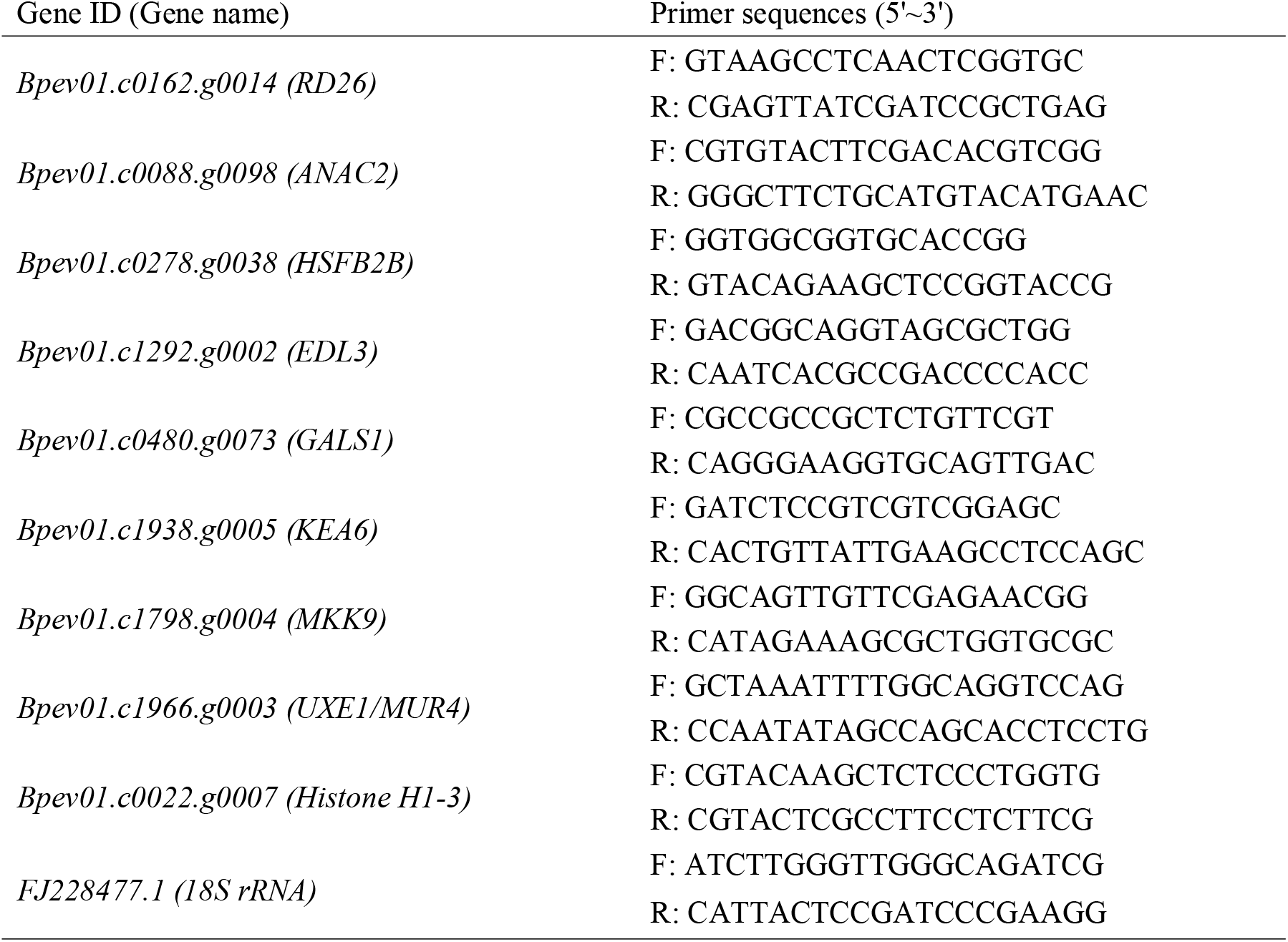
Primer sequences used for RT-qPCR assay.

## Results

### Analysis of genes differentially expressed in *B. halophila* and *B. platyphylla*

To study the transcriptional dynamics under salt stress in birch, we downloaded and analyzed the transcriptome data reported by previous studies (Shao et al. 2018a; Jia et al. 2022). Compared to 0 h, there were 339 up-regulated genes and 617 down-regulated genes in BHLeaf after salt stress for 24 h, 1784 up-regulated genes and 1171 down-regulated genes in BPLeaf after salt stress for 24 h, and 2320 up-regulated genes and 1507 down-regulated genes in BPRoot after salt stress for 24 h (Fig. 1a). These results suggested that after salt treatment, the number of genes differentially expressed in the leaves of *B. halophila* was similar to that of roots in *B. platyphylla*, while the number of genes with altered expression levels from leaves of two birch species were greatly differed.

**Fig. 1.**
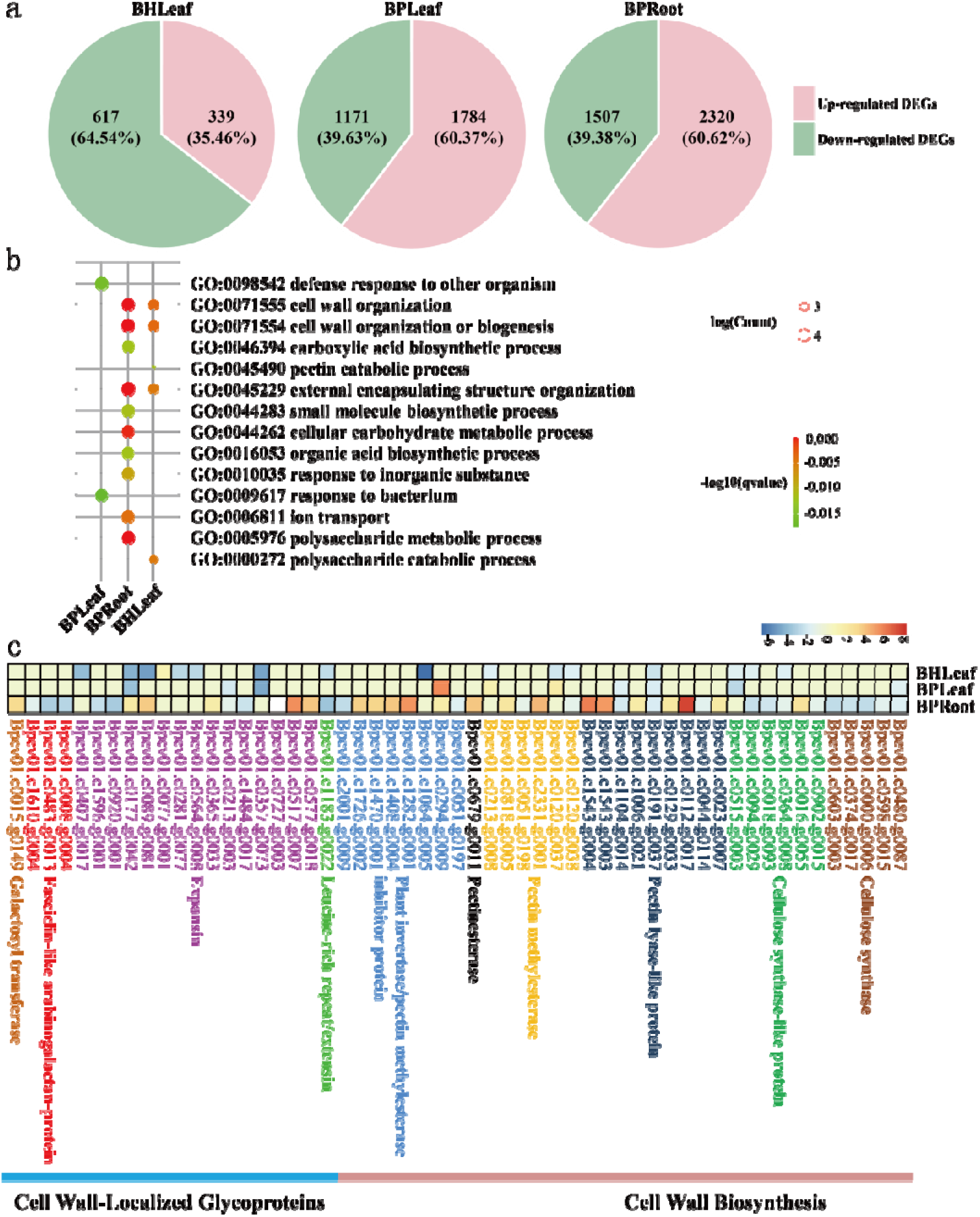
Analysis of DEGs in *B. halophila* and *B. platyphylla*. a, Number of up-regulated and down-regulated genes in BHLeaf, BPLeaf, and BPRoot. b, GO enrichment analysis of DEGs in BHLeaf, BPLeaf, and BPRoot. The 10 most represented GO terms are shown. c, Expression patterns of DEGs significantly enriched in cell wall organization or biogenesis.

### Genes related to cell wall organization or biogenesis occurred in leaves of B. halophila and roots of *B. platyphylla*

GO enrichment analysis showed that many DEGs from BHLeaf were significantly enriched in cell wall organization or biogenesis (GO:0071554), cell wall organization (GO:0071555), polysaccharide catabolic process (GO:0000272), external encapsulating structure organization (GO:0045229), and pectin catabolic process (GO:0045490) (*q-value* <0.05) (Fig. 1b). Many DEGs from BPLeaf were significantly involved in defense response to other organism (GO:0098542) and response to bacterium (GO:0009617) (*q-value* <0.05) (Fig. 1b). Many DEGs from BPRoot were significantly related to cell wall organization or biogenesis (GO:0071554), polysaccharide metabolic process (GO:0005976), ion transport (GO:0006811), external encapsulating structure organization (GO:0045229), cell wall organization (GO:0071555), response to inorganic substance (GO:0010035), small molecule biosynthetic process (GO:0044283), cellular carbohydrate metabolic process (GO:0044262), organic acid biosynthetic process (GO:0016053), and carboxylic acid biosynthetic process (GO:0046394) (*q-value* <0.05) (Fig. 1b). Accordingly, it seemed that the cell wall organization or biogenesis was a common biological process in the leaves of *B. halophila* and roots of *B. platyphylla* exposed to 200 mM NaCl solutions.

To further study the DEGs significantly enriched in cell wall organization or biogenesis, we performed the observation on the enzymatic products generated by these DEGs. In our study, these genes could be classified into two groups (Fig. 1c). Specifically, genes encoding the cellulose synthase, cellulose synthase-like protein, pectin lyase-like protein, pectin methylesterase, pectinesterase, and plant invertase/pectin methylesterase inhibitor protein were associated with the cell wall biosynthesis. Genes encoding leucine-rich repeat/extension, expansin, fasciclin-like arabinogalactan-protein, and galactosyl transferase were in relation to cell wall-localized glycoproteins. In most cases, the expressions of these genes mentioned earlier showed a decrease in leaves of *B. halophila*, while there was an increased expressions of these genes in leaves and roots of *B. platyphylla*. As a result, it appeared that the opposite expression levels of genes involved in cell wall organization or biogenesis were present in two birch species.

### Expression profiles of genes from leaves of B. halophila shared similarity to roots of *B. platyphylla*

In the present study, a series of DEGs detected at one or more tissues of two birch species were regarded as a dataset, which was further grouped into 10 clusters by STEM software using the STEM clustering method algorithm (Fig. 2). Among them, the expression profiles of genes from cluster 3, 4, 5, 6, and 9 reached the significant difference (*p* < 0.05). Genes from cluster 9 were up-regulated in all tissues of two species, and the expression folds of these genes were increased in leaves and roots of *B. platyphylla* when compared to the leaves of *B. halophila*. With regards to cluster 3, 5, and 6, the expression levels of genes from leaves of *B. halophila* showed similarity to that of roots of *B. platyphylla*. Besides, genes from cluster 4 exhibited the opposite expression patterns between the tissues of two birch species. Therefore, we proposed that the expression profiles of genes in leaves of *B. halophila* were similar to that of the roots of *B. platyphylla*, while the expression profiles of genes from leaves of two birch species were greatly differed.

**Fig. 2.**
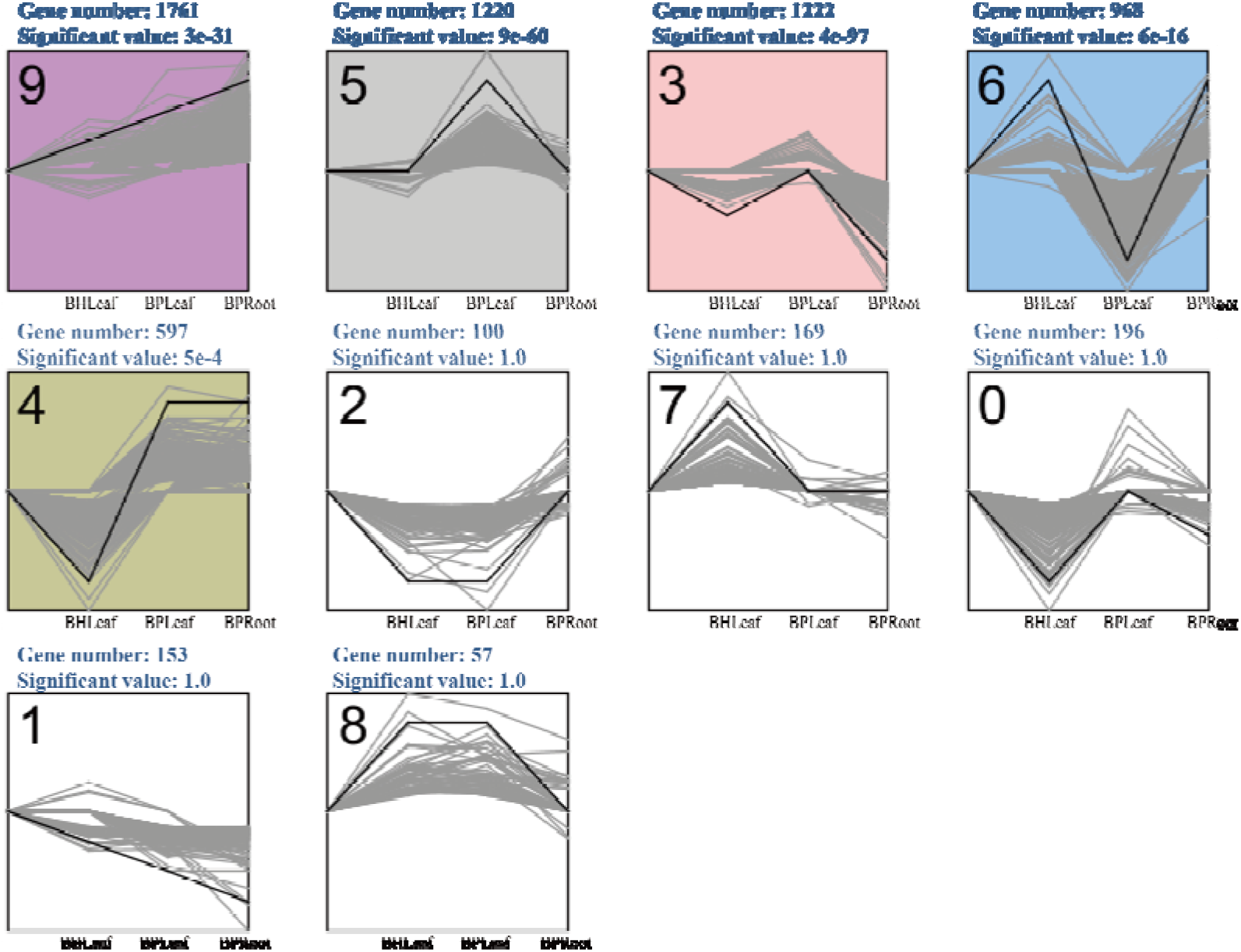
Cluster analysis of DEGs detected in one or more tissues of *B. halophila* and *B. platyphylla*. The expression levels were represented by log_2_FC.

To further understand the genes from a certain cluster which biological processes participated in, we performed GO enrichment analysis (Table 2). In our work, genes from the cluster 3 were mainly involved in cell wall organization or biogenesis (*q-value* < 0.05). Genes from the cluster 5 were mainly implicated in ribosome biogenesis, response to wounding, as well as defense response to other organism and fungus (*q-value* < 0.05). Genes from the cluster 6 were mainly related to plastid organization, chloroplast organization, and chloroplast RNA processing (*q-value* < 0.05). Genes from the cluster 9 were mainly in relation to photosynthesis, oxylipin biosynthetic and metabolic process, abscisic acid (ABA) response, inorganic substance, and water deprivation (*q-value* < 0.05).

**Table 2.** GO enrichment analysis of genes in each cluster divided by STEM software.

### Genes responsible for hormone biosynthesis and signaling showed altered expression in *B. platyphylla* different from *B. halophila*

On account of the *Arabidopsis* genes associated with salt stress response, signaling, and tolerance reported in public articles, we found 485 homologous genes from birch genome and 224 genes displayed altered expression in one or more tissues of two species after salt stress for 24 h when compared to 0 h (Table 3 and Fig. 3). These results might imply the functional roles of these homologous genes in controlling salt stress response in birch plants. Based on the expression patterns of these genes in two species, we tended to observe whether specific biological processes occurred during salt stress.

**Fig. 3.**
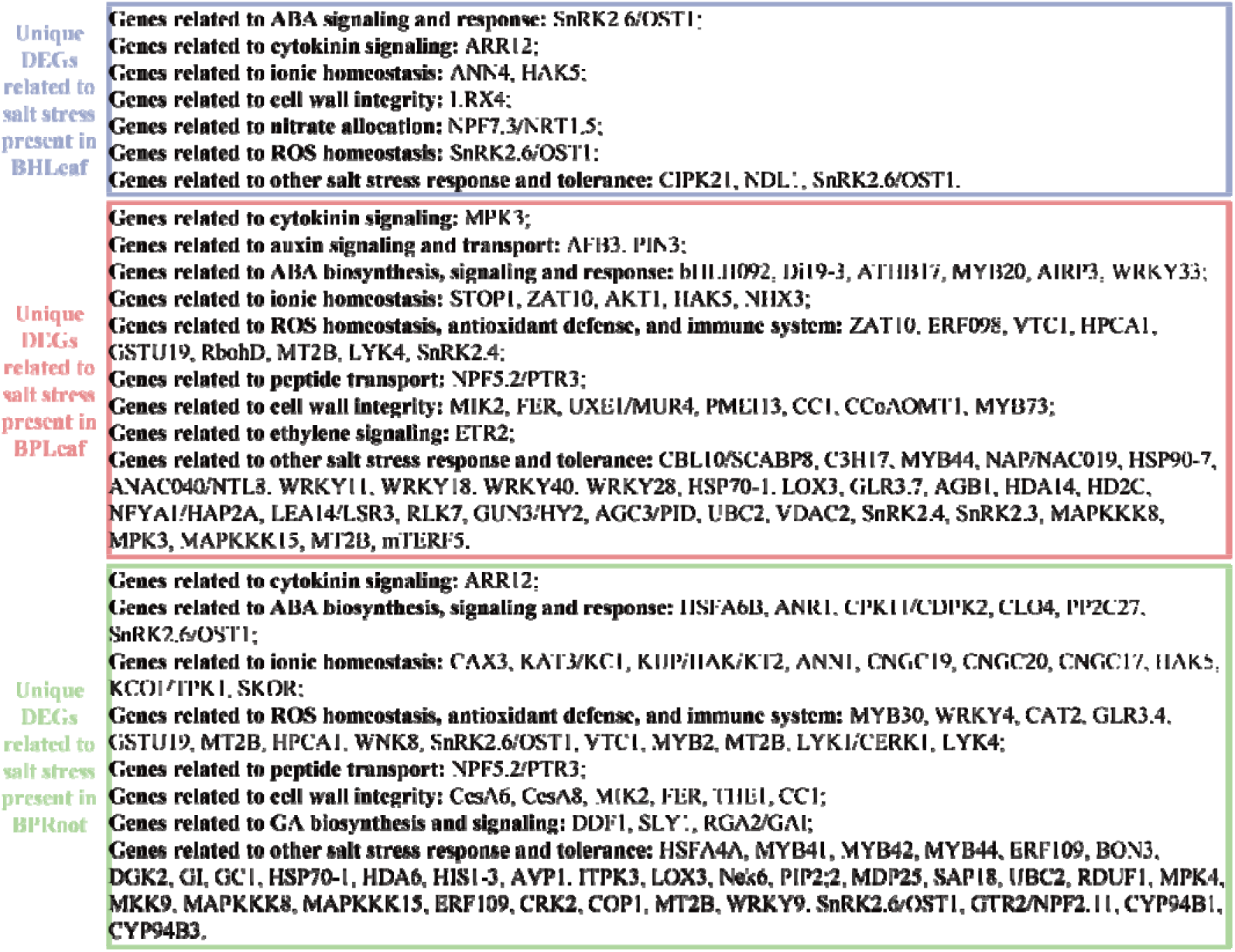
Analysis of DEGs uniquely identified in different tissues of *B. halophila* and *B. platyphylla*. Details of these genes are listed in Table S3.

**Table 3.** Genes related to salt stress response differentially expressed in *B. halophila* and *B. platyphylla*.

Of which, nine DEGs (four up-regulated genes and five down-regulated genes), 73 DEGs (52 up-regulated genes and 21 down-regulated genes), and 84 DEGs (53 up-regulated genes and 31 down-regulated genes) were uniquely present in BHLeaf, BPLeaf, and BPRoot, respectively. Furthermore, nine DEGs from BHLeaf were mainly involved in ABA signaling and response, cytokinin signaling, ionic homeostasis, cell wall integrity, nitrate allocation, and reactive oxygen species (ROS) homeostasis (Table 3 and Fig. 3). 21 DEGs from BPLeaf were mainly involved in cytokinin signaling, auxin signaling and transport, ABA biosynthesis, signaling and response, ionic homeostasis, ROS homeostasis, antioxidant defense, immune system, peptide transport, cell wall integrity, and ethylene signaling (Table 3 and Fig. 3). 84 DEGs from BPRoot were mainly involved in cytokinin signaling, ABA biosynthesis, ABA signaling, ABA response, ionic homeostasis, ROS homeostasis, antioxidant defense, and immune system, peptide transport, cell wall integrity, and gibberellins (GAs) biosynthesis and signaling (Table 3 and Fig. 3). As a result, we thought that the hormone biosynthesis, signaling, and response in most cases was altered in *B. platyphylla* instead of *B. halophila*.

### Inference of the unique molecular regulatory mechanisms responsible for salt stress response in two birch species

We further analyzed the DEGs between two birch species using a Venn diagram (Fig. 4). In our work, 121 DEGs were shared in BHLeaf, BPLeaf, and BPRoot. There were 444, 1877, and 2668 DEGs uniquely present in BHLeaf, BPLeaf, and BPRoot. These results possibly indicated a remarkable difference occurred in different tissues of two birch species subjected to salt treatment.

**Fig. 4.**
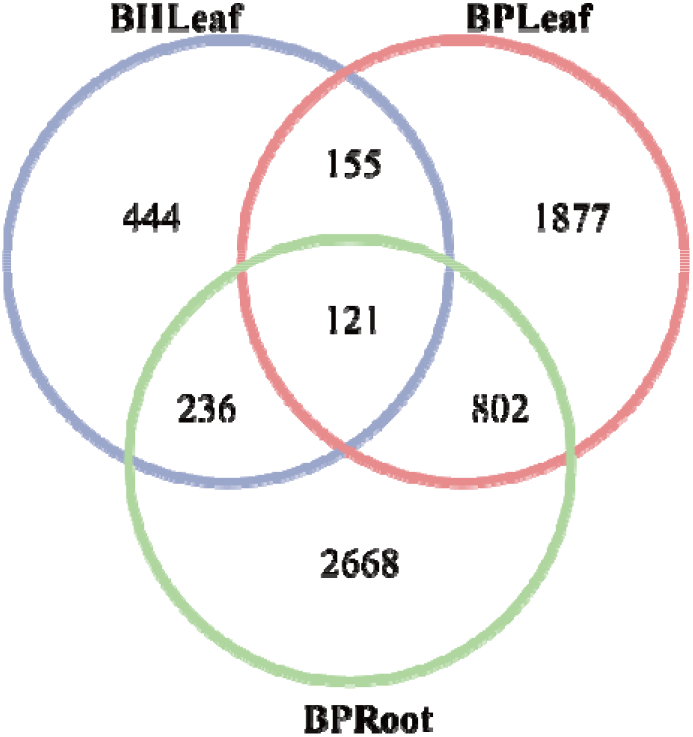
Venn diagram analysis of DEGs among different tissues of two birch species.

To uncover the candidate genes governing salt stress response in birch, we analyzed the up-regulated and down-regulated DEGs uniquely present in BHLeaf, BPLeaf, and BPRoot. Of which, we respectively found 299 (67.34%) down-regulated genes and 145 (32.66%) upregulated genes present in BHLeaf, 848 (45.18%) down-regulated genes and 1029 (54.82%) up-regulated genes present in BPLeaf, as well as 1150 (43.10%) down-regulated genes and 1518 (56.90%) up-regulated genes present in BPRoot (Fig. 5a).

**Fig. 5.**
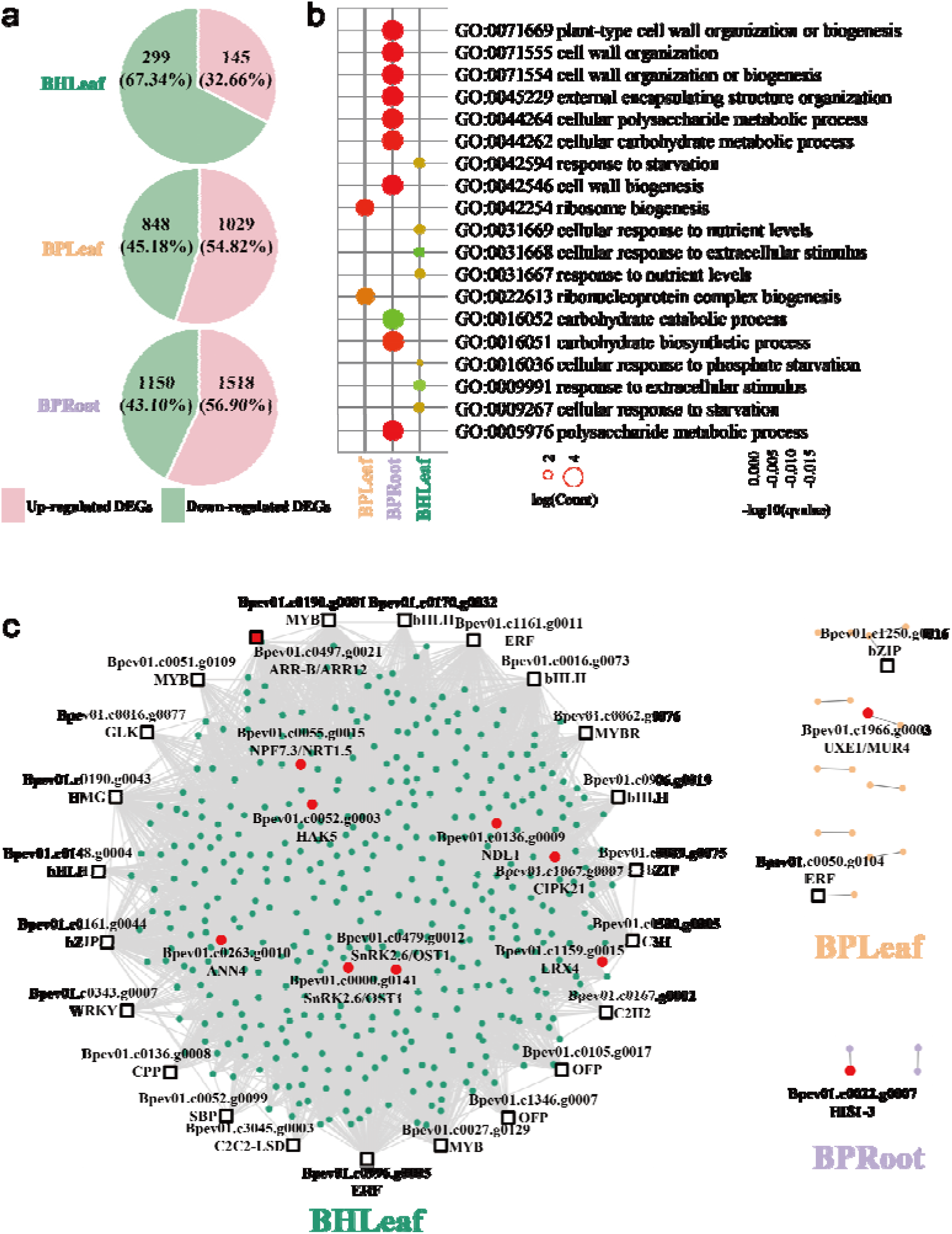
Analysis of DEGs uniquely present in *B. halophila* or *B. platyphylla*. a, Number of up-regulated and down-regulated genes uniquely present in BHLeaf, BPLeaf, and BPRoot. b, GO enrichment of DEGs uniquely present in BHLeaf, BPLeaf, and BPRoot. The 10 most represented GO terms are shown. c, Inferred GRNs involved in salt stress response of two birch species. The connectivity of gene to gene was screened by the threshold of weight values more than 0.01. The red circles represented birch genes showing the closest homology to *Arabidopsis* genes related to salt stress reported in public articles. The black squares represented TFs.

GO enrichment analysis showed that DEGs uniquely present in BHLeaf were significantly involved in cellular response to phosphate starvation (GO:0016036), cellular response to starvation (GO:0009267), response to nutrient levels (GO:0031667), cellular response to nutrient levels (GO:0031669), response to starvation (GO:0042594), response to extracellular stimulus (GO:0009991), and cellular response to extracellular stimulus (GO:0031668) (*q-value* < 0.05); DEGs uniquely present in BPLeaf were significantly related to ribosome biogenesis (GO:0042254) and ribonucleoprotein complex biogenesis (GO:0022613) (*q-value* < 0.05); and DEGs uniquely present in BPRoot were significantly involved in cell wall organization or biogenesis (GO:0071554), polysaccharide metabolic process (GO:0005976), external encapsulating structure organization (GO:0045229), cell wall organization (GO:0071555), cellular carbohydrate metabolic process (GO:0044262), carbohydrate biosynthetic process (GO:0016051), cellular polysaccharide metabolic process (GO:0044264), plant-type cell wall organization or biogenesis (GO:0071669), cell wall biogenesis (GO:0042546), and carbohydrate catabolic process (GO:0016052) (*q-value* < 0.05) (Fig. 5b).

Gene regulatory networks (GRNs) are powerful abstractions of biological systems (Huynh-Thu and Sanguinetti 2019). In this study, we adopted the GENIE3 software to construct the GRNs relevant for salt stress response with the parameters of weight values above 0.01 in two birch species (Fig. 5c, Table 4). The GRN of leaves from *B. halophila* was consisted of 444 genes. Among them, 9 genes showed the closest homology to *Arabidopsis* genes associated with salt stress response, signaling, and tolerance (Table 3), which included *Bpev01.c0000.g0141* (*SnRK2.6/OST1, Sucrose non-fermenting 1 (SNF1)-related protein kinase 2.6/Open stomata 1*), *Bpev01.c0052.g0003* (*HAK5, High affinity K^+^ transporter 5*), *Bpev01.c0055.g0015* (*NPF7.3/NRT1.5, Peptide transporter family 7.3/Nitrate transporter 1.5*), *Bpev01.c0136.g0009* (*NDL1, N-MYC downregulated-like 1*), *Bpev01.c0263.g0010* (*ANN4, ANNEXIN 4*), *Bpev01.c0479.g0012* (*SnRK2.6/OST1*), *Bpev01.c0497.g0021* (*ARR12, Arabidopsis response regulator 12*), *Bpev01.c1067.g000*7 (*CIPK21, Calcineurin B-like protein (CBL)-interacting protein kinase 21*), and *Bpev01.c1159.g0015* (*LRX4, Leucine-rich repeat extensin 4*). These results implied that genes discovered from our GRN of *B. halophila* probably acted as potential factors in regulating salt tolerance of birch trees.

**Table 4.** One inferred GRN relevant for salt stress response uniquely present in leaves of *B. halophila*.

Compared to *B. halophila*, only a few genes were identified from the GRNs of *B. platyphylla* (Fig. 5c, Table 5 and S6). Based on the homologous information of *Arabidopsis* genes associated with salt stress response, signaling, and tolerance reported in public articles (Table 3), we found a gene (*Bpev01.c1966.g0003*) encoding UDP-d-xylose 4-epimerase 1/MURUS4 (UXE1/MUR4) from the GRN of BPLeaf and a gene (*Bpev01.c0022.g0007*) encoding Histone H1-3 (HIS1-3) from the GRN of BPRoot. These results indicated that the genes identified from our GRNs were possibly capable of salt stress regulation in *B. platyphylla*.

**Table 5.** One inferred GRN relevant for salt stress response uniquely present in leaves of *B. platyphylla*.

**Table 6.** One inferred GRN relevant for salt stress response uniquely present in roots of *B. platyphylla*.

### Prediction of shared genetic regulatory mechanisms controlling salt stress response in two birch species

We studied the expression patterns of 121 shared DEGs in two birch species (Fig. 6a). Of which, compared to 0 h, 59 up-regulated genes and 62 down-regulated genes were detected in BHLeaf, 69 up-regulated genes and 52 down-regulated genes in BPLeaf, and 76 up-regulated genes and 45 down-regulated genes in BPRoot. Moreover, in any tissues of two birch species, 40 DEGs had decreased expression levels, 25 DEGs showed increased or decreased expression levels, and 56 DEGs exhibited increased expression levels.

**Fig. 6.**
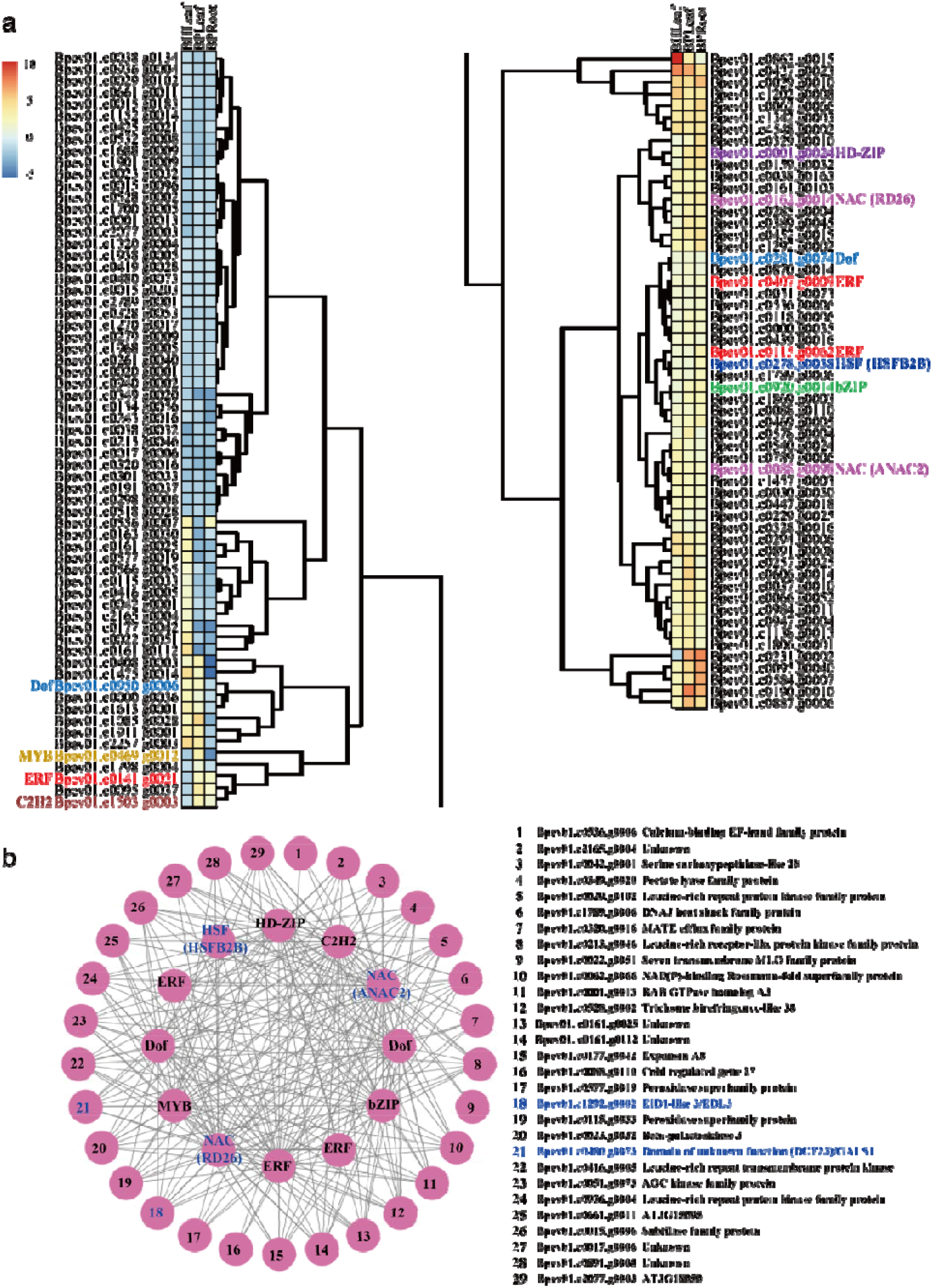
Analysis of shared DEGs between *B. halophila* and *B. platyphylla*. a, Expression patterns of shared DEGs among BHLeaf, BPLeaf, and BPRoot. HD-ZIP, Homeodomain-leucine zipper transcription factor; NAC, Petunia no apical meristem and ATAF1, ATAF2, and CUC2; ERF, Ethylene response factor; bZIP, Basic region/leucine zipper motif transcription factor; Dof, DNA binding with one finger transcription factor; MYB, Myeloblastosis; C2H2, Cys2His2-type zinc-finger transcription factor; HSF, Heat shock transcription factor. b, Inferred GRN involved in salt stress response commonly present in *B. halophila* and *B. platyphylla*. Genes homologous to *Arabidopsis* genes related to salt stress signaling and tolerance were marked in blue.

Further, based on the expression profiles of shared DEGs under the parameter of probability threshold above 0.01, we inferred a common GRN related to salt stress response between two birch species (Fig. 6b, Table 7). Analysis of this shared GRN showed that three TFs, including *Bpev01.c0162.g0014*, *Bpev01.c0088.g0098*, *and Bpev01.c0278.g0038*, were closely homologous to *responsive to desiccation 26* (*RD26*), *Arabidopsis NAC domain containing protein 2* (*ANAC2*), and *heat shock transcription factor B2B* (*HSFB2B*). Additionally, six genes, including *Bpev01.c1292.g0002*, *Bpev01.c0480.g0073*, *Bpev01.c1938.g0005*, *Bpev01.c1798.g0004*, *Bpev01.c0294.g0006*, and *Bpev01.c0452.g0015*, were respectively highly homologous to *EID1-like 3* (*EDL3*), *domain of unknown function* (*DUF23*) (*GALS1*), *K^+^ efflux antiporter 6* (*KEA6*), *mitogen-activated protein (MAP) kinase kinase 9* (*MKK9*), *nine-cis-epoxy-carotenoid dioxygenase 3* (*NCED3*), and *plant U-box 19* (*PUB19*). Accordingly, we suggested that these nine genes might play a critical role in response to salt stress in birch seedlings.

**Table 7.** One inferred GRN relevant for salt stress response shared in different tissues of *B. halophila* and *B. platyphylla*.

### The expression profiles of candidate genes related to salt stress response in birch

To further examine the accuracy of these constructed GRNs responsible for the regulation of salt stress response, we randomly selected nine genes and performed the RT-qPCR assay (Fig. 7). In our study, contrary to 0 h, the expression of *Bpev01.c0162.g0014* (*RD26*), *Bpev01.c0088.g0098* (*ANAC2*), *Bpev01.c0278.g0038* (*HSFB2B*), and *Bpev01.c1292.g0002* (*EDL3*) was activated in roots and leaves treated by 200 mM NaCl for 24 h. Compared to 0 h, the expression of *Bpev01.c0480.g0073* (*GALS1*) and *Bpev01.c1938.g0005* (*KEA6*) was inhibited in roots and leaves subjected to 200 mM NaCl for 24 h. After salt treatment for 24 h, *Bpev01.c1798.g0004* (*MKK9*), *Bpev01.c1966.g0003* (*UXE1/MUR4*), and *Bpev01.c0022.g0007* (*HIS1-3*) were weakly expressed in roots and strongly expressed in leaves, as compared to 0 h. Accordingly, we suggested that the genes identified from our GRNs were definitely induced by 200 mM NaCl solutions, involving the involvement of these genes in affecting salt stress response in birch.

**Fig. 7.**
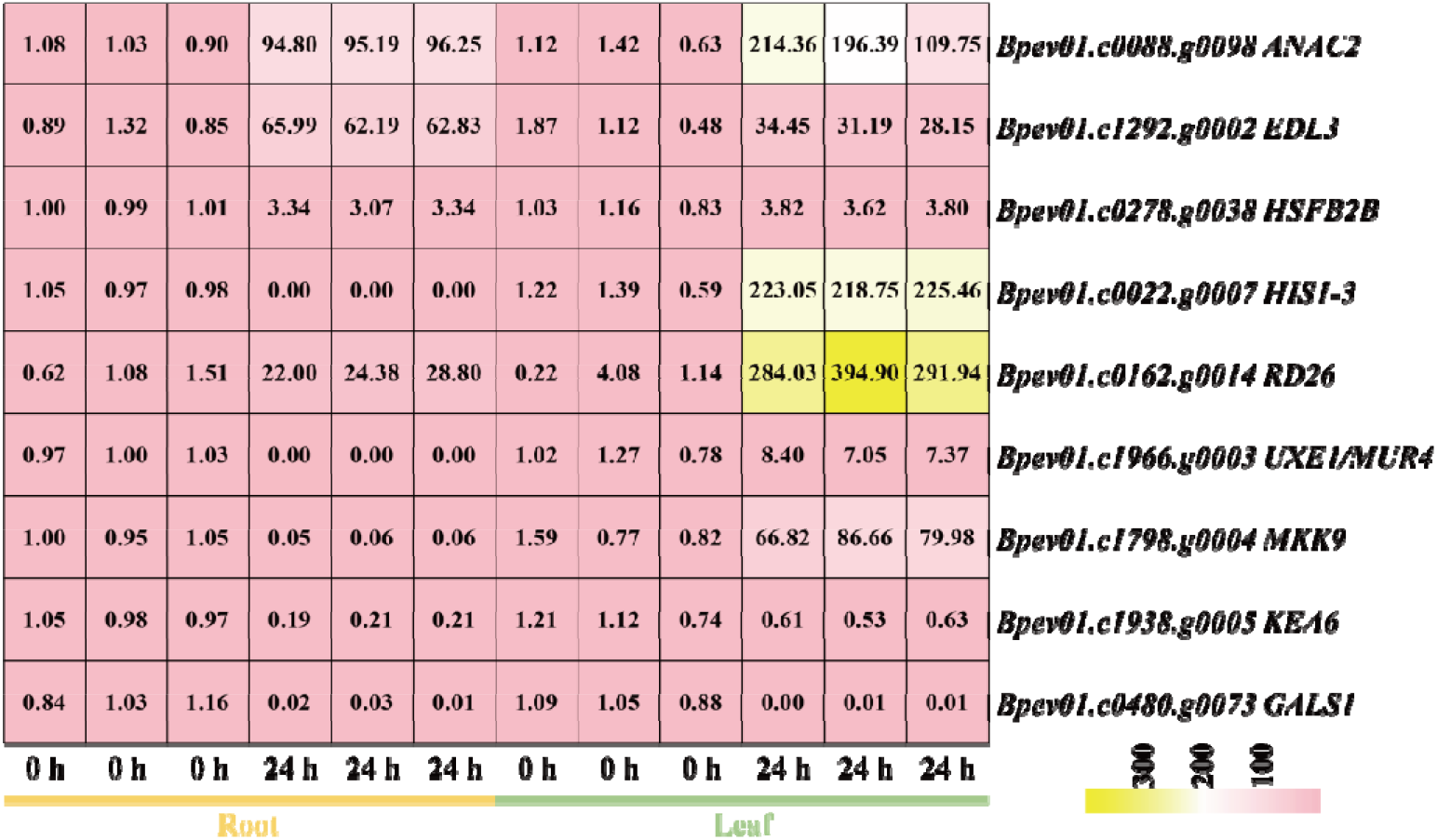
Expression profiles of candidate genes related to salt stress response in birch using RT-qPCR assay.

## Discussion

Salinity is one of the rising problems causing plant yield losses that happens all over the world, especially in arid and semiarid regions (Hassani et al. 2021). To deal with salt damage, it is definitely urgent for us to create salinity-resistant plant varieties through effective genetic engineering strategies. Thus, screening for salt-tolerant gene alleles from halophytes and transforming them into glycophytes grown well in saline soils are of great significance (Yuan et al. 2020). Birch is a pioneer hardwood tree species with ecological, economic, and evolutionary importance due to the presence of strong adaptation to environmental stimuli, outstanding phenotypic characteristics, wood properties, and medicinal uses (Chen et al. 2021a). Understanding the interplay between environmental stimuli and genetic factors in birch plants can allow us to accelerate the genetic improvement of forestry species strongly adapted to salt-affected areas.

A series of physiological and molecular mechanisms can be triggered when plants are exposed to high saline solutions (Ding et al. 2024). Unraveling these mechanisms and identifying critical genes involved in salt response are crucial for developing salt-tolerant varieties (Xiao and Zhou 2023). In this study, the number of DEGs discovered in leaves of *B. halophila* showed a decrease as compared to leaves and roots of *B. platyphylla*. Due to the fact that halophytes are naturally grown in high saline soils, halophytes probably have stronger adaptability to salt stress than glycophytes (Chen and Wang 2024). Once the halophytes are supplemented by high concentrations of NaCl solutions, halophytes enable themselves to maintain the intracellular steady-state level in vivo regardless of dramatic transcriptional changes (Barros et al. 2021). These specific coping strategies to survive salt stress might be the consequence of halophytes and glycophytes having been adapted to distinct natural habitats over a long period of evolution (Acosta-Motos et al. 2017). Meanwhile, the number of down-regulated genes was more than that of up-regulated genes in *B. halophila*, which was probably attributed to halophytes’ ability to homeostasis maintenance at transcriptional levels (Barros et al. 2021). The increased expression folds of genes found in glycophytes as a result of salt stress may be more flexible for them to respond to salt stress than halophytes (İbrahimova et al. 2021).

Salt stress is a major threat to plant growth and developmental activities, which compels plants to initiate multiple salt responsive strategies to promote their survival rates under high saline conditions (Xiao and Zhou 2023; Goudarzi et al. 2024). In our study, the expression profiles of DEGs detected in leaves of *B. halophila* showed a high similarity to that of roots in *B. platyphylla*, which might be explained by the possibility of their needs to adopt similar responsive mechanisms to overcome salt damage in two birch species. For instance, both of the halophytes’ leaves and glycophytes’ roots take actions to up-regulate salt stress-related genes to reduce intracellular osmotic potential and maintain water balance, further driving the force to synthesize and accumulate osmoregulatory substances (Cheeseman 2015). Beside, salt stress can disrupt ion homeostasis within plant cells. In order to maintain ion homeostasis, genes related to ion channels and transporters in the leaves of halophytes and the roots of glycophytes can be activated (Bian et al. 2025) Moreover, salt stress can induce a large amounts of ROS accumulation through oxidative stresses (Lodeyro and Carrillo 2015). To address this challenge, the leaves of halophytes and the roots of glycophytes enhanced the activity of antioxidant enzymes and activated the expression of related genes to remove ROS (Rahman et al. 2021).

Cell wall is a dynamic and complex structure that surrounds every cell of a plant, which provides both physical support and a means of cell-to-cell communication (Zhang et al. 2021a). The composition and structure of cell walls should be tightly regulated to ensure dynamic changes upon encountering environmental stimuli or to meet the needs of the rapidly growing cells (Shao et al. 2018a; Zhang et al. 2021b). High salinity can disrupt cell wall integrity, normal signaling, and nutritional functions (Dabravolski and Isayenkov 2023). In the present study, some DEGs were significantly enriched in cell wall organization or biogenesis in two birch species. Such results might support the viewpoint that plant cell wall is the first line of defense against salt stimulus (Zhang et al. 2021a) and its status is constantly monitored to optimize plant growth and development through influencing a series of stress response mechanisms (Rui and Dinneny 2020). Moreover, we respectively found many genes related to cell wall biosynthesis and encoding cell wall-localized glycoproteins down-regulated in *B. halophila* and up-regulated in *B. platyphylla*, suggesting the occurrence of a decreased cell wall biosynthesis in *B. halophila* under salt stress. Previous study indicates that a decreased cell wall biosynthesis may be a way for plants to adjust themselves to salt adaptation (Vaahtera et al. 2019; Ahmad et al. 2023). The defective cell wall biosynthesis makes plants sensitive to salt stress (Zhang et al. 2020; Dabravolski and Isayenkov 2023), which was opposite to our findings of leaves of *B. halophila* with the supplement of 200 mM NaCl solutions. It is assumed that halophytes and glycophytes probably have developed independent molecular mechanisms to alleviate or eliminate the negative effects of salt stress during the long-term evolutionary process.

Accumulating evidences indicate that phytohormones not only control plant growth, development, and productivity under normal conditions but also mediate the sophisticated crosstalk between salinity signals and plant growth adaptation (Yu et al. 2020). Unlike *B. halophila*, we found some genes relevant for hormone biosynthesis and signaling displaying altered expressions in *B. platyphylla*, which was likely to imply the presence of postponed hormonal biosynthesis and signaling pathways in leaves of *B. halophila* supplemented by high saline solutions. Generally speaking, plant hormone-related pathways activated at late stage are beneficial for plants to assess the degree of salt tolerance and optimize resource allocation (Verma et al. 2016). Halophytes can alleviate the detrimental effects caused by salt exposure or re-schedule salt-sensitive developmental stages by the activation of hormonal pathways at late stage (Zhao et al. 2020a). These findings indicated the abnormal hormone biosynthesis and signaling could confer seedlings of *B. halophila* to have strong adaptability and plasticity in response to salt stress.

It is well documented that a wide range of genetic drivers governing salt stress signaling and tolerance have been extensively studied in woody plants (Zhang et al. 2021b), we employed a random forests algorithm to construct several GRNs responsible for decoding the molecular mechanisms underpinning salt stress response of two birch species. In this shared GRN between two birch species, we found nine genes homologous to *Arabidopsis* genes associated with salt stress response, signaling, and tolerance (Table 3), including *RD26*, *ANAC2*, *HSFB2B*, *EDL3*, *GALS1*, *KEA6*, *MKK9*, *NCED3*, and *PUB19*. Researchers have demonstrated that SRM1 (*Salt-Related MYB1*) is a MYB-like transcription factor and plays a negative role in seed germination under saline conditions by regulating the expression levels of *RD26* in *Arabidopsis* (Wang et al. 2015). *ANAC2* is well known as its function in incorporating the salinity stimuli into plant lateral root developmental processes via ethylene and auxin signaling pathways (He et al. 2005). In *Arabidopsis*, *HSFB2B* represses a morning clock gene *PSEUDO-RESPONSE REGULATOR 7* (*PRR7*), which functions as a key factor in sustaining the circadian rhythm under high saline conditions (Kolmos et al. 2014). *EDL3* is presumed to act as a component of an ubiquitin ligase complex that specifically directs negatively acting factors in ABA signaling to degradation via the proteasome, thereby integrating hormone signal transduction towards salt injury (Koops et al. 2011). Galactan synthesized by GALS1 can form the side chains of pectic rhamnogalacturonan I. The excessive accumulation of galactan contributes to the improvement of *Arabidopsis* plants’ salt tolerance (Yan et al. 2023). In *Arabidopsis*, *histone deacetylase 15* (*HDA15*) over-expressed transgenic plants exhibit strong salt tolerance by activating an ABA biosynthetic gene *NCED3* (Truong et al. 2021). Under salt stress, one plant U-box armadillo repeat protein, *PUB19*, can work together with *PUB18* to inhibit seed germination in *Arabidopsis* (Bergler and Hoth 2011). Expression of active *MKK9* protein enhances the sensitivity of transgenic seedlings to salt stress, whereas loss of *MKK9* activity enhances salt tolerance (Xu et al. 2008), indicating a role for *MKK9* in mediating salt stress response (Xu et al. 2008; Shen et al. 2019; Liu et al. 2020). *KEA6*, belongs to *KEA* family, works coordinately with KEA4, KEA5, *Na^+^/H^+^ exchanger 5* (*NHX5*), and *NHX6* to facilitate endosomal pH homeostasis and salt tolerance (Zhu et al. 2018). In the present study, we randomly selected several genes and performed the RT-qPCR assay to study their expression profiles under salt stress for 0 and 24 h. According to the RT-qPCR results, *RD26*, *ANAC2*, *HSFB2B*, *EDL3*, and *GALS1* could be activated or inhibited by 200 mM NaCl solutions in *B. platyphylla*. We therefore proposed this shared GRN relevant for unraveling how birch plants reacted to salt stress. Although ecological, physiological, and genetic variations occurred between halophytes and glycophytes, they appeared to share a set of genetic elements to alleviate and endure salt-induced phytotoxicity in birch.

Under salt stress, halophytes can activate the expression levels of a large number of genes to maintain the cellular and ionic homeostasis, which is inconsistent with glycophytes (Kumar et al. 2024). In the GRN of *B. halophila*, seven genes were closely homologously to *Arabidopsis* genes associated with salt stress response, signaling, and tolerance (Table 3), namely *SnRK2.6/OST1*, *HAK5*, *NDL1*, *ANN4*, *ARR12*, *CIPK21*, and *LRX4*. Given the importance of these genes mentioned above in salt stress control in *Arabidopsis* (Pandey et al. 2015; Kawa et al. 2020; Zhao et al. 2020a; Gupta et al. 2021; Yan et al. 2021; Liu et al. 2023), it was reasonably thought that our GRN was reliable for addressing the molecular mechanisms underlying salt stress response in *B. halophila*. Moreover, analysis of the GRNs constructed in *B. platyphylla* showed that two genes coding for *UXE1*/*MUR4* and *HIS1-3*. Under high salt conditions, *UXE1*/*MUR4* has a positive influence on UDP-arabinopyranose biosynthesis during root development (Zhao et al. 2019). The disruption of one linker histone variant *HIS1-3* can enhance plants’ salt tolerance, leading to the benefits of regulating the transcriptional levels of SOS (salt overly sensitive) genes (Wu et al. 2022). RT-qPCR results showed that these two genes identified from our GRNs could be induced by 200 mM NaCl solutions in *B. platyphylla*. We therefore suggested that our GRNs in *B. platyphylla* could make a contribution to explaining the distinctive mechanisms of salt stress response from *B. halophila*. These results further confirmed that during the long-term evolution, halophytes have built diverse regulatory mechanisms to accommodate saline environments by orchestrating the synthesis, signaling, and metabolism of various substances via multiple pathways (Volkov and Amtmann 2006; Cheeseman 2015; Zhao et al. 2021; Rahman et al. 2022; Wang et al. 2023), which exhibited the specific genetic features present in a halophytic species in comparison with a closely related glycophytic species.

## Conclusions

One important feature of halophytes is endowed with special morphological, anatomical, and physiological characteristics as well as genetic regulatory networks to well grow and reproduce in high saline soils. Accordingly, halophytes can be selected as a desirable model to uncover the molecular mechanisms underlying salinity tolerance. In our study, the expression patterns of genes in leaves of *B. halophila* showed a similarity to roots of *B. platyphylla*. Genes significantly enriched in cell wall organization or biogenesis were detected in leaves of *B. halophila* and roots of *B. platyphylla*. Genes related to hormone biosynthesis and signaling were expressed at differential levels in *B. platyphylla*. Moreover, we respectively revealed the potential molecular mechanisms of salt stress response commonly and uniquely occurring in two species. Our results could explain the salt-resistant variations at the molecular level between *B. halophila* and *B. platyphylla*, which can provide insights into the genetic control of birch response to salt stress and offer some informative clues to the genetic improvement of glycophytes’ salt tolerance.

## Supporting information

Table 1 Primer sequences used for RT-qPCR assay.

Table 2 GO enrichment analysis of genes in each cluster divided by STEM software.

Table 3 Genes related to salt stress response differentially expressed in B. halophila and B. platyphylla.

Table 4 One inferred GRN relevant for salt stress response uniquely present in leaves of B. halophila.

Table 5 One inferred GRN relevant for salt stress response uniquely present in leaves of B. platyphylla.

Table 6 One inferred GRN relevant for salt stress response uniquely present in roots of B. platyphylla.

Table 7 One inferred GRN relevant for salt stress response shared in different tissues of B. halophila and B. platyphylla.

Quantitative related information

## Abbreviations

BHLeaf: leaves of *B. halophila* treated with 200 mM NaCl for 24 h
BPLeaf: leaves of *B. platyphylla* treated with 200 mM NaCl for 24 h
BPRoot: roots of *B. platyphylla* treated with 200 mM NaCl for 24 h
DEG: differentially expressed gene
FDR: false discovery rate
FPKM: fragments per kilobase per million mapped reads
GO: Gene Ontology
GRN: gene regulatory network
NAA: naphthalene acetic acid
ROS: reactive oxygen species
RT-qPCR: reverse transcription quantitative PCR
SRA: Sequence Read Archive
STEM: Short Time-series Expression Miner
TF: transcription factor
WPM: woody plant medium.

## Acknowledgements

The authors thank the investigators who generated and made publicly available the transcriptome datasets used in this study, as well as the developers of the open-source bioinformatics tools used for data analysis. The authors also thank their institutions for providing research facilities and support. No specific funding was received for this study.

## Author contributions

Zhi LI: Conceptualization, Data curation, Formal analysis, Investigation, Methodology, Software, Visualization, Writing—original draft. Qingjing HUANG: Data curation, Formal analysis, Investigation, Validation. Liangju LI: Data curation, Formal analysis, Investigation, Validation. Lixiu YAN: Conceptualization, Project administration, Resources, Supervision, Writing—review and editing. Bangmeng ZHOU: Data curation, Formal analysis, Software, Validation, Visualization. Jie CHENG: Conceptualization, Project administration, Resources, Supervision, Writing—review and editing. All authors read and approved the final manuscript.

## Data and materials availability

The datasets generated and analyzed during the present study are available from the corresponding author on reasonable request.

## Declaration of ethical and legal standards compliance

Not applicable.

## Tables 1-7 in supplementary materials

