## Supplementary material for "Transcriptome analysis provides insights into candidate genetic regulators of birch response to salt stress": Table 1 Primer sequences used for RT-qPCR assay.

| **Table S1 Primer sequences used for RT-qPCR assay.** | |
| --- | --- |
| **Gene ID (Gene name)** | **Primer sequences (5'~3')** |
| ***Bpev01.c0162.g0014 (RD26)*** | **F: GTAAGCCTCAACTCGGTGC** |
|  | **R: CGAGTTATCGATCCGCTGAG** |
| ***Bpev01.c0088.g0098 (ANAC2)*** | **F: CGTGTACTTCGACACGTCGG** |
|  | **R: GGGCTTCTGCATGTACATGAAC** |
| ***Bpev01.c0278.g0038 (HSFB2B)*** | **F: GGTGGCGGTGCACCGG** |
|  | **R: GTACAGAAGCTCCGGTACCG** |
| ***Bpev01.c1292.g0002 (EDL3)*** | **F: GACGGCAGGTAGCGCTGG** |
|  | **R: CAATCACGCCGACCCCACC** |
| ***Bpev01.c0480.g0073 (GALS1)*** | **F: CGCCGCCGCTCTGTTCGT** |
|  | **R: CAGGGAAGGTGCAGTTGAC** |
| ***Bpev01.c1938.g0005 (KEA6)*** | **F: GATCTCCGTCGTCGGAGC** |
|  | **R: CACTGTTATTGAAGCCTCCAGC** |
| ***Bpev01.c1798.g0004 (MKK9)*** | **F: GGCAGTTGTTCGAGAACGG** |
|  | **R: CATAGAAAGCGCTGGTGCGC** |
| ***Bpev01.c1966.g0003 (UXE1/MUR4)*** | **F: GCTAAATTTTGGCAGGTCCAG** |
|  | **R: CCAATATAGCCAGCACCTCCTG** |
| ***Bpev01.c0022.g0007 (Histone H1-3)*** | **F: CGTACAAGCTCTCCCTGGTG** |
|  | **R: CGTACTCGCCTTCCTCTTCG** |
| ***FJ228477.1 (18S rRNA)*** | **F: ATCTTGGGTTGGGCAGATCG** |
|  | **R: CATTACTCCGATCCCGAAGG** |
