## Supplementary material for "Transcriptome analysis provides insights into candidate genetic regulators of birch response to salt stress": Table 2 GO enrichment analysis of genes in each cluster divided by STEM software.

| **Table S2 GO enrichment analysis of genes in the clusters divided by STEM software.** | | | | | |
| --- | --- | --- | --- | --- | --- |
| **GO ID** | **Description** | **q-value** | **Gene ID** | **Count** | **Cluster ID** |
| **GO:0009834** | **plant-type secondary cell wall biogenesis** | **2.58749E-13** | **Bpev01.c0000.g0006/Bpev01.c0008.g0004/Bpev01.c0015.g0038/Bpev01.c0015.g0046/Bpev01.c0022.g0005/Bpev01.c0052.g0019/Bpev01.c0088.g0120/Bpev01.c0115.g0065/Bpev01.c0121.g0011/Bpev01.c0161.g0013/Bpev01.c0166.g0009/Bpev01.c0167.g0004/Bpev01.c0169.g0033/Bpev01.c0189.g0059/Bpev01.c0190.g0082/Bpev01.c0275.g0062/Bpev01.c0288.g0009/Bpev01.c0344.g0013/Bpev01.c0374.g0017/Bpev01.c0383.g0004/Bpev01.c0483.g0013/Bpev01.c0603.g0003/Bpev01.c0817.g0012/Bpev01.c1610.g0004/Bpev01.c2001.g0008** | **25** | **Cluster 3** |
| **GO:0042546** | **cell wall biogenesis** | **6.53444E-12** | **Bpev01.c0000.g0006/Bpev01.c0008.g0004/Bpev01.c0015.g0038/Bpev01.c0015.g0046/Bpev01.c0018.g0088/Bpev01.c0022.g0005/Bpev01.c0029.g0110/Bpev01.c0052.g0019/Bpev01.c0088.g0120/Bpev01.c0115.g0065/Bpev01.c0121.g0011/Bpev01.c0154.g0062/Bpev01.c0161.g0013/Bpev01.c0163.g0015/Bpev01.c0166.g0009/Bpev01.c0167.g0004/Bpev01.c0169.g0033/Bpev01.c0189.g0059/Bpev01.c0190.g0014/Bpev01.c0190.g0082/Bpev01.c0265.g0024/Bpev01.c0275.g0062/Bpev01.c0288.g0009/Bpev01.c0344.g0013/Bpev01.c0374.g0017/Bpev01.c0383.g0004/Bpev01.c0483.g0013/Bpev01.c0598.g0015/Bpev01.c0603.g0003/Bpev01.c0621.g0009/Bpev01.c0781.g0019/Bpev01.c0817.g0012/Bpev01.c0892.g0012/Bpev01.c0892.g0015/Bpev01.c0892.g0016/Bpev01.c0901.g0009/Bpev01.c1082.g0001/Bpev01.c1610.g0004/Bpev01.c1659.g0003/Bpev01.c2001.g0008/Bpev01.c2157.g0002** | **41** |  |
| **GO:0071554** | **cell wall organization or biogenesis** | **6.53444E-12** | **Bpev01.c0000.g0006/Bpev01.c0008.g0004/Bpev01.c0015.g0038/Bpev01.c0015.g0046/Bpev01.c0015.g0262/Bpev01.c0018.g0088/Bpev01.c0018.g0093/Bpev01.c0022.g0005/Bpev01.c0029.g0110/Bpev01.c0044.g0114/Bpev01.c0051.g0197/Bpev01.c0051.g0198/Bpev01.c0052.g0019/Bpev01.c0088.g0024/Bpev01.c0088.g0120/Bpev01.c0094.g0029/Bpev01.c0115.g0065/Bpev01.c0121.g0011/Bpev01.c0154.g0062/Bpev01.c0161.g0013/Bpev01.c0163.g0015/Bpev01.c0166.g0009/Bpev01.c0166.g0033/Bpev01.c0167.g0004/Bpev01.c0169.g0024/Bpev01.c0169.g0033/Bpev01.c0189.g0059/Bpev01.c0190.g0014/Bpev01.c0190.g0082/Bpev01.c0214.g0029/Bpev01.c0253.g0016/Bpev01.c0265.g0024/Bpev01.c0275.g0062/Bpev01.c0288.g0009/Bpev01.c0332.g0002/Bpev01.c0344.g0013/Bpev01.c0374.g0017/Bpev01.c0383.g0004/Bpev01.c0420.g0022/Bpev01.c0483.g0013/Bpev01.c0502.g0008/Bpev01.c0598.g0015/Bpev01.c0603.g0003/Bpev01.c0614.g0012/Bpev01.c0621.g0009/Bpev01.c0639.g0023/Bpev01.c0727.g0003/Bpev01.c0739.g0007/Bpev01.c0781.g0019/Bpev01.c0817.g0012/Bpev01.c0845.g0002/Bpev01.c0848.g0015/Bpev01.c0892.g0012/Bpev01.c0892.g0015/Bpev01.c0892.g0016/Bpev01.c0901.g0009/Bpev01.c0902.g0015/Bpev01.c0920.g0001/Bpev01.c1082.g0001/Bpev01.c1183.g0022/Bpev01.c1484.g0017/Bpev01.c1596.g0001/Bpev01.c1610.g0004/Bpev01.c1659.g0003/Bpev01.c2001.g0008/Bpev01.c2001.g0009/Bpev01.c2157.g0002** | **67** |  |
| **GO:0071669** | **plant-type cell wall organization or biogenesis** | **7.19274E-12** | **Bpev01.c0000.g0006/Bpev01.c0008.g0004/Bpev01.c0015.g0038/Bpev01.c0015.g0046/Bpev01.c0018.g0088/Bpev01.c0022.g0005/Bpev01.c0029.g0110/Bpev01.c0052.g0019/Bpev01.c0088.g0120/Bpev01.c0115.g0065/Bpev01.c0121.g0011/Bpev01.c0161.g0013/Bpev01.c0163.g0015/Bpev01.c0166.g0009/Bpev01.c0166.g0033/Bpev01.c0167.g0004/Bpev01.c0169.g0033/Bpev01.c0189.g0059/Bpev01.c0190.g0014/Bpev01.c0190.g0082/Bpev01.c0265.g0024/Bpev01.c0275.g0062/Bpev01.c0288.g0009/Bpev01.c0332.g0002/Bpev01.c0344.g0013/Bpev01.c0374.g0017/Bpev01.c0383.g0004/Bpev01.c0483.g0013/Bpev01.c0598.g0015/Bpev01.c0603.g0003/Bpev01.c0639.g0023/Bpev01.c0727.g0003/Bpev01.c0817.g0012/Bpev01.c0892.g0012/Bpev01.c0920.g0001/Bpev01.c1484.g0017/Bpev01.c1596.g0001/Bpev01.c1610.g0004/Bpev01.c2001.g0008/Bpev01.c2157.g0002** | **40** |  |
| **GO:0009832** | **plant-type cell wall biogenesis** | **1.80544E-11** | **Bpev01.c0000.g0006/Bpev01.c0008.g0004/Bpev01.c0015.g0038/Bpev01.c0015.g0046/Bpev01.c0018.g0088/Bpev01.c0022.g0005/Bpev01.c0029.g0110/Bpev01.c0052.g0019/Bpev01.c0088.g0120/Bpev01.c0115.g0065/Bpev01.c0121.g0011/Bpev01.c0161.g0013/Bpev01.c0163.g0015/Bpev01.c0166.g0009/Bpev01.c0167.g0004/Bpev01.c0169.g0033/Bpev01.c0189.g0059/Bpev01.c0190.g0082/Bpev01.c0265.g0024/Bpev01.c0275.g0062/Bpev01.c0288.g0009/Bpev01.c0344.g0013/Bpev01.c0374.g0017/Bpev01.c0383.g0004/Bpev01.c0483.g0013/Bpev01.c0598.g0015/Bpev01.c0603.g0003/Bpev01.c0817.g0012/Bpev01.c0892.g0012/Bpev01.c1610.g0004/Bpev01.c2001.g0008/Bpev01.c2157.g0002** | **32** |  |
| **GO:0000271** | **polysaccharide biosynthetic process** | **9.75722E-10** | **Bpev01.c0000.g0006/Bpev01.c0015.g0038/Bpev01.c0015.g0046/Bpev01.c0015.g0150/Bpev01.c0015.g0262/Bpev01.c0018.g0088/Bpev01.c0029.g0110/Bpev01.c0052.g0019/Bpev01.c0088.g0024/Bpev01.c0088.g0120/Bpev01.c0151.g0003/Bpev01.c0154.g0062/Bpev01.c0163.g0015/Bpev01.c0166.g0009/Bpev01.c0166.g0033/Bpev01.c0189.g0059/Bpev01.c0190.g0014/Bpev01.c0253.g0016/Bpev01.c0265.g0024/Bpev01.c0275.g0062/Bpev01.c0288.g0009/Bpev01.c0344.g0013/Bpev01.c0374.g0017/Bpev01.c0430.g0012/Bpev01.c0502.g0008/Bpev01.c0522.g0040/Bpev01.c0582.g0012/Bpev01.c0598.g0015/Bpev01.c0603.g0003/Bpev01.c0621.g0009/Bpev01.c0679.g0017/Bpev01.c0781.g0019/Bpev01.c0845.g0002/Bpev01.c0848.g0015/Bpev01.c1082.g0001/Bpev01.c2001.g0008/Bpev01.c2157.g0002** | **37** |  |
| **GO:0010383** | **cell wall polysaccharide metabolic process** | **9.25782E-09** | **Bpev01.c0015.g0038/Bpev01.c0015.g0046/Bpev01.c0029.g0110/Bpev01.c0052.g0019/Bpev01.c0088.g0120/Bpev01.c0115.g0065/Bpev01.c0154.g0062/Bpev01.c0166.g0009/Bpev01.c0166.g0033/Bpev01.c0189.g0059/Bpev01.c0190.g0014/Bpev01.c0265.g0024/Bpev01.c0275.g0062/Bpev01.c0288.g0009/Bpev01.c0344.g0013/Bpev01.c0603.g0003/Bpev01.c0621.g0009/Bpev01.c0739.g0007/Bpev01.c0781.g0019/Bpev01.c0892.g0012/Bpev01.c0892.g0015/Bpev01.c0892.g0016/Bpev01.c0901.g0009/Bpev01.c1082.g0001/Bpev01.c1659.g0003/Bpev01.c2001.g0008/Bpev01.c2157.g0002** | **27** |  |
| **GO:0005976** | **polysaccharide metabolic process** | **1.16838E-08** | **Bpev01.c0000.g0006/Bpev01.c0015.g0038/Bpev01.c0015.g0046/Bpev01.c0015.g0150/Bpev01.c0015.g0262/Bpev01.c0016.g0180/Bpev01.c0018.g0088/Bpev01.c0027.g0103/Bpev01.c0029.g0110/Bpev01.c0044.g0114/Bpev01.c0051.g0197/Bpev01.c0051.g0198/Bpev01.c0052.g0019/Bpev01.c0088.g0024/Bpev01.c0088.g0120/Bpev01.c0115.g0065/Bpev01.c0151.g0003/Bpev01.c0154.g0062/Bpev01.c0163.g0015/Bpev01.c0166.g0009/Bpev01.c0166.g0033/Bpev01.c0189.g0059/Bpev01.c0190.g0014/Bpev01.c0214.g0029/Bpev01.c0253.g0016/Bpev01.c0265.g0024/Bpev01.c0275.g0062/Bpev01.c0288.g0009/Bpev01.c0332.g0002/Bpev01.c0344.g0013/Bpev01.c0374.g0017/Bpev01.c0430.g0012/Bpev01.c0502.g0008/Bpev01.c0522.g0023/Bpev01.c0522.g0024/Bpev01.c0522.g0040/Bpev01.c0582.g0012/Bpev01.c0598.g0015/Bpev01.c0603.g0003/Bpev01.c0621.g0009/Bpev01.c0679.g0017/Bpev01.c0739.g0007/Bpev01.c0781.g0019/Bpev01.c0845.g0002/Bpev01.c0848.g0015/Bpev01.c0892.g0012/Bpev01.c0892.g0015/Bpev01.c0892.g0016/Bpev01.c0901.g0009/Bpev01.c1082.g0001/Bpev01.c1659.g0003/Bpev01.c2001.g0008/Bpev01.c2001.g0009/Bpev01.c2157.g0002** | **54** |  |
| **GO:0044264** | **cellular polysaccharide metabolic process** | **1.86367E-08** | **Bpev01.c0000.g0006/Bpev01.c0015.g0038/Bpev01.c0015.g0046/Bpev01.c0016.g0180/Bpev01.c0029.g0110/Bpev01.c0052.g0019/Bpev01.c0088.g0120/Bpev01.c0115.g0065/Bpev01.c0151.g0003/Bpev01.c0154.g0062/Bpev01.c0166.g0009/Bpev01.c0166.g0033/Bpev01.c0189.g0059/Bpev01.c0190.g0014/Bpev01.c0214.g0029/Bpev01.c0253.g0016/Bpev01.c0265.g0024/Bpev01.c0275.g0062/Bpev01.c0288.g0009/Bpev01.c0332.g0002/Bpev01.c0344.g0013/Bpev01.c0374.g0017/Bpev01.c0522.g0023/Bpev01.c0522.g0024/Bpev01.c0522.g0040/Bpev01.c0582.g0012/Bpev01.c0598.g0015/Bpev01.c0603.g0003/Bpev01.c0621.g0009/Bpev01.c0679.g0017/Bpev01.c0739.g0007/Bpev01.c0781.g0019/Bpev01.c0845.g0002/Bpev01.c0892.g0012/Bpev01.c0892.g0015/Bpev01.c0892.g0016/Bpev01.c0901.g0009/Bpev01.c1082.g0001/Bpev01.c1659.g0003/Bpev01.c2001.g0008/Bpev01.c2157.g0002** | **41** |  |
| **GO:0071555** | **cell wall organization** | **2.2769E-08** | **Bpev01.c0000.g0006/Bpev01.c0015.g0046/Bpev01.c0015.g0262/Bpev01.c0018.g0088/Bpev01.c0018.g0093/Bpev01.c0022.g0005/Bpev01.c0029.g0110/Bpev01.c0044.g0114/Bpev01.c0051.g0197/Bpev01.c0051.g0198/Bpev01.c0052.g0019/Bpev01.c0088.g0024/Bpev01.c0088.g0120/Bpev01.c0094.g0029/Bpev01.c0161.g0013/Bpev01.c0163.g0015/Bpev01.c0166.g0009/Bpev01.c0166.g0033/Bpev01.c0169.g0024/Bpev01.c0189.g0059/Bpev01.c0190.g0014/Bpev01.c0214.g0029/Bpev01.c0253.g0016/Bpev01.c0265.g0024/Bpev01.c0288.g0009/Bpev01.c0332.g0002/Bpev01.c0374.g0017/Bpev01.c0502.g0008/Bpev01.c0598.g0015/Bpev01.c0603.g0003/Bpev01.c0614.g0012/Bpev01.c0621.g0009/Bpev01.c0639.g0023/Bpev01.c0727.g0003/Bpev01.c0781.g0019/Bpev01.c0845.g0002/Bpev01.c0848.g0015/Bpev01.c0892.g0012/Bpev01.c0892.g0015/Bpev01.c0892.g0016/Bpev01.c0901.g0009/Bpev01.c0902.g0015/Bpev01.c0920.g0001/Bpev01.c1183.g0022/Bpev01.c1484.g0017/Bpev01.c1596.g0001/Bpev01.c1659.g0003/Bpev01.c2001.g0008/Bpev01.c2001.g0009/Bpev01.c2157.g0002** | **50** |  |
| **GO:0045229** | **external encapsulating structure organization** | **1.38233E-07** | **Bpev01.c0000.g0006/Bpev01.c0015.g0046/Bpev01.c0015.g0262/Bpev01.c0018.g0088/Bpev01.c0018.g0093/Bpev01.c0022.g0005/Bpev01.c0029.g0110/Bpev01.c0044.g0114/Bpev01.c0051.g0197/Bpev01.c0051.g0198/Bpev01.c0052.g0019/Bpev01.c0088.g0024/Bpev01.c0088.g0120/Bpev01.c0094.g0029/Bpev01.c0161.g0013/Bpev01.c0163.g0015/Bpev01.c0166.g0009/Bpev01.c0166.g0033/Bpev01.c0169.g0024/Bpev01.c0189.g0059/Bpev01.c0190.g0014/Bpev01.c0214.g0029/Bpev01.c0253.g0016/Bpev01.c0265.g0024/Bpev01.c0288.g0009/Bpev01.c0332.g0002/Bpev01.c0374.g0017/Bpev01.c0502.g0008/Bpev01.c0598.g0015/Bpev01.c0603.g0003/Bpev01.c0614.g0012/Bpev01.c0621.g0009/Bpev01.c0639.g0023/Bpev01.c0727.g0003/Bpev01.c0781.g0019/Bpev01.c0845.g0002/Bpev01.c0848.g0015/Bpev01.c0892.g0012/Bpev01.c0892.g0015/Bpev01.c0892.g0016/Bpev01.c0901.g0009/Bpev01.c0902.g0015/Bpev01.c0920.g0001/Bpev01.c1183.g0022/Bpev01.c1484.g0017/Bpev01.c1596.g0001/Bpev01.c1659.g0003/Bpev01.c2001.g0008/Bpev01.c2001.g0009/Bpev01.c2157.g0002** | **50** |  |
| **GO:0010410** | **hemicellulose metabolic process** | **4.34483E-07** | **Bpev01.c0015.g0038/Bpev01.c0015.g0046/Bpev01.c0052.g0019/Bpev01.c0088.g0120/Bpev01.c0115.g0065/Bpev01.c0154.g0062/Bpev01.c0190.g0014/Bpev01.c0275.g0062/Bpev01.c0288.g0009/Bpev01.c0344.g0013/Bpev01.c0621.g0009/Bpev01.c0739.g0007/Bpev01.c0781.g0019/Bpev01.c0892.g0012/Bpev01.c0892.g0015/Bpev01.c0892.g0016/Bpev01.c0901.g0009/Bpev01.c1082.g0001/Bpev01.c1659.g0003/Bpev01.c2001.g0008** | **20** |  |
| **GO:0045491** | **xylan metabolic process** | **6.10078E-07** | **Bpev01.c0015.g0038/Bpev01.c0015.g0046/Bpev01.c0052.g0019/Bpev01.c0088.g0120/Bpev01.c0115.g0065/Bpev01.c0154.g0062/Bpev01.c0190.g0014/Bpev01.c0275.g0062/Bpev01.c0288.g0009/Bpev01.c0344.g0013/Bpev01.c0621.g0009/Bpev01.c0739.g0007/Bpev01.c0781.g0019/Bpev01.c2001.g0008** | **14** |  |
| **GO:0044036** | **cell wall macromolecule metabolic process** | **7.48087E-07** | **Bpev01.c0015.g0038/Bpev01.c0015.g0046/Bpev01.c0029.g0110/Bpev01.c0052.g0019/Bpev01.c0088.g0120/Bpev01.c0115.g0065/Bpev01.c0154.g0062/Bpev01.c0166.g0009/Bpev01.c0166.g0033/Bpev01.c0189.g0059/Bpev01.c0190.g0014/Bpev01.c0265.g0024/Bpev01.c0275.g0062/Bpev01.c0288.g0009/Bpev01.c0344.g0013/Bpev01.c0420.g0022/Bpev01.c0603.g0003/Bpev01.c0621.g0009/Bpev01.c0739.g0007/Bpev01.c0781.g0019/Bpev01.c0892.g0012/Bpev01.c0892.g0015/Bpev01.c0892.g0016/Bpev01.c0901.g0009/Bpev01.c1082.g0001/Bpev01.c1659.g0003/Bpev01.c2001.g0008/Bpev01.c2157.g0002** | **28** |  |
| **GO:0070592** | **cell wall polysaccharide biosynthetic process** | **9.21395E-07** | **Bpev01.c0015.g0038/Bpev01.c0015.g0046/Bpev01.c0029.g0110/Bpev01.c0052.g0019/Bpev01.c0088.g0120/Bpev01.c0154.g0062/Bpev01.c0166.g0009/Bpev01.c0189.g0059/Bpev01.c0190.g0014/Bpev01.c0265.g0024/Bpev01.c0275.g0062/Bpev01.c0288.g0009/Bpev01.c0344.g0013/Bpev01.c0621.g0009/Bpev01.c0781.g0019/Bpev01.c1082.g0001/Bpev01.c2001.g0008/Bpev01.c2157.g0002** | **18** |  |
| **GO:0045492** | **xylan biosynthetic process** | **9.36005E-07** | **Bpev01.c0015.g0038/Bpev01.c0015.g0046/Bpev01.c0052.g0019/Bpev01.c0088.g0120/Bpev01.c0154.g0062/Bpev01.c0190.g0014/Bpev01.c0275.g0062/Bpev01.c0288.g0009/Bpev01.c0344.g0013/Bpev01.c0621.g0009/Bpev01.c0781.g0019/Bpev01.c2001.g0008** | **12** |  |
| **GO:0033692** | **cellular polysaccharide biosynthetic process** | **1.09274E-06** | **Bpev01.c0000.g0006/Bpev01.c0015.g0038/Bpev01.c0015.g0046/Bpev01.c0029.g0110/Bpev01.c0052.g0019/Bpev01.c0088.g0120/Bpev01.c0151.g0003/Bpev01.c0154.g0062/Bpev01.c0166.g0009/Bpev01.c0189.g0059/Bpev01.c0190.g0014/Bpev01.c0253.g0016/Bpev01.c0265.g0024/Bpev01.c0275.g0062/Bpev01.c0288.g0009/Bpev01.c0344.g0013/Bpev01.c0374.g0017/Bpev01.c0522.g0040/Bpev01.c0582.g0012/Bpev01.c0598.g0015/Bpev01.c0603.g0003/Bpev01.c0621.g0009/Bpev01.c0679.g0017/Bpev01.c0781.g0019/Bpev01.c0845.g0002/Bpev01.c1082.g0001/Bpev01.c2001.g0008/Bpev01.c2157.g0002** | **28** |  |
| **GO:0016051** | **carbohydrate biosynthetic process** | **3.82195E-06** | **Bpev01.c0000.g0006/Bpev01.c0015.g0038/Bpev01.c0015.g0046/Bpev01.c0015.g0150/Bpev01.c0015.g0262/Bpev01.c0018.g0088/Bpev01.c0029.g0110/Bpev01.c0052.g0019/Bpev01.c0088.g0024/Bpev01.c0088.g0120/Bpev01.c0151.g0003/Bpev01.c0154.g0062/Bpev01.c0163.g0015/Bpev01.c0166.g0009/Bpev01.c0166.g0033/Bpev01.c0189.g0059/Bpev01.c0190.g0014/Bpev01.c0253.g0016/Bpev01.c0265.g0024/Bpev01.c0275.g0062/Bpev01.c0288.g0009/Bpev01.c0344.g0013/Bpev01.c0374.g0017/Bpev01.c0430.g0012/Bpev01.c0502.g0008/Bpev01.c0522.g0040/Bpev01.c0582.g0012/Bpev01.c0598.g0015/Bpev01.c0603.g0003/Bpev01.c0621.g0009/Bpev01.c0659.g0005/Bpev01.c0679.g0017/Bpev01.c0781.g0019/Bpev01.c0845.g0002/Bpev01.c0848.g0015/Bpev01.c1082.g0001/Bpev01.c2001.g0008/Bpev01.c2157.g0002/Bpev01.c2327.g0001** | **39** |  |
| **GO:0044038** | **cell wall macromolecule biosynthetic process** | **7.40407E-06** | **Bpev01.c0015.g0038/Bpev01.c0015.g0046/Bpev01.c0029.g0110/Bpev01.c0052.g0019/Bpev01.c0088.g0120/Bpev01.c0154.g0062/Bpev01.c0166.g0009/Bpev01.c0189.g0059/Bpev01.c0190.g0014/Bpev01.c0265.g0024/Bpev01.c0275.g0062/Bpev01.c0288.g0009/Bpev01.c0344.g0013/Bpev01.c0621.g0009/Bpev01.c0781.g0019/Bpev01.c1082.g0001/Bpev01.c2001.g0008/Bpev01.c2157.g0002** | **18** |  |
| **GO:0070589** | **cellular component macromolecule biosynthetic process** | **7.40407E-06** | **Bpev01.c0015.g0038/Bpev01.c0015.g0046/Bpev01.c0029.g0110/Bpev01.c0052.g0019/Bpev01.c0088.g0120/Bpev01.c0154.g0062/Bpev01.c0166.g0009/Bpev01.c0189.g0059/Bpev01.c0190.g0014/Bpev01.c0265.g0024/Bpev01.c0275.g0062/Bpev01.c0288.g0009/Bpev01.c0344.g0013/Bpev01.c0621.g0009/Bpev01.c0781.g0019/Bpev01.c1082.g0001/Bpev01.c2001.g0008/Bpev01.c2157.g0002** | **18** |  |
| **GO:0034637** | **cellular carbohydrate biosynthetic process** | **2.15297E-05** | **Bpev01.c0000.g0006/Bpev01.c0015.g0038/Bpev01.c0015.g0046/Bpev01.c0029.g0110/Bpev01.c0052.g0019/Bpev01.c0088.g0120/Bpev01.c0151.g0003/Bpev01.c0154.g0062/Bpev01.c0166.g0009/Bpev01.c0189.g0059/Bpev01.c0190.g0014/Bpev01.c0253.g0016/Bpev01.c0265.g0024/Bpev01.c0275.g0062/Bpev01.c0288.g0009/Bpev01.c0344.g0013/Bpev01.c0374.g0017/Bpev01.c0522.g0040/Bpev01.c0582.g0012/Bpev01.c0598.g0015/Bpev01.c0603.g0003/Bpev01.c0621.g0009/Bpev01.c0679.g0017/Bpev01.c0781.g0019/Bpev01.c0845.g0002/Bpev01.c1082.g0001/Bpev01.c2001.g0008/Bpev01.c2157.g0002** | **28** |  |
| **GO:0010413** | **glucuronoxylan metabolic process** | **3.68858E-05** | **Bpev01.c0015.g0046/Bpev01.c0052.g0019/Bpev01.c0088.g0120/Bpev01.c0288.g0009/Bpev01.c0621.g0009/Bpev01.c0781.g0019/Bpev01.c2001.g0008** | **7** |  |
| **GO:0010417** | **glucuronoxylan biosynthetic process** | **3.68858E-05** | **Bpev01.c0015.g0046/Bpev01.c0052.g0019/Bpev01.c0088.g0120/Bpev01.c0288.g0009/Bpev01.c0621.g0009/Bpev01.c0781.g0019/Bpev01.c2001.g0008** | **7** |  |
| **GO:0045489** | **pectin biosynthetic process** | **7.12079E-05** | **Bpev01.c0015.g0262/Bpev01.c0018.g0088/Bpev01.c0088.g0024/Bpev01.c0163.g0015/Bpev01.c0166.g0033/Bpev01.c0190.g0014/Bpev01.c0265.g0024/Bpev01.c0430.g0012/Bpev01.c0502.g0008/Bpev01.c0621.g0009/Bpev01.c0781.g0019/Bpev01.c0848.g0015/Bpev01.c2157.g0002** | **13** |  |
| **GO:0046271** | **phenylpropanoid catabolic process** | **8.53811E-05** | **Bpev01.c0115.g0060/Bpev01.c0146.g0026/Bpev01.c0150.g0006/Bpev01.c0190.g0082/Bpev01.c0383.g0004/Bpev01.c0803.g0009/Bpev01.c0817.g0012/Bpev01.c1378.g0027/Bpev01.c1786.g0017/Bpev01.c2473.g0001** | **10** |  |
| **GO:0044262** | **cellular carbohydrate metabolic process** | **8.73991E-05** | **Bpev01.c0000.g0006/Bpev01.c0015.g0038/Bpev01.c0015.g0046/Bpev01.c0016.g0180/Bpev01.c0029.g0110/Bpev01.c0052.g0019/Bpev01.c0088.g0120/Bpev01.c0115.g0065/Bpev01.c0151.g0003/Bpev01.c0154.g0062/Bpev01.c0166.g0009/Bpev01.c0166.g0033/Bpev01.c0189.g0059/Bpev01.c0190.g0014/Bpev01.c0214.g0029/Bpev01.c0253.g0016/Bpev01.c0265.g0024/Bpev01.c0275.g0062/Bpev01.c0288.g0009/Bpev01.c0332.g0002/Bpev01.c0344.g0013/Bpev01.c0374.g0017/Bpev01.c0430.g0012/Bpev01.c0522.g0023/Bpev01.c0522.g0024/Bpev01.c0522.g0040/Bpev01.c0582.g0012/Bpev01.c0598.g0015/Bpev01.c0603.g0003/Bpev01.c0621.g0009/Bpev01.c0679.g0017/Bpev01.c0739.g0007/Bpev01.c0781.g0019/Bpev01.c0845.g0002/Bpev01.c0892.g0012/Bpev01.c0892.g0015/Bpev01.c0892.g0016/Bpev01.c0901.g0009/Bpev01.c1082.g0001/Bpev01.c1288.g0001/Bpev01.c1659.g0003/Bpev01.c2001.g0008/Bpev01.c2157.g0002** | **43** |  |
| **GO:0045488** | **pectin metabolic process** | **0.000443549** | **Bpev01.c0015.g0262/Bpev01.c0018.g0088/Bpev01.c0044.g0114/Bpev01.c0051.g0197/Bpev01.c0051.g0198/Bpev01.c0088.g0024/Bpev01.c0163.g0015/Bpev01.c0166.g0033/Bpev01.c0190.g0014/Bpev01.c0265.g0024/Bpev01.c0332.g0002/Bpev01.c0430.g0012/Bpev01.c0502.g0008/Bpev01.c0603.g0003/Bpev01.c0621.g0009/Bpev01.c0781.g0019/Bpev01.c0848.g0015/Bpev01.c2001.g0009/Bpev01.c2157.g0002** | **19** |  |
| **GO:0046274** | **lignin catabolic process** | **0.000474193** | **Bpev01.c0115.g0060/Bpev01.c0146.g0026/Bpev01.c0150.g0006/Bpev01.c0190.g0082/Bpev01.c0383.g0004/Bpev01.c0817.g0012/Bpev01.c1378.g0027/Bpev01.c1786.g0017/Bpev01.c2473.g0001** | **9** |  |
| **GO:0010393** | **galacturonan metabolic process** | **0.000675539** | **Bpev01.c0015.g0262/Bpev01.c0018.g0088/Bpev01.c0044.g0114/Bpev01.c0051.g0197/Bpev01.c0051.g0198/Bpev01.c0088.g0024/Bpev01.c0163.g0015/Bpev01.c0166.g0033/Bpev01.c0190.g0014/Bpev01.c0265.g0024/Bpev01.c0332.g0002/Bpev01.c0430.g0012/Bpev01.c0502.g0008/Bpev01.c0603.g0003/Bpev01.c0621.g0009/Bpev01.c0781.g0019/Bpev01.c0848.g0015/Bpev01.c2001.g0009/Bpev01.c2157.g0002** | **19** |  |
| **GO:0009699** | **phenylpropanoid biosynthetic process** | **0.001064638** | **Bpev01.c0042.g0040/Bpev01.c0095.g0022/Bpev01.c0146.g0026/Bpev01.c0150.g0006/Bpev01.c0190.g0082/Bpev01.c0210.g0051/Bpev01.c0383.g0004/Bpev01.c0420.g0004/Bpev01.c0803.g0009/Bpev01.c0817.g0012/Bpev01.c0833.g0002/Bpev01.c1097.g0002/Bpev01.c1378.g0027/Bpev01.c1786.g0017/Bpev01.c2473.g0001/Bpev01.c2500.g0002** | **16** |  |
| **GO:0006073** | **cellular glucan metabolic process** | **0.001303654** | **Bpev01.c0000.g0006/Bpev01.c0016.g0180/Bpev01.c0029.g0110/Bpev01.c0151.g0003/Bpev01.c0166.g0009/Bpev01.c0189.g0059/Bpev01.c0190.g0014/Bpev01.c0214.g0029/Bpev01.c0253.g0016/Bpev01.c0332.g0002/Bpev01.c0374.g0017/Bpev01.c0522.g0023/Bpev01.c0522.g0024/Bpev01.c0522.g0040/Bpev01.c0582.g0012/Bpev01.c0598.g0015/Bpev01.c0603.g0003/Bpev01.c0679.g0017/Bpev01.c0845.g0002/Bpev01.c0892.g0012/Bpev01.c0892.g0015/Bpev01.c0892.g0016/Bpev01.c0901.g0009/Bpev01.c1082.g0001/Bpev01.c1659.g0003** | **25** |  |
| **GO:0044042** | **glucan metabolic process** | **0.001303654** | **Bpev01.c0000.g0006/Bpev01.c0016.g0180/Bpev01.c0029.g0110/Bpev01.c0151.g0003/Bpev01.c0166.g0009/Bpev01.c0189.g0059/Bpev01.c0190.g0014/Bpev01.c0214.g0029/Bpev01.c0253.g0016/Bpev01.c0332.g0002/Bpev01.c0374.g0017/Bpev01.c0522.g0023/Bpev01.c0522.g0024/Bpev01.c0522.g0040/Bpev01.c0582.g0012/Bpev01.c0598.g0015/Bpev01.c0603.g0003/Bpev01.c0679.g0017/Bpev01.c0845.g0002/Bpev01.c0892.g0012/Bpev01.c0892.g0015/Bpev01.c0892.g0016/Bpev01.c0901.g0009/Bpev01.c1082.g0001/Bpev01.c1659.g0003** | **25** |  |
| **GO:0009698** | **phenylpropanoid metabolic process** | **0.001346435** | **Bpev01.c0042.g0040/Bpev01.c0095.g0022/Bpev01.c0115.g0060/Bpev01.c0146.g0026/Bpev01.c0150.g0006/Bpev01.c0190.g0082/Bpev01.c0210.g0051/Bpev01.c0383.g0004/Bpev01.c0420.g0004/Bpev01.c0617.g0016/Bpev01.c0803.g0009/Bpev01.c0817.g0012/Bpev01.c0833.g0002/Bpev01.c1097.g0002/Bpev01.c1378.g0027/Bpev01.c1786.g0017/Bpev01.c2473.g0001/Bpev01.c2500.g0002** | **18** |  |
| **GO:0009664** | **plant-type cell wall organization** | **0.00261652** | **Bpev01.c0018.g0088/Bpev01.c0029.g0110/Bpev01.c0163.g0015/Bpev01.c0166.g0009/Bpev01.c0166.g0033/Bpev01.c0189.g0059/Bpev01.c0190.g0014/Bpev01.c0265.g0024/Bpev01.c0332.g0002/Bpev01.c0603.g0003/Bpev01.c0639.g0023/Bpev01.c0727.g0003/Bpev01.c0920.g0001/Bpev01.c1484.g0017/Bpev01.c1596.g0001/Bpev01.c2157.g0002** | **16** |  |
| **GO:0044550** | **secondary metabolite biosynthetic process** | **0.00506703** | **Bpev01.c0042.g0040/Bpev01.c0095.g0022/Bpev01.c0146.g0026/Bpev01.c0150.g0006/Bpev01.c0190.g0082/Bpev01.c0210.g0051/Bpev01.c0383.g0004/Bpev01.c0420.g0004/Bpev01.c0803.g0009/Bpev01.c0817.g0012/Bpev01.c0833.g0002/Bpev01.c1097.g0002/Bpev01.c1378.g0027/Bpev01.c1786.g0017/Bpev01.c2473.g0001/Bpev01.c2500.g0002/Bpev01.c3183.g0002** | **17** |  |
| **GO:0019748** | **secondary metabolic process** | **0.006355013** | **Bpev01.c0042.g0040/Bpev01.c0073.g0013/Bpev01.c0095.g0022/Bpev01.c0115.g0060/Bpev01.c0146.g0026/Bpev01.c0150.g0006/Bpev01.c0161.g0086/Bpev01.c0190.g0082/Bpev01.c0210.g0051/Bpev01.c0383.g0004/Bpev01.c0420.g0004/Bpev01.c0617.g0016/Bpev01.c0803.g0009/Bpev01.c0817.g0012/Bpev01.c0833.g0002/Bpev01.c1097.g0002/Bpev01.c1378.g0027/Bpev01.c1786.g0017/Bpev01.c2473.g0001/Bpev01.c2500.g0002/Bpev01.c3183.g0002** | **21** |  |
| **GO:0007010** | **cytoskeleton organization** | **0.007697305** | **Bpev01.c0025.g0003/Bpev01.c0052.g0119/Bpev01.c0213.g0014/Bpev01.c0214.g0051/Bpev01.c0307.g0004/Bpev01.c0343.g0030/Bpev01.c0389.g0057/Bpev01.c0442.g0036/Bpev01.c0574.g0005/Bpev01.c0597.g0026/Bpev01.c0598.g0015/Bpev01.c0621.g0013/Bpev01.c0639.g0024/Bpev01.c0645.g0032/Bpev01.c0817.g0020/Bpev01.c0893.g0006/Bpev01.c0948.g0001/Bpev01.c1335.g0008/Bpev01.c1805.g0001/Bpev01.c2160.g0006/Bpev01.c2443.g0001** | **21** |  |
| **GO:0009809** | **lignin biosynthetic process** | **0.00874559** | **Bpev01.c0095.g0022/Bpev01.c0146.g0026/Bpev01.c0150.g0006/Bpev01.c0190.g0082/Bpev01.c0210.g0051/Bpev01.c0383.g0004/Bpev01.c0817.g0012/Bpev01.c1378.g0027/Bpev01.c1786.g0017/Bpev01.c2473.g0001** | **10** |  |
| **GO:0007017** | **microtubule-based process** | **0.00874559** | **Bpev01.c0052.g0119/Bpev01.c0101.g0038/Bpev01.c0213.g0014/Bpev01.c0214.g0051/Bpev01.c0240.g0005/Bpev01.c0307.g0004/Bpev01.c0442.g0036/Bpev01.c0597.g0026/Bpev01.c0598.g0015/Bpev01.c0621.g0013/Bpev01.c0645.g0032/Bpev01.c0828.g0018/Bpev01.c0909.g0005/Bpev01.c1112.g0008/Bpev01.c1211.g0019/Bpev01.c1287.g0001/Bpev01.c1335.g0008/Bpev01.c1805.g0001/Bpev01.c2443.g0001** | **19** |  |
| **GO:0051273** | **beta-glucan metabolic process** | **0.010535467** | **Bpev01.c0000.g0006/Bpev01.c0029.g0110/Bpev01.c0151.g0003/Bpev01.c0166.g0009/Bpev01.c0189.g0059/Bpev01.c0190.g0014/Bpev01.c0214.g0029/Bpev01.c0253.g0016/Bpev01.c0374.g0017/Bpev01.c0522.g0023/Bpev01.c0522.g0024/Bpev01.c0582.g0012/Bpev01.c0598.g0015/Bpev01.c0603.g0003/Bpev01.c0845.g0002** | **15** |  |
| **GO:0009250** | **glucan biosynthetic process** | **0.015443637** | **Bpev01.c0000.g0006/Bpev01.c0029.g0110/Bpev01.c0151.g0003/Bpev01.c0166.g0009/Bpev01.c0189.g0059/Bpev01.c0190.g0014/Bpev01.c0253.g0016/Bpev01.c0374.g0017/Bpev01.c0522.g0040/Bpev01.c0582.g0012/Bpev01.c0598.g0015/Bpev01.c0603.g0003/Bpev01.c0679.g0017/Bpev01.c0845.g0002/Bpev01.c1082.g0001** | **15** |  |
| **GO:0030243** | **cellulose metabolic process** | **0.017939104** | **Bpev01.c0000.g0006/Bpev01.c0029.g0110/Bpev01.c0151.g0003/Bpev01.c0166.g0009/Bpev01.c0189.g0059/Bpev01.c0190.g0014/Bpev01.c0214.g0029/Bpev01.c0374.g0017/Bpev01.c0522.g0023/Bpev01.c0522.g0024/Bpev01.c0582.g0012/Bpev01.c0598.g0015/Bpev01.c0603.g0003** | **13** |  |
| **GO:0051274** | **beta-glucan biosynthetic process** | **0.018445133** | **Bpev01.c0000.g0006/Bpev01.c0029.g0110/Bpev01.c0151.g0003/Bpev01.c0166.g0009/Bpev01.c0189.g0059/Bpev01.c0190.g0014/Bpev01.c0253.g0016/Bpev01.c0374.g0017/Bpev01.c0582.g0012/Bpev01.c0598.g0015/Bpev01.c0603.g0003/Bpev01.c0845.g0002** | **12** |  |
| **GO:0009808** | **lignin metabolic process** | **0.025586237** | **Bpev01.c0095.g0022/Bpev01.c0115.g0060/Bpev01.c0146.g0026/Bpev01.c0150.g0006/Bpev01.c0190.g0082/Bpev01.c0210.g0051/Bpev01.c0383.g0004/Bpev01.c0817.g0012/Bpev01.c1378.g0027/Bpev01.c1786.g0017/Bpev01.c2473.g0001** | **11** |  |
| **GO:0030244** | **cellulose biosynthetic process** | **0.029798939** | **Bpev01.c0000.g0006/Bpev01.c0029.g0110/Bpev01.c0151.g0003/Bpev01.c0166.g0009/Bpev01.c0189.g0059/Bpev01.c0190.g0014/Bpev01.c0374.g0017/Bpev01.c0582.g0012/Bpev01.c0598.g0015/Bpev01.c0603.g0003** | **10** |  |
| **GO:0031507** | **heterochromatin assembly** | **0.032280455** | **Bpev01.c0205.g0025/Bpev01.c0452.g0007/Bpev01.c0874.g0014/Bpev01.c1202.g0019/Bpev01.c2417.g0001** | **5** |  |
| **GO:0097435** | **supramolecular fiber organization** | **0.036636651** | **Bpev01.c0025.g0003/Bpev01.c0343.g0030/Bpev01.c0389.g0057/Bpev01.c0574.g0005/Bpev01.c0597.g0026/Bpev01.c0598.g0015/Bpev01.c0639.g0024/Bpev01.c0817.g0020/Bpev01.c0893.g0006/Bpev01.c0948.g0001/Bpev01.c1335.g0008/Bpev01.c2160.g0006/Bpev01.c2443.g0001** | **13** |  |
| **GO:0042254** | **ribosome biogenesis** | **0.009158075** | **Bpev01.c0026.g0010/Bpev01.c0062.g0044/Bpev01.c0121.g0017/Bpev01.c0129.g0102/Bpev01.c0190.g0007/Bpev01.c0206.g0007/Bpev01.c0298.g0013/Bpev01.c0328.g0038/Bpev01.c0352.g0004/Bpev01.c0404.g0026/Bpev01.c0427.g0016/Bpev01.c0457.g0048/Bpev01.c0540.g0021/Bpev01.c0541.g0007/Bpev01.c0677.g0008/Bpev01.c1029.g0001/Bpev01.c1054.g0012/Bpev01.c1539.g0002/Bpev01.c2261.g0001** | **19** | **Cluster 5** |
| **GO:0098542** | **defense response to other organism** | **0.009158075** | **Bpev01.c0022.g0034/Bpev01.c0022.g0035/Bpev01.c0051.g0134/Bpev01.c0078.g0001/Bpev01.c0082.g0047/Bpev01.c0082.g0056/Bpev01.c0105.g0018/Bpev01.c0114.g0055/Bpev01.c0115.g0107/Bpev01.c0121.g0041/Bpev01.c0121.g0045/Bpev01.c0145.g0021/Bpev01.c0155.g0001/Bpev01.c0170.g0008/Bpev01.c0224.g0041/Bpev01.c0279.g0011/Bpev01.c0286.g0040/Bpev01.c0349.g0036/Bpev01.c0357.g0001/Bpev01.c0427.g0008/Bpev01.c0437.g0001/Bpev01.c0439.g0005/Bpev01.c0439.g0007/Bpev01.c0439.g0019/Bpev01.c0442.g0007/Bpev01.c0494.g0001/Bpev01.c0566.g0069/Bpev01.c0568.g0002/Bpev01.c0576.g0003/Bpev01.c0579.g0016/Bpev01.c0659.g0011/Bpev01.c0796.g0003/Bpev01.c0956.g0002/Bpev01.c1021.g0001/Bpev01.c1036.g0001/Bpev01.c1040.g0017/Bpev01.c1170.g0005/Bpev01.c1605.g0001/Bpev01.c1650.g0004/Bpev01.c1816.g0002/Bpev01.c1844.g0004/Bpev01.c2282.g0005/Bpev01.c2316.g0002/Bpev01.c2571.g0003/Bpev01.c3249.g0002/Bpev01.c3848.g0001** | **46** |  |
| **GO:0022613** | **ribonucleoprotein complex biogenesis** | **0.012081485** | **Bpev01.c0026.g0010/Bpev01.c0029.g0154/Bpev01.c0062.g0044/Bpev01.c0121.g0017/Bpev01.c0129.g0102/Bpev01.c0190.g0007/Bpev01.c0206.g0007/Bpev01.c0298.g0013/Bpev01.c0328.g0038/Bpev01.c0352.g0004/Bpev01.c0404.g0026/Bpev01.c0427.g0016/Bpev01.c0457.g0048/Bpev01.c0540.g0021/Bpev01.c0541.g0007/Bpev01.c0547.g0008/Bpev01.c0677.g0008/Bpev01.c0817.g0001/Bpev01.c1029.g0001/Bpev01.c1054.g0012/Bpev01.c1539.g0002/Bpev01.c2261.g0001** | **22** |  |
| **GO:0009611** | **response to wounding** | **0.02350419** | **Bpev01.c0027.g0075/Bpev01.c0105.g0018/Bpev01.c0115.g0107/Bpev01.c0189.g0010/Bpev01.c0279.g0011/Bpev01.c0390.g0007/Bpev01.c0556.g0017/Bpev01.c0796.g0003/Bpev01.c0821.g0011/Bpev01.c0956.g0002/Bpev01.c1036.g0001/Bpev01.c1086.g0011/Bpev01.c1754.g0001/Bpev01.c1798.g0004** | **14** |  |
| **GO:0006558** | **L-phenylalanine metabolic process** | **0.02350419** | **Bpev01.c0000.g0226/Bpev01.c0051.g0086/Bpev01.c0051.g0089/Bpev01.c0884.g0007/Bpev01.c1165.g0003/Bpev01.c1165.g0004** | **6** |  |
| **GO:1902221** | **erythrose 4-phosphate/phosphoenolpyruvate family amino acid metabolic process** | **0.02350419** | **Bpev01.c0000.g0226/Bpev01.c0051.g0086/Bpev01.c0051.g0089/Bpev01.c0884.g0007/Bpev01.c1165.g0003/Bpev01.c1165.g0004** | **6** |  |
| **GO:0050832** | **defense response to fungus** | **0.023817391** | **Bpev01.c0082.g0047/Bpev01.c0082.g0056/Bpev01.c0114.g0055/Bpev01.c0115.g0107/Bpev01.c0121.g0041/Bpev01.c0121.g0045/Bpev01.c0170.g0008/Bpev01.c0224.g0041/Bpev01.c0279.g0011/Bpev01.c0427.g0008/Bpev01.c0437.g0001/Bpev01.c0442.g0007/Bpev01.c0494.g0001/Bpev01.c0579.g0016/Bpev01.c1021.g0001/Bpev01.c1170.g0005/Bpev01.c1605.g0001/Bpev01.c1650.g0004/Bpev01.c1816.g0002/Bpev01.c3848.g0001** | **20** |  |
| **GO:0009657** | **plastid organization** | **0.005735327** | **Bpev01.c0015.g0033/Bpev01.c0088.g0011/Bpev01.c0134.g0091/Bpev01.c0171.g0018/Bpev01.c0213.g0004/Bpev01.c0243.g0037/Bpev01.c0286.g0029/Bpev01.c0386.g0014/Bpev01.c0467.g0017/Bpev01.c0484.g0019/Bpev01.c0572.g0010/Bpev01.c0633.g0007/Bpev01.c0884.g0013/Bpev01.c1060.g0027/Bpev01.c1435.g0009/Bpev01.c1674.g0006/Bpev01.c1689.g0006** | **17** | **Cluster 6** |
| **GO:0009658** | **chloroplast organization** | **0.005735327** | **Bpev01.c0015.g0033/Bpev01.c0088.g0011/Bpev01.c0134.g0091/Bpev01.c0171.g0018/Bpev01.c0213.g0004/Bpev01.c0243.g0037/Bpev01.c0286.g0029/Bpev01.c0386.g0014/Bpev01.c0484.g0019/Bpev01.c0572.g0010/Bpev01.c0633.g0007/Bpev01.c0884.g0013/Bpev01.c1435.g0009/Bpev01.c1674.g0006/Bpev01.c1689.g0006** | **15** |  |
| **GO:0031425** | **chloroplast RNA processing** | **0.012516404** | **Bpev01.c0029.g0125/Bpev01.c0101.g0081/Bpev01.c0134.g0091/Bpev01.c0243.g0037/Bpev01.c0981.g0007/Bpev01.c1689.g0006** | **6** |  |
| **GO:0015979** | **photosynthesis** | **1.10476E-06** | **Bpev01.c0009.g0001/Bpev01.c0015.g0245/Bpev01.c0022.g0041/Bpev01.c0042.g0007/Bpev01.c0052.g0108/Bpev01.c0062.g0075/Bpev01.c0080.g0006/Bpev01.c0080.g0094/Bpev01.c0094.g0002/Bpev01.c0147.g0018/Bpev01.c0162.g0029/Bpev01.c0210.g0009/Bpev01.c0212.g0008/Bpev01.c0224.g0043/Bpev01.c0245.g0030/Bpev01.c0329.g0002/Bpev01.c0340.g0026/Bpev01.c0355.g0003/Bpev01.c0362.g0012/Bpev01.c0373.g0010/Bpev01.c0407.g0010/Bpev01.c0454.g0016/Bpev01.c0480.g0020/Bpev01.c0517.g0008/Bpev01.c0540.g0029/Bpev01.c0595.g0009/Bpev01.c0621.g0012/Bpev01.c0717.g0022/Bpev01.c0796.g0011/Bpev01.c1040.g0049/Bpev01.c1116.g0009/Bpev01.c1138.g0008/Bpev01.c1275.g0002/Bpev01.c1708.g0002/Bpev01.c1767.g0010/Bpev01.c1891.g0002** | **36** | **Cluster 9** |
| **GO:0031407** | **oxylipin metabolic process** | **0.001833524** | **Bpev01.c0094.g0025/Bpev01.c0094.g0026/Bpev01.c0473.g0005/Bpev01.c0523.g0011/Bpev01.c0523.g0013/Bpev01.c0555.g0007/Bpev01.c1365.g0005/Bpev01.c1400.g0005/Bpev01.c1828.g0002/Bpev01.c2571.g0002/Bpev01.c2571.g0004** | **11** |  |
| **GO:0031408** | **oxylipin biosynthetic process** | **0.001833524** | **Bpev01.c0094.g0025/Bpev01.c0094.g0026/Bpev01.c0473.g0005/Bpev01.c0523.g0011/Bpev01.c0523.g0013/Bpev01.c0555.g0007/Bpev01.c1365.g0005/Bpev01.c1400.g0005/Bpev01.c1828.g0002/Bpev01.c2571.g0002/Bpev01.c2571.g0004** | **11** |  |
| **GO:0019684** | **photosynthesis, light reaction** | **0.003334561** | **Bpev01.c0009.g0001/Bpev01.c0015.g0245/Bpev01.c0042.g0007/Bpev01.c0052.g0108/Bpev01.c0080.g0094/Bpev01.c0094.g0002/Bpev01.c0162.g0029/Bpev01.c0224.g0043/Bpev01.c0362.g0012/Bpev01.c0373.g0010/Bpev01.c0517.g0008/Bpev01.c0595.g0009/Bpev01.c0621.g0012/Bpev01.c0717.g0022/Bpev01.c0796.g0011/Bpev01.c1040.g0049/Bpev01.c1138.g0008/Bpev01.c1767.g0010/Bpev01.c1891.g0002** | **19** |  |
| **GO:0016052** | **carbohydrate catabolic process** | **0.003334561** | **Bpev01.c0023.g0007/Bpev01.c0044.g0089/Bpev01.c0044.g0090/Bpev01.c0112.g0017/Bpev01.c0127.g0037/Bpev01.c0142.g0009/Bpev01.c0173.g0011/Bpev01.c0223.g0011/Bpev01.c0228.g0005/Bpev01.c0274.g0048/Bpev01.c0420.g0017/Bpev01.c0420.g0024/Bpev01.c0477.g0023/Bpev01.c0497.g0017/Bpev01.c0497.g0018/Bpev01.c0497.g0019/Bpev01.c0679.g0011/Bpev01.c0800.g0025/Bpev01.c0818.g0005/Bpev01.c1060.g0025/Bpev01.c1067.g0009/Bpev01.c1105.g0007/Bpev01.c1136.g0004/Bpev01.c1282.g0001/Bpev01.c1389.g0003/Bpev01.c1408.g0004/Bpev01.c1411.g0006/Bpev01.c1470.g0001/Bpev01.c1534.g0006/Bpev01.c1534.g0007/Bpev01.c1534.g0008/Bpev01.c1726.g0002/Bpev01.c1736.g0001/Bpev01.c2331.g0001** | **34** |  |
| **GO:0071241** | **cellular response to inorganic substance** | **0.008564583** | **Bpev01.c0016.g0138/Bpev01.c0094.g0025/Bpev01.c0094.g0026/Bpev01.c0120.g0044/Bpev01.c0166.g0019/Bpev01.c0222.g0002/Bpev01.c0443.g0026/Bpev01.c0804.g0002/Bpev01.c0804.g0003/Bpev01.c1544.g0007** | **10** |  |
| **GO:0000272** | **polysaccharide catabolic process** | **0.008564583** | **Bpev01.c0023.g0007/Bpev01.c0112.g0017/Bpev01.c0127.g0037/Bpev01.c0173.g0011/Bpev01.c0228.g0005/Bpev01.c0420.g0017/Bpev01.c0420.g0024/Bpev01.c0497.g0017/Bpev01.c0497.g0018/Bpev01.c0497.g0019/Bpev01.c0679.g0011/Bpev01.c0818.g0005/Bpev01.c1060.g0025/Bpev01.c1282.g0001/Bpev01.c1389.g0003/Bpev01.c1408.g0004/Bpev01.c1470.g0001/Bpev01.c1534.g0006/Bpev01.c1534.g0007/Bpev01.c1534.g0008/Bpev01.c1726.g0002/Bpev01.c1736.g0001/Bpev01.c2331.g0001** | **23** |  |
| **GO:0097305** | **response to alcohol** | **0.008753676** | **Bpev01.c0016.g0175/Bpev01.c0038.g0120/Bpev01.c0052.g0056/Bpev01.c0052.g0093/Bpev01.c0052.g0196/Bpev01.c0062.g0034/Bpev01.c0089.g0019/Bpev01.c0094.g0025/Bpev01.c0094.g0026/Bpev01.c0115.g0098/Bpev01.c0120.g0044/Bpev01.c0163.g0028/Bpev01.c0190.g0010/Bpev01.c0215.g0034/Bpev01.c0222.g0002/Bpev01.c0329.g0008/Bpev01.c0329.g0009/Bpev01.c0329.g0010/Bpev01.c0357.g0068/Bpev01.c0365.g0004/Bpev01.c0366.g0025/Bpev01.c0437.g0007/Bpev01.c0439.g0048/Bpev01.c0455.g0029/Bpev01.c0497.g0017/Bpev01.c0497.g0018/Bpev01.c0497.g0019/Bpev01.c0557.g0034/Bpev01.c0578.g0015/Bpev01.c0781.g0015/Bpev01.c1085.g0017/Bpev01.c1085.g0021/Bpev01.c1085.g0022/Bpev01.c1085.g0024/Bpev01.c1143.g0001/Bpev01.c1259.g0010/Bpev01.c1292.g0002/Bpev01.c2190.g0001/Bpev01.c2243.g0001/Bpev01.c4953.g0001** | **40** |  |
| **GO:0009737** | **response to abscisic acid** | **0.008753676** | **Bpev01.c0016.g0175/Bpev01.c0038.g0120/Bpev01.c0052.g0056/Bpev01.c0052.g0093/Bpev01.c0052.g0196/Bpev01.c0062.g0034/Bpev01.c0094.g0025/Bpev01.c0094.g0026/Bpev01.c0115.g0098/Bpev01.c0120.g0044/Bpev01.c0163.g0028/Bpev01.c0190.g0010/Bpev01.c0215.g0034/Bpev01.c0222.g0002/Bpev01.c0329.g0008/Bpev01.c0329.g0009/Bpev01.c0329.g0010/Bpev01.c0357.g0068/Bpev01.c0365.g0004/Bpev01.c0366.g0025/Bpev01.c0437.g0007/Bpev01.c0439.g0048/Bpev01.c0455.g0029/Bpev01.c0497.g0017/Bpev01.c0497.g0018/Bpev01.c0497.g0019/Bpev01.c0557.g0034/Bpev01.c0578.g0015/Bpev01.c0781.g0015/Bpev01.c1085.g0017/Bpev01.c1085.g0021/Bpev01.c1085.g0022/Bpev01.c1085.g0024/Bpev01.c1143.g0001/Bpev01.c1259.g0010/Bpev01.c1292.g0002/Bpev01.c2190.g0001/Bpev01.c2243.g0001/Bpev01.c4953.g0001** | **39** |  |
| **GO:0010035** | **response to inorganic substance** | **0.009949003** | **Bpev01.c0016.g0138/Bpev01.c0038.g0120/Bpev01.c0052.g0093/Bpev01.c0052.g0196/Bpev01.c0062.g0034/Bpev01.c0080.g0104/Bpev01.c0092.g0017/Bpev01.c0094.g0025/Bpev01.c0094.g0026/Bpev01.c0114.g0054/Bpev01.c0115.g0098/Bpev01.c0120.g0044/Bpev01.c0129.g0055/Bpev01.c0142.g0009/Bpev01.c0142.g0068/Bpev01.c0163.g0028/Bpev01.c0166.g0019/Bpev01.c0222.g0002/Bpev01.c0228.g0005/Bpev01.c0245.g0007/Bpev01.c0329.g0008/Bpev01.c0329.g0009/Bpev01.c0329.g0010/Bpev01.c0329.g0013/Bpev01.c0343.g0021/Bpev01.c0437.g0007/Bpev01.c0439.g0048/Bpev01.c0442.g0039/Bpev01.c0443.g0026/Bpev01.c0455.g0029/Bpev01.c0467.g0016/Bpev01.c0522.g0014/Bpev01.c0555.g0007/Bpev01.c0577.g0002/Bpev01.c0578.g0015/Bpev01.c0640.g0005/Bpev01.c0717.g0010/Bpev01.c0804.g0002/Bpev01.c0804.g0003/Bpev01.c0879.g0002/Bpev01.c0901.g0016/Bpev01.c1086.g0008/Bpev01.c1135.g0005/Bpev01.c1136.g0004/Bpev01.c1143.g0001/Bpev01.c1222.g0002/Bpev01.c1290.g0001/Bpev01.c1292.g0002/Bpev01.c1465.g0009/Bpev01.c1544.g0007/Bpev01.c2571.g0002/Bpev01.c2571.g0004** | **52** |  |
| **GO:0033993** | **response to lipid** | **0.011975152** | **Bpev01.c0000.g0045/Bpev01.c0015.g0205/Bpev01.c0016.g0175/Bpev01.c0038.g0120/Bpev01.c0052.g0056/Bpev01.c0052.g0093/Bpev01.c0052.g0196/Bpev01.c0062.g0034/Bpev01.c0094.g0025/Bpev01.c0094.g0026/Bpev01.c0115.g0098/Bpev01.c0120.g0044/Bpev01.c0163.g0028/Bpev01.c0190.g0010/Bpev01.c0215.g0034/Bpev01.c0222.g0002/Bpev01.c0329.g0008/Bpev01.c0329.g0009/Bpev01.c0329.g0010/Bpev01.c0357.g0068/Bpev01.c0363.g0019/Bpev01.c0365.g0004/Bpev01.c0366.g0025/Bpev01.c0437.g0007/Bpev01.c0439.g0048/Bpev01.c0455.g0029/Bpev01.c0497.g0017/Bpev01.c0497.g0018/Bpev01.c0497.g0019/Bpev01.c0557.g0034/Bpev01.c0578.g0015/Bpev01.c0671.g0019/Bpev01.c0781.g0015/Bpev01.c1085.g0017/Bpev01.c1085.g0021/Bpev01.c1085.g0022/Bpev01.c1085.g0024/Bpev01.c1143.g0001/Bpev01.c1202.g0045/Bpev01.c1259.g0010/Bpev01.c1292.g0002/Bpev01.c1477.g0005/Bpev01.c2190.g0001/Bpev01.c2243.g0001/Bpev01.c3140.g0002/Bpev01.c4953.g0001** | **46** |  |
| **GO:0009765** | **photosynthesis, light harvesting** | **0.015608462** | **Bpev01.c0042.g0007/Bpev01.c0052.g0108/Bpev01.c0080.g0094/Bpev01.c0362.g0012/Bpev01.c0517.g0008/Bpev01.c0796.g0011/Bpev01.c1040.g0049/Bpev01.c1138.g0008/Bpev01.c1767.g0010** | **9** |  |
| **GO:0009414** | **response to water deprivation** | **0.020217691** | **Bpev01.c0038.g0120/Bpev01.c0052.g0093/Bpev01.c0052.g0196/Bpev01.c0062.g0034/Bpev01.c0092.g0017/Bpev01.c0114.g0054/Bpev01.c0115.g0098/Bpev01.c0129.g0055/Bpev01.c0142.g0068/Bpev01.c0163.g0028/Bpev01.c0222.g0002/Bpev01.c0228.g0005/Bpev01.c0329.g0008/Bpev01.c0329.g0009/Bpev01.c0329.g0010/Bpev01.c0343.g0021/Bpev01.c0437.g0007/Bpev01.c0443.g0026/Bpev01.c0455.g0029/Bpev01.c0522.g0014/Bpev01.c0577.g0002/Bpev01.c0578.g0015/Bpev01.c0640.g0005/Bpev01.c1143.g0001/Bpev01.c1290.g0001/Bpev01.c1292.g0002** | **26** |  |
| **GO:0009415** | **response to water** | **0.023155761** | **Bpev01.c0038.g0120/Bpev01.c0052.g0093/Bpev01.c0052.g0196/Bpev01.c0062.g0034/Bpev01.c0092.g0017/Bpev01.c0114.g0054/Bpev01.c0115.g0098/Bpev01.c0129.g0055/Bpev01.c0142.g0068/Bpev01.c0163.g0028/Bpev01.c0222.g0002/Bpev01.c0228.g0005/Bpev01.c0329.g0008/Bpev01.c0329.g0009/Bpev01.c0329.g0010/Bpev01.c0343.g0021/Bpev01.c0437.g0007/Bpev01.c0443.g0026/Bpev01.c0455.g0029/Bpev01.c0522.g0014/Bpev01.c0577.g0002/Bpev01.c0578.g0015/Bpev01.c0640.g0005/Bpev01.c1143.g0001/Bpev01.c1290.g0001/Bpev01.c1292.g0002** | **26** |  |
| **GO:0006012** | **galactose metabolic process** | **0.023400964** | **Bpev01.c0142.g0016/Bpev01.c0142.g0017/Bpev01.c0142.g0018/Bpev01.c0278.g0041/Bpev01.c0358.g0003/Bpev01.c0995.g0013/Bpev01.c1104.g0025** | **7** |  |
| **GO:0001101** | **response to acid chemical** | **0.025435104** | **Bpev01.c0038.g0120/Bpev01.c0052.g0093/Bpev01.c0052.g0196/Bpev01.c0062.g0034/Bpev01.c0092.g0017/Bpev01.c0114.g0054/Bpev01.c0115.g0098/Bpev01.c0129.g0055/Bpev01.c0142.g0068/Bpev01.c0163.g0028/Bpev01.c0215.g0034/Bpev01.c0222.g0002/Bpev01.c0228.g0005/Bpev01.c0329.g0008/Bpev01.c0329.g0009/Bpev01.c0329.g0010/Bpev01.c0343.g0021/Bpev01.c0437.g0007/Bpev01.c0443.g0026/Bpev01.c0455.g0029/Bpev01.c0522.g0014/Bpev01.c0577.g0002/Bpev01.c0578.g0015/Bpev01.c0640.g0005/Bpev01.c1143.g0001/Bpev01.c1290.g0001/Bpev01.c1292.g0002** | **27** |  |
| **GO:0006865** | **amino acid transport** | **0.029268023** | **Bpev01.c0017.g0014/Bpev01.c0026.g0005/Bpev01.c0052.g0224/Bpev01.c0391.g0004/Bpev01.c0393.g0001/Bpev01.c0639.g0019/Bpev01.c0667.g0005/Bpev01.c0902.g0013/Bpev01.c1040.g0021/Bpev01.c1128.g0004/Bpev01.c1749.g0001/Bpev01.c2410.g0002** | **12** |  |
| **GO:0009768** | **photosynthesis, light harvesting in photosystem I** | **0.029268023** | **Bpev01.c0042.g0007/Bpev01.c0080.g0094/Bpev01.c0796.g0011/Bpev01.c1040.g0049/Bpev01.c1138.g0008/Bpev01.c1767.g0010** | **6** |  |
