## Supplementary material for "Transcriptome analysis provides insights into candidate genetic regulators of birch response to salt stress": Table 3 Genes related to salt stress response differentially expressed in B. halophila and B. platyphylla.

| Table S3 Genes related to salt stress response differentially expressed in B. halophila and B. platyphylla. | | | | | | |
| --- | --- | --- | --- | --- | --- | --- |
| **Gene ID** | **BHLeaf** | **BPLeaf** | **BPRoot** | Gene ID in Arabidopsis | **Abbreviation name of the gene** | **Full name of the gene** |
| **Bpev01.c0000.g0173** | **0** | **1.467135669** | **0** | **AT1G34370** | **STOP1** | **Sensitive to proton rhizotoxicity 1** |
| **Bpev01.c0594.g0007** | **0** | **1.187869916** | **1.539640597** | **AT1G34370** | **STOP1** | **Sensitive to proton rhizotoxicity 1** |
| **Bpev01.c0241.g0015** | **0** | **-1.614546388** | **0** | **AT1G12820** | **AFB3** | **Auxin signaling F-box 3** |
| **Bpev01.c0568.g0030** | **0** | **1.083341521** | **0** | **AT4G34460** | **AGB1** | **GTP binding protein beta 1** |
| **Bpev01.c0827.g0002** | **0** | **-2.419225075** | **0** | **AT2G34650** | **PID** | **PINOID** |
| **Bpev01.c0564.g0027** | **0** | **1.823831904** | **1.695224001** | **AT3G46550** | **FLA4** | **Fasciclin-like arabinogalactan-protein 4** |
| **Bpev01.c0206.g0004** | **0** | **-1.464708911** | **0** | **AT2G26650** | **AKT1** | **Arabidopsis K+ transporter 1** |
| **Bpev01.c2235.g0014** | **-1.255427457** | **-1.797657468** | **0** | **AT2G26650** | **AKT1** | **Arabidopsis K+ transporter 1** |
| **Bpev01.c0151.g0003** | **0** | **0** | **-1.827407876** | **AT1G35720** | **ANN1** | **Annexin 1** |
| **Bpev01.c0263.g0010** | **-3.88913318** | **0** | **0** | **AT2G38750** | **ANN4** | **Annexin 4** |
| **Bpev01.c0546.g0008** | **0** | **0** | **1.767811324** | **AT2G14210** | **ANR1** | **AGAMOUS-like 44** |
| **Bpev01.c0088.g0098** | **2.665155551** | **2.458671839** | **2.041951506** | **AT1G01720** | **ANAC2** | **Arabidopsis NAC domain containing protein 2** |
| **Bpev01.c0390.g0007** | **0** | **4.034701321** | **1.780214405** | **AT1G01720** | **ANAC2** | **Arabidopsis NAC domain containing protein 2** |
| **Bpev01.c0398.g0003** | **0** | **2.885237856** | **0** | **AT5G43650** | **bHLH092** | **bHLH (basic-helix-loop-helix) transcription factor 92** |
| **Bpev01.c0804.g0002** | **0** | **0** | **2.230647585** | **AT1G08860** | **BON3** | **BONZAI 3** |
| **Bpev01.c0804.g0003** | **0** | **0** | **2.382138545** | **AT1G08860** | **BON3** | **BONZAI 3** |
| **Bpev01.c0018.g0031** | **0** | **3.554835737** | **0** | **AT1G27730** | **STZ** | **Salt tolerance zinc finger** |
| **Bpev01.c0280.g0034** | **0** | **4.191944848** | **0** | **AT1G27730** | **STZ** | **Salt tolerance zinc finger** |
| **Bpev01.c0488.g0029** | **0** | **-1.335883964** | **0** | **AT2G02160** | **C3H17** | **CCCH-type zinc finger protein 17** |
| **Bpev01.c0134.g0106** | **0** | **0** | **-1.848697965** | **AT4G35090** | **CAT2** | **Catalase 2** |
| **Bpev01.c0134.g0111** | **0** | **-1.29615706** | **1.559007902** | **AT4G35090** | **CAT2** | **Catalase 2** |
| **Bpev01.c0710.g0010** | **0** | **0** | **-4.363238279** | **AT3G51860** | **CAX3** | **Cation exchanger 3** |
| **Bpev01.c0333.g0007** | **0** | **-1.952239793** | **0** | **AT4G33000** | **CBL10/SCABP8** | **Calcineurin B-like protein 10/SOS3-like calcium-binding protein 8** |
| **Bpev01.c0389.g0037** | **0** | **-1.053569659** | **0** | **AT1G45688** | **CC1** | **Companion of cellulose synthase 1** |
| **Bpev01.c0389.g0042** | **0** | **-1.453605615** | **0** | **AT1G45688** | **CC1** | **Companion of cellulose synthase 1** |
| **Bpev01.c1116.g0014** | **0** | **0** | **-1.80295224** | **AT1G45688** | **CC1** | **Companion of cellulose synthase 1** |
| **Bpev01.c0328.g0008** | **0** | **1.66091518** | **0** | **AT4G34050** | **CCoAOMT1** | **Caffeoyl coenzyme a o-methyltransferase 1** |
| **Bpev01.c0480.g0087** | **0** | **-1.572420986** | **-1.664227553** | **AT5G64740** | **CesA6** | **Cellulose synthase 6** |
| **Bpev01.c0598.g0015** | **0** | **0** | **-1.343584037** | **AT5G64740** | **CesA6** | **Cellulose synthase 6** |
| **Bpev01.c0374.g0017** | **0** | **0** | **-2.28185** | **AT4G18780** | **CesA8** | **Cellulose synthase 8** |
| **Bpev01.c1067.g0007** | **-1.060393137** | **0** | **0** | **AT5G57630** | **CIPK21** | **CBL-interacting protein kinase 21** |
| **Bpev01.c0578.g0015** | **0** | **0** | **1.366395479** | **AT1G70670** | **CLO4** | **Caleosin 4** |
| **Bpev01.c1480.g0004** | **0** | **-2.390221633** | **7.497846696** | **AT1G01340** | **CNGC10** | **Cyclic nucleotide gated channel 10** |
| **Bpev01.c0025.g0030** | **0** | **0** | **-1.003710741** | **AT4G30360** | **CNGC17** | **Cyclic nucleotide-gated channel 17** |
| **Bpev01.c1463.g0002** | **0** | **0** | **1.426978612** | **AT3G17690** | **CNGC19** | **Cyclic nucleotide gated channel 19** |
| **Bpev01.c2384.g0001** | **0** | **0** | **1.122081941** | **AT3G17700** | **CNGC20** | **Cyclic nucleotide gated channel 20** |
| **Bpev01.c0299.g0035** | **0** | **0** | **1.036169691** | **AT2G32950** | **COP1** | **Constitutive photomorphogenic 1** |
| **Bpev01.c0518.g0018** | **0** | **0** | **1.189587442** | **AT1G35670** | **CPK11/CDPK2** | **Calcium-dependent protein kinase 2** |
| **Bpev01.c1700.g0003** | **0** | **1.226983311** | **1.00226371** | **AT2G17290** | **CPK6/CDPK3** | **Calcium-dependent protein kinase 3** |
| **Bpev01.c0080.g0070** | **0** | **0** | **1.694653986** | **AT1G70520** | **CRK2** | **Cysteine-rich RLK (Receptor-like protein kinase) 2** |
| **Bpev01.c0080.g0071** | **5.761948283** | **0** | **6.462199824** | **AT1G70520** | **CRK2** | **Cysteine-rich RLK (Receptor-like protein kinase) 2** |
| **Bpev01.c0135.g0103** | **0** | **-1.030529176** | **0** | **AT3G51550** | **CrRLK1L/FER** | **FERONIA** |
| **Bpev01.c0164.g0005** | **0** | **0** | **-2.750761365** | **AT3G51550** | **CrRLK1L/FER** | **FERONIA** |
| **Bpev01.c0166.g0004** | **0** | **0** | **-2.769879742** | **AT3G51550** | **CrRLK1L/FER** | **FERONIA** |
| **Bpev01.c0224.g0041** | **0** | **1.3817157** | **0** | **AT3G51550** | **CrRLK1L/FER** | **FERONIA** |
| **Bpev01.c1179.g0005** | **0** | **3.327391495** | **0** | **AT3G51550** | **CrRLK1L/FER** | **FERONIA** |
| **Bpev01.c2243.g0001** | **1.632896873** | **0** | **5.321708496** | **AT3G51550** | **CrRLK1L/FER** | **FERONIA** |
| **Bpev01.c1281.g0013** | **0** | **0** | **-1.391897354** | **AT5G54380** | **CrRLK1L/THE1** | **THESEUS1** |
| **Bpev01.c0364.g0008** | **-1.636534627** | **0** | **-1.223804711** | **AT1G02730** | **CSLD5/SOS6** | **Cellulose synthase-like D5** |
| **Bpev01.c0050.g0052** | **0** | **0** | **2.940520921** | **AT5G63450** | **CYP94B1** | **Cytochrome P450, family 94, subfamily B, polypeptide 1** |
| **Bpev01.c0050.g0054** | **0** | **0** | **2.246623445** | **AT5G63450** | **CYP94B1** | **Cytochrome P450, family 94, subfamily B, polypeptide 1** |
| **Bpev01.c0050.g0058** | **0** | **2.075499636** | **2.668628721** | **AT5G63450** | **CYP94B1** | **Cytochrome P450, family 94, subfamily B, polypeptide 1** |
| **Bpev01.c0050.g0053** | **0** | **0** | **4.172616582** | **AT3G48520** | **CYP94B3** | **Cytochrome P450, family 94, subfamily B, polypeptide 3** |
| **Bpev01.c2045.g0002** | **0** | **0** | **4.425820567** | **AT3G48520** | **CYP94B3** | **Cytochrome P450, family 94, subfamily B, polypeptide 3** |
| **Bpev01.c0145.g0025** | **0** | **0** | **-1.318832863** | **AT1G14920** | **GAI/RGA2** | **Gibberellic acid insensitive/Restoration on growth on ammonia 2** |
| **Bpev01.c2235.g0001** | **0** | **0** | **-1.501224155** | **AT1G14920** | **GAI/RGA2** | **Gibberellic acid insensitive/Restoration on growth on ammonia 2** |
| **Bpev01.c2151.g0001** | **0** | **0** | **-1.365065705** | **AT5G63770** | **DGK2** | **Diacylglycerol kinase 2** |
| **Bpev01.c0142.g0056** | **0** | **1.024023479** | **0** | **AT3G05700** | **Di19-3** | **Drought-induced protein 19** |
| **Bpev01.c0820.g0011** | **0** | **1.14643168** | **0** | **AT3G05700** | **Di19-3** | **Drought-induced protein 19** |
| **Bpev01.c0820.g0012** | **0** | **1.156529821** | **0** | **AT3G05700** | **Di19-3** | **Drought-induced protein 19** |
| **Bpev01.c0073.g0013** | **0** | **0** | **-3.316884752** | **AT1G12610** | **DDF1** | **Dwarf and delayed flowering 1** |
| **Bpev01.c0038.g0113** | **0** | **2.215877293** | **1.566742936** | **AT3G55270** | **DSP-1d/MKP1** | **Mitogen-activated protein kinase phosphatase 1** |
| **Bpev01.c0454.g0017** | **0** | **3.882831594** | **1.728142875** | **AT3G02800** | **DSP-3a/PFA/DSP3** | **Plant and fungi atypical dual-speciﬁcity phosphatase 3** |
| **Bpev01.c1292.g0002** | **2.694043123** | **2.525203076** | **4.18879459** | **AT3G63060** | **EDL3** | **EID1-like 3** |
| **Bpev01.c0050.g0100** | **0** | **5.398477049** | **0** | **AT3G23230** | **ERF098** | **Ethylene-responsive factor 98** |
| **Bpev01.c0764.g0011** | **0** | **5.515708144** | **0** | **AT3G23230** | **ERF098** | **Ethylene-responsive factor 98** |
| **Bpev01.c0764.g0012** | **0** | **4.260502944** | **2.517003839** | **AT3G23230** | **ERF098** | **Ethylene-responsive factor 98** |
| **Bpev01.c0027.g0191** | **0** | **0** | **-2.850254226** | **AT4G34410** | **ERF109** | **Ethylene-responsive factor 109** |
| **Bpev01.c0328.g0028** | **2.309955807** | **0** | **-1.519509791** | **AT4G34410** | **ERF109** | **Ethylene-responsive factor 109** |
| **Bpev01.c0774.g0007** | **0** | **2.34608366** | **0** | **AT3G23150** | **ETR2** | **Ethylene response 2** |
| **Bpev01.c1159.g0015** | **-2.044827465** | **0** | **0** | **AT3G24480** | **LRX4** | **Leucine-rich repeat extensin 4** |
| **Bpev01.c0480.g0073** | **-1.389413412** | **-1.424526567** | **-1.150616608** | **AT2G33570** | **GALS1** | **Domain of unknown function (DUF23)** |
| **Bpev01.c1347.g0002** | **0** | **0** | **1.052104497** | **AT5G05930** | **GC1** | **Guanylyl cyclase 1** |
| **Bpev01.c1086.g0008** | **0** | **0** | **1.345853236** | **AT1G22770** | **GI** | **Gigantea protein** |
| **Bpev01.c0575.g0003** | **0** | **0** | **7.134205547** | **AT1G05200** | **GLR3.4** | **Glutamate receptor 3.4** |
| **Bpev01.c0292.g0003** | **0** | **-1.848236349** | **0** | **AT2G32400** | **GLR3.7** | **Glutamate receptor 3.7** |
| **Bpev01.c0401.g0016** | **1.375494753** | **0** | **3.689498805** | **AT4G26080** | **ABI1** | **ABA insensitive 1** |
| **Bpev01.c0051.g0075** | **1.531999146** | **1.091447858** | **0** | **AT5G57050** | **ABI2** | **ABA insensitive 2** |
| **Bpev01.c0455.g0029** | **0** | **0** | **1.205182069** | **AT2G33700** | **PP2C27** | **Protein phosphatase 2C 27** |
| **Bpev01.c0129.g0031** | **1.156435665** | **2.436335352** | **0** | **AT3G62260** | **PP2C49** | **Protein phosphatase 2C 49** |
| **Bpev01.c0931.g0003** | **0** | **2.719502829** | **1.268378494** | **AT2G30250** | **WRKY25** | **WRKY DNA-binding protein 25** |
| **Bpev01.c0154.g0042** | **0** | **4.291796224** | **0** | **AT2G38470** | **WRKY33** | **WRKY DNA-binding protein 33** |
| **Bpev01.c0357.g0035** | **0** | **0** | **1.097312083** | **AT1G13960** | **WRKY4** | **WRKY DNA-binding protein 4** |
| **Bpev01.c0511.g0003** | **0** | **3.600957678** | **0** | **AT4G31800** | **WRKY18** | **WRKY DNA-binding protein 18** |
| **Bpev01.c0511.g0004** | **0** | **3.690595813** | **1.897447646** | **AT4G31800** | **WRKY18** | **WRKY DNA-binding protein 18** |
| **Bpev01.c0044.g0060** | **0** | **3.765414823** | **0** | **AT1G80840** | **WRKY40** | **WRKY DNA-binding protein 40** |
| **Bpev01.c1331.g0002** | **0** | **0** | **3.038622598** | **AT1G68150** | **WRKY9** | **WRKY DNA-binding protein 9** |
| **Bpev01.c0362.g0010** | **0** | **1.6167037** | **0** | **AT4G18170** | **WRKY28** | **WRKY DNA-binding protein 28** |
| **Bpev01.c1530.g0008** | **0** | **2.513795604** | **2.779231306** | **AT4G18170** | **WRKY28** | **WRKY DNA-binding protein 28** |
| **Bpev01.c0078.g0001** | **0** | **2.551427132** | **0** | **AT4G31550** | **WRKY11** | **WRKY DNA-binding protein 11** |
| **Bpev01.c0606.g0010** | **0** | **3.245868829** | **-1.024855747** | **AT4G31550** | **WRKY11** | **WRKY DNA-binding protein 11** |
| **Bpev01.c0289.g0003** | **0** | **0** | **1.737217428** | **AT1G78380** | **GSTU19** | **Glutathione S-transferase TAU 19** |
| **Bpev01.c0289.g0007** | **0** | **2.083878593** | **0** | **AT1G78380** | **GSTU19** | **Glutathione S-transferase TAU 19** |
| **Bpev01.c0223.g0024** | **0** | **-1.262785008** | **0** | **AT3G09150** | **GUN3/HY2** | **Genomes uncoupled 3/Elongated hypocotyl 2** |
| **Bpev01.c0052.g0003** | **6.057037092** | **0** | **0** | **AT4G13420** | **HAK5** | **High affinity K+ transporter 5** |
| **Bpev01.c0052.g0004** | **0** | **0** | **1.338052442** | **AT4G13420** | **HAK5** | **High affinity K+ transporter 5** |
| **Bpev01.c0088.g0072** | **0** | **0** | **4.334383789** | **AT4G13420** | **HAK5** | **High affinity K+ transporter 5** |
| **Bpev01.c0579.g0001** | **0** | **3.66497354** | **0** | **AT4G13420** | **HAK5** | **High affinity K+ transporter 5** |
| **Bpev01.c0579.g0002** | **0** | **3.997621102** | **2.51821169** | **AT4G13420** | **HAK5** | **High affinity K+ transporter 5** |
| **Bpev01.c0579.g0003** | **0** | **3.83574957** | **2.34953763** | **AT4G13420** | **HAK5** | **High affinity K+ transporter 5** |
| **Bpev01.c0902.g0018** | **0** | **1.32748293** | **0** | **AT5G03740** | **HD2C** | **Histone deacetylase 2C** |
| **Bpev01.c2132.g0002** | **0** | **2.016258957** | **0** | **AT2G01430** | **ATHB17** | **Homeobox-leucine zipper protein 17** |
| **Bpev01.c0022.g0007** | **0** | **0** | **1.000191352** | **AT2G18050** | **HIS1-3** | **Histone H1-3** |
| **Bpev01.c0051.g0059** | **2.450364636** | **0** | **1.390991987** | **AT2G26150** | **HSFA2** | **Heat shock transcription factor A2** |
| **Bpev01.c0237.g0047** | **0** | **2.722924143** | **-1.290408803** | **AT2G26150** | **HSFA2** | **Heat shock transcription factor A2** |
| **Bpev01.c0230.g0017** | **0** | **0** | **1.784599445** | **AT4G18880** | **HSFA4A** | **Heat shock transcription factor A4A** |
| **Bpev01.c0615.g0008** | **0** | **0** | **2.081664558** | **AT3G22830** | **HSFA6B** | **Heat shock transcription factor A6B** |
| **Bpev01.c0278.g0038** | **1.154610286** | **1.143353775** | **2.420449713** | **AT4G11660** | **HSFB2B** | **Heat shock transcription factor B2B** |
| **Bpev01.c0030.g0025** | **0** | **0** | **2.25590887** | **AT5G02500** | **HSP70-1** | **Heat shock cognate protein 70-1** |
| **Bpev01.c0427.g0008** | **0** | **1.576721052** | **0** | **AT5G02500** | **HSP70-1** | **Heat shock cognate protein 70-1** |
| **Bpev01.c0494.g0001** | **0** | **4.576865357** | **1.949236701** | **AT5G02500** | **HSP70-1** | **Heat shock cognate protein 70-1** |
| **Bpev01.c0495.g0009** | **0** | **2.093487739** | **0** | **AT5G02500** | **HSP70-1** | **Heat shock cognate protein 70-1** |
| **Bpev01.c0634.g0026** | **0** | **0** | **3.757565044** | **AT5G02500** | **HSP70-1** | **Heat shock cognate protein 70-1** |
| **Bpev01.c0634.g0034** | **0** | **2.287259847** | **0** | **AT5G02500** | **HSP70-1** | **Heat shock cognate protein 70-1** |
| **Bpev01.c2531.g0001** | **0** | **0** | **-2.09339376** | **AT5G02500** | **HSP70-1** | **Heat shock cognate protein 70-1** |
| **Bpev01.c2531.g0002** | **0** | **0** | **-1.590876493** | **AT5G02500** | **HSP70-1** | **Heat shock cognate protein 70-1** |
| **Bpev01.c2531.g0003** | **0** | **0** | **-1.659262136** | **AT5G02500** | **HSP70-1** | **Heat shock cognate protein 70-1** |
| **Bpev01.c2761.g0001** | **0** | **4.39324826** | **1.904317037** | **AT5G02500** | **HSP70-1** | **Heat shock cognate protein 70-1** |
| **Bpev01.c0214.g0061** | **0** | **1.209198492** | **0** | **AT4G24190** | **HSP90-7** | **Heat shock protein 90-7** |
| **Bpev01.c0343.g0008** | **0** | **0** | **1.583607275** | **AT4G08170** | **ITPK3** | **Inositol 1,3,4-trisphosphate 5/6-kinase 3** |
| **Bpev01.c0828.g0012** | **0** | **-1.966288013** | **1.912876177** | **AT4G08170** | **ITPK3** | **Inositol 1,3,4-trisphosphate 5/6-kinase 3** |
| **Bpev01.c0118.g0004** | **0** | **3.42470099** | **2.248264215** | **AT4G32650** | **KAT3/KC1** | **Potassium channel in Arabidopsis thaliana 3** |
| **Bpev01.c2670.g0006** | **0** | **0** | **2.022843315** | **AT4G32650** | **KAT3/KC1** | **Potassium channel in Arabidopsis thaliana 3** |
| **Bpev01.c0112.g0010** | **0** | **0** | **-2.242272963** | **AT5G55630** | **KCO1/TPK1** | **Ca2+ activated outward rectifying K+ channel 1/Two pore K channel 1** |
| **Bpev01.c0112.g0011** | **0** | **0** | **-3.836809708** | **AT5G55630** | **KCO1/TPK1** | **Ca2+ activated outward rectifying K+ channel 1/Two pore K channel 1** |
| **Bpev01.c1938.g0005** | **-1.007042849** | **-1.182809782** | **-1.029424821** | **AT5G11800** | **KEA6** | **K+ efflux antiporter 6** |
| **Bpev01.c0177.g0045** | **-3.166828532** | **0** | **-1.34227159** | **AT2G40540** | **KUP/HAK/KT2** | **Potassium transporter 2** |
| **Bpev01.c1083.g0001** | **0** | **0** | **5.703723123** | **AT2G40540** | **KUP/HAK/KT2** | **Potassium transporter 2** |
| **Bpev01.c0720.g0001** | **0** | **2.277972227** | **0** | **AT1G01470** | **LEA14/LSR3** | **Late embryogenesis abundant 14/Light stress-regulated 3** |
| **Bpev01.c0720.g0003** | **1.866209038** | **0** | **2.03496983** | **AT1G01470** | **LEA14/LSR3** | **Late embryogenesis abundant 14/Light stress-regulated 3** |
| **Bpev01.c0045.g0033** | **0** | **2.17882636** | **0** | **AT1G17420** | **LOX3** | **Lipoxygenase 3** |
| **Bpev01.c1365.g0005** | **0** | **0** | **3.354684023** | **AT1G17420** | **LOX3** | **Lipoxygenase 3** |
| **Bpev01.c0875.g0001** | **0** | **-8.957917325** | **0** | **AT5G49760** | **CARD1/HPCA1** | **Cannot respond to DMBQ 1** |
| **Bpev01.c1681.g0005** | **0** | **0** | **2.201167086** | **AT5G49760** | **CARD1/HPCA1** | **Cannot respond to DMBQ 1** |
| **Bpev01.c0146.g0007** | **0** | **2.486346405** | **0** | **AT1G09970** | **RLK7** | **Receptor-like kinase 7** |
| **Bpev01.c0146.g0008** | **0** | **3.054576193** | **0** | **AT1G09970** | **RLK7** | **Receptor-like kinase 7** |
| **Bpev01.c0568.g0002** | **0** | **1.577445343** | **0** | **AT1G09970** | **RLK7** | **Receptor-like kinase 7** |
| **Bpev01.c0264.g0039** | **-1.765555368** | **0** | **-1.87636062** | **AT5G46330** | **FLS2** | **Flagellin-sensitive** |
| **Bpev01.c0000.g0001** | **0** | **-2.076299327** | **0** | **AT4G08850** | **MIK2** | **MDIS1-interacting receptor like kinase 2** |
| **Bpev01.c0776.g0010** | **0** | **3.10736124** | **0** | **AT4G08850** | **MIK2** | **MDIS1-interacting receptor like kinase 2** |
| **Bpev01.c1603.g0005** | **0** | **0** | **1.604913314** | **AT4G08850** | **MIK2** | **MDIS1-interacting receptor like kinase 2** |
| **Bpev01.c1603.g0006** | **0** | **0** | **1.65783928** | **AT4G08850** | **MIK2** | **MDIS1-interacting receptor like kinase 2** |
| **Bpev01.c3151.g0001** | **0** | **5.995206098** | **0** | **AT4G08850** | **MIK2** | **MDIS1-interacting receptor like kinase 2** |
| **Bpev01.c0577.g0035** | **0** | **-1.583835276** | **-1.150425047** | **AT1G31420** | **FEI1** | **FEI 1** |
| **Bpev01.c0064.g0001** | **-1.417844573** | **-2.151664985** | **0** | **AT2G26330** | **ER** | **ERECTA** |
| **Bpev01.c0127.g0022** | **0** | **0** | **1.343688306** | **AT3G21630** | **LYK1/CERK1** | **Chitin elicitor receptor kinase 1/LYSM-containing receptor-like kinase 1** |
| **Bpev01.c1000.g0001** | **0** | **1.527200052** | **1.954498185** | **AT3G21630** | **LYK1/CERK1** | **Chitin elicitor receptor kinase 1/LYSM-containing receptor-like kinase 1** |
| **Bpev01.c0015.g0257** | **0** | **2.153093466** | **0** | **AT2G23770** | **LYK4** | **LYSM-containing receptor-like kinase 4** |
| **Bpev01.c0015.g0258** | **0** | **3.427858601** | **1.819357235** | **AT2G23770** | **LYK4** | **LYSM-containing receptor-like kinase 4** |
| **Bpev01.c0324.g0019** | **0** | **0** | **1.885057682** | **AT2G23770** | **LYK4** | **LYSM-containing receptor-like kinase 4** |
| **Bpev01.c0127.g0041** | **0** | **0** | **2.299273463** | **AT5G55090** | **MAPKKK15** | **Mitogen-activated protein kinase kinase kinase 15** |
| **Bpev01.c0565.g0028** | **0** | **4.638814352** | **0** | **AT5G55090** | **MAPKKK15** | **Mitogen-activated protein kinase kinase kinase 15** |
| **Bpev01.c0274.g0037** | **0** | **1.191094873** | **0** | **AT4G08500** | **MAPKKK8** | **Mitogen-activated protein kinase kinase kinase 8** |
| **Bpev01.c0629.g0009** | **0** | **0** | **1.11314082** | **AT4G08500** | **MAPKKK8** | **Mitogen-activated protein kinase kinase kinase 8** |
| **Bpev01.c0443.g0026** | **0** | **0** | **1.421546608** | **AT4G20260** | **MDP25** | **Microtubule-destabilizing protein 25** |
| **Bpev01.c1798.g0004** | **-1.02641906** | **1.146050555** | **-1.211212995** | **AT1G73500** | **MKK9** | **Mitogen-activated protein kinase kinase 9** |
| **Bpev01.c3183.g0002** | **0** | **0** | **-2.27574596** | **AT1G73500** | **MKK9** | **Mitogen-activated protein kinase kinase 9** |
| **Bpev01.c1036.g0001** | **0** | **2.09900409** | **0** | **AT3G45640** | **MPK3** | **Mitogen-activated protein kinase 3** |
| **Bpev01.c0120.g0044** | **0** | **0** | **1.417457348** | **AT4G01370** | **MPK4** | **Mitogen-activated protein kinase 4** |
| **Bpev01.c0564.g0023** | **0** | **1.746716921** | **0** | **AT3G09390** | **MT2B** | **Metallothionein 2A** |
| **Bpev01.c0564.g0024** | **0** | **0** | **-1.919397403** | **AT3G09390** | **MT2B** | **Metallothionein 2A** |
| **Bpev01.c0066.g0053** | **0** | **-1.032032025** | **-1.13710351** | **AT4G14605** | **mTERF5** | **mTERF defective in Arabidopsis 1** |
| **Bpev01.c1435.g0009** | **0** | **-1.749246035** | **0** | **AT4G14605** | **mTERF5** | **mTERF defective in Arabidopsis 1** |
| **Bpev01.c0135.g0042** | **0** | **-2.325933613** | **0** | **AT1G52890** | **NAP/NAC019** | **NAC domain containing protein 19** |
| **Bpev01.c0162.g0014** | **2.133356046** | **3.106708431** | **3.552409459** | **AT4G27410** | **RD26** | **Responsive to desiccation 26** |
| **Bpev01.c0357.g0068** | **0** | **1.779702938** | **3.47990739** | **AT1G69490** | **NAP/NAC29/NAP** | **NAC domain containing protein 29** |
| **Bpev01.c0294.g0006** | **4.266772067** | **3.238573593** | **2.301255716** | **AT3G14440** | **NCED3** | **Nine-cis-epoxycarotenoid dioxygenase 3** |
| **Bpev01.c0136.g0009** | **-3.215701215** | **0** | **0** | **AT5G56750** | **NDL1** | **N-MYC downregulated-like 1** |
| **Bpev01.c2443.g0001** | **0** | **0** | **-6.362393414** | **AT3G20860** | **Nek6** | **NIMA-related kinase 5** |
| **Bpev01.c0777.g0010** | **0** | **1.682716605** | **0** | **AT5G12840** | **NFYA1/HAP2A** | **Nuclear factor Y, subunit A1** |
| **Bpev01.c0609.g0013** | **0** | **1.635477546** | **0** | **AT5G55470** | **NHX3** | **Na+/H+ (sodium hydrogen) exchanger 3** |
| **Bpev01.c0257.g0003** | **0** | **0** | **2.086148089** | **AT5G62680** | **GTR2/NPF2.11** | **Glucosinolate transporter-2/NRT1/ PTR family 2.11** |
| **Bpev01.c0050.g0160** | **0** | **-4.52764235** | **0** | **AT5G46050** | **NPF5.2/PTR3** | **NRT1/PTR familuy 5.2/Peptide transporter 3** |
| **Bpev01.c0215.g0034** | **0** | **0** | **1.812348927** | **AT5G46050** | **NPF5.2/PTR3** | **NRT1/PTR familuy 5.2/Peptide transporter 3** |
| **Bpev01.c0055.g0015** | **-1.257858667** | **0** | **0** | **AT1G32450** | **NPF7.3/NRT1.5** | **NRT1/PTR family 7.3/Nitrate transporter 1.5** |
| **Bpev01.c1465.g0009** | **0** | **2.556055927** | **3.471937206** | **AT1G32450** | **NPF7.3/NRT1.5** | **NRT1/PTR family 7.3/Nitrate transporter 1.5** |
| **Bpev01.c0796.g0007** | **0** | **1.837643585** | **0** | **AT2G27300** | **ANAC040/NTL8** | **Arabidopsis nac domain containing protein 40/NTM1-like 8** |
| **Bpev01.c0157.g0032** | **0** | **1.073593742** | **-1.985673161** | **AT4G34200** | **PGDH1** | **D-3-phosphoglycerate dehydrogenase** |
| **Bpev01.c1162.g0001** | **0** | **-1.306567861** | **0** | **AT1G70940** | **PIN3** | **PIN-FORMED 3** |
| **Bpev01.c0483.g0002** | **0** | **2.430723486** | **1.852072988** | **AT3G53420** | **PIP2;1** | **Plasma membrane intrinsic protein 2;1** |
| **Bpev01.c0042.g0019** | **0** | **0** | **-1.440668474** | **AT2G37170** | **PIP2;2** | **Plasma membrane intrinsic protein 2;2** |
| **Bpev01.c0142.g0070** | **-1.135061945** | **0** | **2.050419965** | **AT3G16785** | **PLDzeta1** | **Phospholipase D ZETA1** |
| **Bpev01.c0068.g0004** | **0** | **5.776195997** | **0** | **AT5G62360** | **PMEI13** | **Pectin methyl-esterase inhibitor 13** |
| **Bpev01.c0452.g0015** | **2.666605437** | **3.575023755** | **4.314333092** | **AT1G60190** | **PUB19** | **Plant U-box 19** |
| **Bpev01.c0129.g0055** | **0** | **0** | **1.790124196** | **AT2G47190** | **MYB2** | **MYB domain protein 2** |
| **Bpev01.c0145.g0045** | **0** | **1.873282711** | **0** | **AT1G66230** | **MYB20** | **MYB domain protein 20** |
| **Bpev01.c0668.g0007** | **0** | **1.380219931** | **2.614411161** | **AT1G22640** | **MYB3** | **MYB domain protein 3** |
| **Bpev01.c0281.g0058** | **0** | **0** | **-2.885896931** | **AT3G28910** | **MYB30** | **MYB domain protein 30** |
| **Bpev01.c0469.g0003** | **0** | **0** | **4.09191991** | **AT4G28110** | **MYB41** | **MYB domain protein 41** |
| **Bpev01.c0074.g0011** | **0** | **0** | **1.275930042** | **AT4G12350** | **MYB42** | **MYB domain protein 42** |
| **Bpev01.c0115.g0098** | **0** | **0** | **7.719090673** | **AT5G67300** | **MYB44** | **MYB domain protein 44** |
| **Bpev01.c0157.g0038** | **0** | **3.892984582** | **0** | **AT5G67300** | **MYB44** | **MYB domain protein 44** |
| **Bpev01.c0015.g0060** | **0** | **1.515780604** | **0** | **AT4G37260** | **MYB73** | **MYB domain protein 73** |
| **Bpev01.c1972.g0012** | **0** | **3.391508712** | **-1.252381736** | **AT4G37260** | **MYB73** | **MYB domain protein 73** |
| **Bpev01.c2080.g0001** | **0** | **2.535016665** | **-1.301627295** | **AT4G37260** | **MYB73** | **MYB domain protein 73** |
| **Bpev01.c0147.g0014** | **0** | **2.253043176** | **0** | **AT5G47910** | **RBOHD** | **Respiratory burst oxidase homologue D** |
| **Bpev01.c1930.g0004** | **0** | **0** | **-1.277783386** | **AT3G46620** | **RDUF1** | **RING and domain of unknown function 1117 1** |
| **Bpev01.c0542.g0003** | **0** | **-3.145789638** | **0** | **AT3G09770** | **AIRP3** | **ABA insensitive ring protein 3** |
| **Bpev01.c1188.g0001** | **0** | **2.016208635** | **0** | **AT3G09770** | **AIRP3** | **ABA insensitive ring protein 3** |
| **Bpev01.c0123.g0020** | **0** | **0** | **1.442519052** | **AT5G63110** | **HDA6** | **Histone deacetylase 6** |
| **Bpev01.c0752.g0004** | **0** | **-1.788091574** | **0** | **AT4G33470** | **HDA14** | **Histone deacetylase 14** |
| **Bpev01.c2413.g0001** | **0** | **-1.964185783** | **0** | **AT4G33470** | **HDA14** | **Histone deacetylase 14** |
| **Bpev01.c1668.g0003** | **0** | **3.521342007** | **2.71615626** | **AT5G04760** | **DIV2** | **DIVARICATA2** |
| **Bpev01.c0717.g0021** | **0** | **0** | **-1.064976068** | **AT2G45640** | **SAP18** | **SIN3 associated polypeptide P18** |
| **Bpev01.c0878.g0011** | **0** | **0** | **1.782047905** | **AT3G02850** | **SKOR** | **STELAR K+ outward rectifier** |
| **Bpev01.c1055.g0001** | **0** | **0** | **-1.492016988** | **AT3G02850** | **SKOR** | **STELAR K+ outward rectifier** |
| **Bpev01.c1028.g0010** | **0** | **0** | **-1.457688796** | **AT4G24210** | **SLY1** | **SLEEPY1** |
| **Bpev01.c0027.g0152** | **0** | **1.208970207** | **0** | **AT5G66880** | **SnRK2.3** | **Sucrose nonfermenting 1 (SNF1)-related protein kinase 2.3** |
| **Bpev01.c0713.g0001** | **0** | **-1.375914314** | **0** | **AT1G10940** | **SnRK2.4** | **Sucrose nonfermenting 1 (SNF1)-related protein kinase 2.4** |
| **Bpev01.c0000.g0141** | **1.88876579** | **0** | **0** | **AT4G33950** | **SnRK2.6/OST1** | **Sucrose non-fermenting 1 (SNF1)-related protein kinase 2.6/Open stomata 1** |
| **Bpev01.c0222.g0002** | **0** | **0** | **1.443280114** | **AT4G33950** | **SnRK2.6/OST1** | **Sucrose non-fermenting 1 (SNF1)-related protein kinase 2.6/Open stomata 1** |
| **Bpev01.c0479.g0012** | **1.023413049** | **0** | **0** | **AT4G33950** | **SnRK2.6/OST1** | **Sucrose non-fermenting 1 (SNF1)-related protein kinase 2.6/Open stomata 1** |
| **Bpev01.c0090.g0024** | **0** | **1.55916008** | **-1.299613747** | **AT2G30360** | **SnRK3.22/CIPK11/PKS5** | **SNF1-related protein kinase 3.22/CBL-interacting protein kinase 11/Protein kinase SOS2-like 5** |
| **Bpev01.c0497.g0021** | **1.391835868** | **0** | **0** | **AT2G25180** | **ARR12** | **Arabidopsis response regulator 12** |
| **Bpev01.c1126.g0008** | **0** | **0** | **4.362025503** | **AT2G25180** | **ARR12** | **Arabidopsis response regulator 12** |
| **Bpev01.c0332.g0002** | **0** | **0** | **-1.427105185** | **AT1G15690** | **AVP1** | **Arabidopsis thaliana V-PPASE 3** |
| **Bpev01.c0364.g0015** | **0** | **0** | **1.632719756** | **AT1G15690** | **AVP1** | **Arabidopsis thaliana V-PPASE 3** |
| **Bpev01.c0101.g0030** | **0** | **0** | **-1.272432637** | **AT2G02760** | **UBC2** | **Ubiquiting-conjugating enzyme 2** |
| **Bpev01.c0508.g0006** | **0** | **-1.449417017** | **0** | **AT2G02760** | **UBC2** | **Ubiquiting-conjugating enzyme 2** |
| **Bpev01.c1966.g0003** | **0** | **1.130679369** | **0** | **AT1G30620** | **UXE1/MUR4** | **UDP-d-xylose 4-epimerase 1/MURUS4** |
| **Bpev01.c0574.g0015** | **0** | **1.43414974** | **0** | **AT5G67500** | **VDAC2** | **Voltage dependent anion channel 2** |
| **Bpev01.c0439.g0019** | **0** | **1.975680822** | **0** | **AT2G39770** | **VTC1** | **Vitamin c defective 1** |
| **Bpev01.c0879.g0002** | **0** | **0** | **1.523602181** | **AT2G39770** | **VTC1** | **Vitamin c defective 1** |
| **Bpev01.c0155.g0014** | **0** | **0** | **1.512280956** | **AT5G41990** | **WNK8** | **With no lysine (K) kinase 8** |
