## Supplementary material for "Transcriptome analysis provides insights into candidate genetic regulators of birch response to salt stress": Table 5 One inferred GRN relevant for salt stress response uniquely present in leaves of B. platyphylla.

| Table S5 One inferred GRN relevant for salt stress response uniquely present in leaves of B. platyphylla. | | |
| --- | --- | --- |
| **Regulatory gene ID** | **Target gene ID** | **Weight value** |
| **Bpev01.c1085.g0027** | **Bpev01.c0050.g0104** | **0.012158055** |
| **Bpev01.c0555.g0027** | **Bpev01.c1907.g0009** | **0.010819165** |
| **Bpev01.c3305.g0001** | **Bpev01.c0018.g0067** | **0.010802469** |
| **Bpev01.c0120.g0016** | **Bpev01.c1250.g0016** | **0.010638298** |
| **Bpev01.c0015.g0127** | **Bpev01.c0878.g0021** | **0.010590015** |
| **Bpev01.c0170.g0055** | **Bpev01.c0624.g0008** | **0.010494753** |
| **Bpev01.c0413.g0004** | **Bpev01.c0394.g0015** | **0.010279001** |
| **Bpev01.c0047.g0008** | **Bpev01.c0299.g0010** | **0.010279001** |
| **Bpev01.c0120.g0016** | **Bpev01.c0121.g0041** | **0.010115607** |
| **Bpev01.c1966.g0003** | **Bpev01.c0348.g0014** | **0.010028653** |
