## Supplementary material for "Transcriptome analysis provides insights into candidate genetic regulators of birch response to salt stress": Table 6 One inferred GRN relevant for salt stress response uniquely present in roots of B. platyphylla.

| **Table S6 One inferred GRN relevant for salt stress response uniquely present in roots of *B. platyphylla.*** | | |
| --- | --- | --- |
| **Regulatory gene ID** | **Target gene ID** | **Weight value** |
| **Bpev01.c0053.g0059** | **Bpev01.c1484.g0017** | **0.01037037** |
| **Bpev01.c0022.g0007** | **Bpev01.c0556.g0003** | **0.010294118** |
