## Supplementary material for "Transcriptome analysis provides insights into candidate genetic regulators of birch response to salt stress": Table 7 One inferred GRN relevant for salt stress response shared in different tissues of B. halophila and B. platyphylla.

| **Table S7 One inferred GRN relevant for salt stress response shared in different tissues of *B. halophila* and *B. platyphylla.*** | | |
| --- | --- | --- |
| **Regulatory gene ID** | **Target gene ID** | **Weight value** |
| **Bpev01.c0407.g0009** | **Bpev01.c0115.g0062** | **0.024616444** |
| **Bpev01.c1503.g0003** | **Bpev01.c0029.g0102** | **0.032497778** |
| **Bpev01.c1503.g0003** | **Bpev01.c2077.g0003** | **-0.029451444** |
| **Bpev01.c0407.g0009** | **Bpev01.c0416.g0005** | **0.024649333** |
| **Bpev01.c1503.g0003** | **Bpev01.c0115.g0033** | **0.029208889** |
| **Bpev01.c1503.g0003** | **Bpev01.c0161.g0112** | **0.028507778** |
| **Bpev01.c1503.g0003** | **Bpev01.c0062.g0066** | **0.028413333** |
| **Bpev01.c0001.g0024** | **Bpev01.c0278.g0038** | **0.018982778** |
| **Bpev01.c0407.g0009** | **Bpev01.c0891.g0008** | **-0.023105222** |
| **Bpev01.c1503.g0003** | **Bpev01.c0480.g0073** | **0.025401556** |
| **Bpev01.c1503.g0003** | **Bpev01.c0115.g0062** | **0.026156** |
| **Bpev01.c1503.g0003** | **Bpev01.c0177.g0042** | **0.024807333** |
| **Bpev01.c0407.g0009** | **Bpev01.c0528.g0002** | **-0.022359778** |
| **Bpev01.c0088.g0098** | **Bpev01.c0115.g0062** | **-0.027617667** |
| **Bpev01.c1503.g0003** | **Bpev01.c0416.g0005** | **0.027390778** |
| **Bpev01.c0407.g0009** | **Bpev01.c0115.g0033** | **0.021159556** |
| **Bpev01.c1503.g0003** | **Bpev01.c0891.g0008** | **-0.023904667** |
| **Bpev01.c0001.g0024** | **Bpev01.c0141.g0021** | **-0.014251667** |
| **Bpev01.c0162.g0014** | **Bpev01.c0416.g0005** | **0.020291556** |
| **Bpev01.c0088.g0098** | **Bpev01.c0141.g0021** | **0.023456778** |
| **Bpev01.c1503.g0003** | **Bpev01.c0141.g0021** | **-0.027806** |
| **Bpev01.c1503.g0003** | **Bpev01.c0213.g0046** | **-0.021745333** |
| **Bpev01.c0162.g0014** | **Bpev01.c2077.g0003** | **-0.017216** |
| **Bpev01.c0407.g0009** | **Bpev01.c0029.g0102** | **0.019895111** |
| **Bpev01.c0088.g0098** | **Bpev01.c0278.g0038** | **-0.028846667** |
| **Bpev01.c0088.g0098** | **Bpev01.c0051.g0073** | **-0.024553556** |
| **Bpev01.c0469.g0012** | **Bpev01.c1292.g0002** | **0.013088889** |
| **Bpev01.c1503.g0003** | **Bpev01.c0278.g0038** | **0.024323** |
| **Bpev01.c0407.g0009** | **Bpev01.c0001.g0013** | **-0.017528778** |
| **Bpev01.c0407.g0009** | **Bpev01.c0177.g0042** | **0.019812111** |
| **Bpev01.c0407.g0009** | **Bpev01.c1789.g0006** | **0.019095** |
| **Bpev01.c1503.g0003** | **Bpev01.c0051.g0073** | **0.025219444** |
| **Bpev01.c1503.g0003** | **Bpev01.c0661.g0011** | **0.018215333** |
| **Bpev01.c1503.g0003** | **Bpev01.c0023.g0032** | **-0.027840444** |
| **Bpev01.c0001.g0024** | **Bpev01.c1292.g0002** | **0.016175444** |
| **Bpev01.c0407.g0009** | **Bpev01.c0278.g0038** | **0.019151222** |
| **Bpev01.c0407.g0009** | **Bpev01.c2077.g0003** | **-0.018281556** |
| **Bpev01.c0001.g0024** | **Bpev01.c1789.g0006** | **0.014261667** |
| **Bpev01.c0162.g0014** | **Bpev01.c1292.g0002** | **0.015120778** |
| **Bpev01.c1503.g0003** | **Bpev01.c0349.g0020** | **0.023965889** |
| **Bpev01.c0407.g0009** | **Bpev01.c0480.g0073** | **0.018563111** |
| **Bpev01.c0088.g0098** | **Bpev01.c0528.g0002** | **0.023353889** |
| **Bpev01.c0162.g0014** | **Bpev01.c0115.g0062** | **0.016376222** |
| **Bpev01.c0162.g0014** | **Bpev01.c0528.g0002** | **-0.017166444** |
| **Bpev01.c0162.g0014** | **Bpev01.c0029.g0102** | **0.014613444** |
| **Bpev01.c0407.g0009** | **Bpev01.c0213.g0046** | **-0.017784556** |
| **Bpev01.c0920.g0014** | **Bpev01.c0480.g0073** | **0.014290667** |
| **Bpev01.c0407.g0009** | **Bpev01.c0141.g0021** | **-0.020250667** |
| **Bpev01.c0920.g0014** | **Bpev01.c0029.g0102** | **0.013592556** |
| **Bpev01.c0088.g0098** | **Bpev01.c2077.g0003** | **0.025092556** |
| **Bpev01.c1503.g0003** | **Bpev01.c0017.g0006** | **0.020049333** |
| **Bpev01.c0281.g0074** | **Bpev01.c0528.g0002** | **-0.013183778** |
| **Bpev01.c0162.g0014** | **Bpev01.c0480.g0073** | **0.016905444** |
| **Bpev01.c0920.g0014** | **Bpev01.c2077.g0003** | **-0.013057556** |
| **Bpev01.c1503.g0003** | **Bpev01.c0161.g0025** | **0.022740444** |
| **Bpev01.c0950.g0006** | **Bpev01.c0177.g0042** | **0.013828444** |
| **Bpev01.c0950.g0006** | **Bpev01.c1292.g0002** | **0.010421667** |
| **Bpev01.c1503.g0003** | **Bpev01.c0042.g0001** | **0.020252222** |
| **Bpev01.c1503.g0003** | **Bpev01.c0936.g0004** | **0.022491444** |
| **Bpev01.c0162.g0014** | **Bpev01.c1789.g0006** | **0.015622667** |
| **Bpev01.c0407.g0009** | **Bpev01.c0349.g0020** | **0.018079556** |
| **Bpev01.c0001.g0024** | **Bpev01.c0062.g0066** | **0.017201444** |
| **Bpev01.c0088.g0098** | **Bpev01.c0416.g0005** | **-0.022061222** |
| **Bpev01.c0950.g0006** | **Bpev01.c0349.g0020** | **0.011207556** |
| **Bpev01.c0001.g0024** | **Bpev01.c0115.g0033** | **0.015728667** |
| **Bpev01.c0001.g0024** | **Bpev01.c0416.g0005** | **0.010805333** |
| **Bpev01.c0001.g0024** | **Bpev01.c0051.g0073** | **0.016530667** |
| **Bpev01.c0920.g0014** | **Bpev01.c0416.g0005** | **0.012398556** |
| **Bpev01.c0088.g0098** | **Bpev01.c0480.g0073** | **-0.022594111** |
| **Bpev01.c0950.g0006** | **Bpev01.c0141.g0021** | **-0.012503222** |
| **Bpev01.c1503.g0003** | **Bpev01.c0528.g0002** | **-0.022276222** |
| **Bpev01.c1503.g0003** | **Bpev01.c1292.g0002** | **0.019989556** |
| **Bpev01.c0162.g0014** | **Bpev01.c0051.g0073** | **0.013605667** |
| **Bpev01.c0162.g0014** | **Bpev01.c0936.g0004** | **0.015573778** |
| **Bpev01.c0920.g0014** | **Bpev01.c0051.g0073** | **0.012748333** |
| **Bpev01.c0920.g0014** | **Bpev01.c0161.g0025** | **0.012870333** |
| **Bpev01.c0920.g0014** | **Bpev01.c0528.g0002** | **-0.012650222** |
| **Bpev01.c0950.g0006** | **Bpev01.c0278.g0038** | **0.011040778** |
| **Bpev01.c1503.g0003** | **Bpev01.c0015.g0096** | **-0.021056222** |
| **Bpev01.c0162.g0014** | **Bpev01.c0349.g0020** | **0.013739556** |
| **Bpev01.c0407.g0009** | **Bpev01.c0015.g0096** | **-0.019699222** |
| **Bpev01.c0920.g0014** | **Bpev01.c0891.g0008** | **-0.013595444** |
| **Bpev01.c0088.g0098** | **Bpev01.c0062.g0066** | **-0.018564333** |
| **Bpev01.c0088.g0098** | **Bpev01.c1789.g0006** | **-0.026384667** |
| **Bpev01.c0950.g0006** | **Bpev01.c1789.g0006** | **0.011285111** |
| **Bpev01.c1503.g0003** | **Bpev01.c0001.g0013** | **-0.016902556** |
| **Bpev01.c0162.g0014** | **Bpev01.c0001.g0013** | **-0.01346** |
| **Bpev01.c0162.g0014** | **Bpev01.c0141.g0021** | **-0.014508667** |
| **Bpev01.c0407.g0009** | **Bpev01.c0051.g0073** | **0.016408444** |
| **Bpev01.c0001.g0024** | **Bpev01.c0891.g0008** | **-0.014555222** |
| **Bpev01.c0281.g0074** | **Bpev01.c1789.g0006** | **0.011849222** |
| **Bpev01.c0407.g0009** | **Bpev01.c0661.g0011** | **0.016604889** |
| **Bpev01.c0088.g0098** | **Bpev01.c0213.g0046** | **0.021949** |
| **Bpev01.c0088.g0098** | **Bpev01.c0349.g0020** | **-0.019672667** |
| **Bpev01.c0001.g0024** | **Bpev01.c0213.g0046** | **-0.013457444** |
| **Bpev01.c0920.g0014** | **Bpev01.c0115.g0033** | **0.012605** |
| **Bpev01.c0088.g0098** | **Bpev01.c0001.g0013** | **0.018678667** |
| **Bpev01.c0088.g0098** | **Bpev01.c1292.g0002** | **-0.020905111** |
| **Bpev01.c0001.g0024** | **Bpev01.c0115.g0062** | **0.014266556** |
| **Bpev01.c0162.g0014** | **Bpev01.c0278.g0038** | **0.015013667** |
| **Bpev01.c0407.g0009** | **Bpev01.c0023.g0032** | **-0.013389** |
| **Bpev01.c0088.g0098** | **Bpev01.c0015.g0096** | **0.024681778** |
| **Bpev01.c0088.g0098** | **Bpev01.c0177.g0042** | **-0.019855556** |
| **Bpev01.c0950.g0006** | **Bpev01.c0051.g0073** | **0.011372778** |
| **Bpev01.c0407.g0009** | **Bpev01.c1292.g0002** | **0.020111556** |
| **Bpev01.c0469.g0012** | **Bpev01.c0161.g0025** | **0.009569556** |
| **Bpev01.c0469.g0012** | **Bpev01.c0349.g0020** | **0.012228333** |
| **Bpev01.c0001.g0024** | **Bpev01.c0029.g0102** | **0.012555111** |
| **Bpev01.c0001.g0024** | **Bpev01.c0177.g0042** | **0.013430222** |
| **Bpev01.c0162.g0014** | **Bpev01.c0161.g0025** | **0.012914778** |
| **Bpev01.c0281.g0074** | **Bpev01.c0051.g0073** | **0.011621667** |
| **Bpev01.c0407.g0009** | **Bpev01.c0936.g0004** | **0.015567222** |
| **Bpev01.c0920.g0014** | **Bpev01.c0115.g0062** | **0.015242889** |
| **Bpev01.c0920.g0014** | **Bpev01.c1789.g0006** | **0.012031556** |
| **Bpev01.c0950.g0006** | **Bpev01.c0528.g0002** | **-0.012547889** |
| **Bpev01.c0001.g0024** | **Bpev01.c0161.g0025** | **0.014919444** |
| **Bpev01.c0001.g0024** | **Bpev01.c0528.g0002** | **-0.014507222** |
| **Bpev01.c0001.g0024** | **Bpev01.c0936.g0004** | **0.014436889** |
| **Bpev01.c0162.g0014** | **Bpev01.c0015.g0096** | **-0.013867667** |
| **Bpev01.c0407.g0009** | **Bpev01.c0161.g0112** | **0.018795222** |
| **Bpev01.c0088.g0098** | **Bpev01.c0023.g0032** | **0.014581889** |
| **Bpev01.c0469.g0012** | **Bpev01.c0416.g0005** | **0.010807111** |
| **Bpev01.c0469.g0012** | **Bpev01.c0480.g0073** | **0.012002222** |
| **Bpev01.c1503.g0003** | **Bpev01.c1789.g0006** | **0.020921222** |
| **Bpev01.c0001.g0024** | **Bpev01.c0015.g0096** | **-0.012880667** |
| **Bpev01.c0920.g0014** | **Bpev01.c1292.g0002** | **0.012945** |
| **Bpev01.c0088.g0098** | **Bpev01.c0029.g0102** | **-0.019628** |
| **Bpev01.c0088.g0098** | **Bpev01.c0661.g0011** | **-0.012115889** |
| **Bpev01.c0001.g0024** | **Bpev01.c0661.g0011** | **0.012668** |
| **Bpev01.c0001.g0024** | **Bpev01.c2077.g0003** | **-0.014007778** |
| **Bpev01.c0162.g0014** | **Bpev01.c0891.g0008** | **-0.014387** |
| **Bpev01.c0281.g0074** | **Bpev01.c0001.g0013** | **-0.007641111** |
| **Bpev01.c0920.g0014** | **Bpev01.c0015.g0096** | **-0.012893778** |
| **Bpev01.c0920.g0014** | **Bpev01.c0213.g0046** | **-0.009445333** |
| **Bpev01.c0088.g0098** | **Bpev01.c0891.g0008** | **0.021929222** |
| **Bpev01.c0950.g0006** | **Bpev01.c2077.g0003** | **-0.012211444** |
| **Bpev01.c0469.g0012** | **Bpev01.c0015.g0096** | **-0.010405444** |
| **Bpev01.c0407.g0009** | **Bpev01.c0161.g0025** | **0.017787667** |
| **Bpev01.c0920.g0014** | **Bpev01.c0278.g0038** | **0.013660667** |
| **Bpev01.c0950.g0006** | **Bpev01.c0161.g0112** | **0.009159111** |
| **Bpev01.c0469.g0012** | **Bpev01.c0213.g0046** | **-0.010617444** |
| **Bpev01.c1503.g0003** | **Bpev01.c2165.g0004** | **0.020169778** |
| **Bpev01.c0001.g0024** | **Bpev01.c0001.g0013** | **-0.012380778** |
| **Bpev01.c0162.g0014** | **Bpev01.c0115.g0033** | **0.011593444** |
| **Bpev01.c0281.g0074** | **Bpev01.c0161.g0025** | **0.009747222** |
| **Bpev01.c0281.g0074** | **Bpev01.c0349.g0020** | **0.011001778** |
| **Bpev01.c0407.g0009** | **Bpev01.c0062.g0066** | **0.016071889** |
| **Bpev01.c0920.g0014** | **Bpev01.c0042.g0001** | **0.012041111** |
| **Bpev01.c0088.g0098** | **Bpev01.c0042.g0001** | **-0.015839778** |
| **Bpev01.c0088.g0098** | **Bpev01.c0320.g0016** | **-0.018406778** |
| **Bpev01.c0001.g0024** | **Bpev01.c0023.g0032** | **-0.011928** |
| **Bpev01.c0162.g0014** | **Bpev01.c0042.g0001** | **0.013763333** |
| **Bpev01.c0162.g0014** | **Bpev01.c0062.g0066** | **0.011929** |
| **Bpev01.c0407.g0009** | **Bpev01.c0042.g0001** | **0.015086778** |
| **Bpev01.c0920.g0014** | **Bpev01.c0936.g0004** | **0.010751556** |
| **Bpev01.c0088.g0098** | **Bpev01.c0115.g0033** | **-0.019555111** |
| **Bpev01.c0162.g0014** | **Bpev01.c0161.g0112** | **0.010846556** |
| **Bpev01.c0162.g0014** | **Bpev01.c0213.g0046** | **-0.013906222** |
| **Bpev01.c0407.g0009** | **Bpev01.c2165.g0004** | **0.015917333** |
| **Bpev01.c0920.g0014** | **Bpev01.c0001.g0013** | **-0.010406** |
| **Bpev01.c0950.g0006** | **Bpev01.c0062.g0066** | **0.011258778** |
| **Bpev01.c1503.g0003** | **Bpev01.c0536.g0006** | **0.013871444** |
| **Bpev01.c0162.g0014** | **Bpev01.c0661.g0011** | **0.010019333** |
| **Bpev01.c0281.g0074** | **Bpev01.c0278.g0038** | **0.009491556** |
| **Bpev01.c0920.g0014** | **Bpev01.c0062.g0066** | **0.010969444** |
| **Bpev01.c0281.g0074** | **Bpev01.c0115.g0033** | **0.009971667** |
| **Bpev01.c0281.g0074** | **Bpev01.c0161.g0112** | **0.009131** |
| **Bpev01.c0281.g0074** | **Bpev01.c1292.g0002** | **0.010356556** |
| **Bpev01.c0920.g0014** | **Bpev01.c0023.g0032** | **-0.012303222** |
| **Bpev01.c0469.g0012** | **Bpev01.c0029.g0102** | **0.009835222** |
| **Bpev01.c0469.g0012** | **Bpev01.c0161.g0112** | **0.011329222** |
| **Bpev01.c0281.g0074** | **Bpev01.c0480.g0073** | **0.008855111** |
| **Bpev01.c0407.g0009** | **Bpev01.c0017.g0006** | **0.013599889** |
| **Bpev01.c0920.g0014** | **Bpev01.c0161.g0112** | **0.010326444** |
| **Bpev01.c0088.g0098** | **Bpev01.c0936.g0004** | **-0.018076889** |
| **Bpev01.c0950.g0006** | **Bpev01.c0213.g0046** | **-0.009629556** |
| **Bpev01.c0950.g0006** | **Bpev01.c0480.g0073** | **0.011614667** |
| **Bpev01.c0469.g0012** | **Bpev01.c2077.g0003** | **-0.009798667** |
| **Bpev01.c0088.g0098** | **Bpev01.c0161.g0112** | **-0.014681778** |
| **Bpev01.c0469.g0012** | **Bpev01.c0062.g0066** | **0.009317222** |
| **Bpev01.c0001.g0024** | **Bpev01.c0161.g0112** | **0.012903222** |
| **Bpev01.c0001.g0024** | **Bpev01.c0349.g0020** | **0.012366444** |
| **Bpev01.c0162.g0014** | **Bpev01.c0177.g0042** | **0.013039667** |
| **Bpev01.c0281.g0074** | **Bpev01.c0023.g0032** | **-0.009145778** |
| **Bpev01.c0088.g0098** | **Bpev01.c2165.g0004** | **-0.017118778** |
| **Bpev01.c0281.g0074** | **Bpev01.c0416.g0005** | **0.008782444** |
| **Bpev01.c0281.g0074** | **Bpev01.c2077.g0003** | **-0.009173** |
| **Bpev01.c0920.g0014** | **Bpev01.c0141.g0021** | **-0.011721** |
| **Bpev01.c0950.g0006** | **Bpev01.c0115.g0033** | **0.009962222** |
| **Bpev01.c0469.g0012** | **Bpev01.c0891.g0008** | **-0.008364333** |
| **Bpev01.c0001.g0024** | **Bpev01.c0042.g0001** | **0.012943778** |
| **Bpev01.c0162.g0014** | **Bpev01.c0023.g0032** | **-0.012210667** |
| **Bpev01.c0281.g0074** | **Bpev01.c0029.g0102** | **0.010064111** |
| **Bpev01.c0950.g0006** | **Bpev01.c0936.g0004** | **0.011427444** |
| **Bpev01.c0469.g0012** | **Bpev01.c0278.g0038** | **0.009237** |
| **Bpev01.c0469.g0012** | **Bpev01.c0528.g0002** | **-0.010511333** |
| **Bpev01.c0469.g0012** | **Bpev01.c1789.g0006** | **0.008062778** |
| **Bpev01.c0950.g0006** | **Bpev01.c0161.g0025** | **0.009623778** |
| **Bpev01.c0162.g0014** | **Bpev01.c2165.g0004** | **0.011449889** |
| **Bpev01.c0281.g0074** | **Bpev01.c0141.g0021** | **-0.007482444** |
| **Bpev01.c0950.g0006** | **Bpev01.c0029.g0102** | **0.010902667** |
| **Bpev01.c0469.g0012** | **Bpev01.c0936.g0004** | **0.008983222** |
| **Bpev01.c0469.g0012** | **Bpev01.c0001.g0013** | **-0.009128889** |
| **Bpev01.c0001.g0024** | **Bpev01.c0017.g0006** | **0.010046** |
| **Bpev01.c0281.g0074** | **Bpev01.c0936.g0004** | **0.008119778** |
| **Bpev01.c0950.g0006** | **Bpev01.c0416.g0005** | **0.009862222** |
| **Bpev01.c0469.g0012** | **Bpev01.c0177.g0042** | **0.009242333** |
| **Bpev01.c1503.g0003** | **Bpev01.c0577.g0019** | **0.014582667** |
| **Bpev01.c0281.g0074** | **Bpev01.c0115.g0062** | **0.009117333** |
| **Bpev01.c0088.g0098** | **Bpev01.c0161.g0025** | **-0.019323** |
| **Bpev01.c0469.g0012** | **Bpev01.c0141.g0021** | **-0.007135111** |
| **Bpev01.c0001.g0024** | **Bpev01.c0480.g0073** | **0.012382889** |
| **Bpev01.c0281.g0074** | **Bpev01.c0177.g0042** | **0.008381222** |
| **Bpev01.c0920.g0014** | **Bpev01.c0177.g0042** | **0.010137889** |
| **Bpev01.c0950.g0006** | **Bpev01.c0001.g0013** | **-0.007921889** |
| **Bpev01.c0950.g0006** | **Bpev01.c0015.g0096** | **-0.009504556** |
| **Bpev01.c0088.g0098** | **Bpev01.c0577.g0019** | **-0.015221222** |
| **Bpev01.c0162.g0014** | **Bpev01.c0017.g0006** | **0.010265667** |
| **Bpev01.c0281.g0074** | **Bpev01.c0213.g0046** | **-0.009768222** |
| **Bpev01.c0920.g0014** | **Bpev01.c0349.g0020** | **0.010096** |
| **Bpev01.c1503.g0003** | **Bpev01.c0022.g0051** | **0.010883556** |
| **Bpev01.c1503.g0003** | **Bpev01.c0320.g0016** | **0.013749111** |
| **Bpev01.c0281.g0074** | **Bpev01.c0042.g0001** | **0.006860889** |
| **Bpev01.c0407.g0009** | **Bpev01.c0577.g0019** | **0.012969** |
| **Bpev01.c0950.g0006** | **Bpev01.c0023.g0032** | **-0.007983222** |
| **Bpev01.c0469.g0012** | **Bpev01.c0023.g0032** | **-0.006882889** |
| **Bpev01.c0281.g0074** | **Bpev01.c0661.g0011** | **0.005961222** |
| **Bpev01.c0281.g0074** | **Bpev01.c2165.g0004** | **0.008539111** |
| **Bpev01.c0950.g0006** | **Bpev01.c2165.g0004** | **0.008229333** |
| **Bpev01.c0469.g0012** | **Bpev01.c0051.g0073** | **0.008339** |
| **Bpev01.c0469.g0012** | **Bpev01.c0115.g0033** | **0.010615667** |
| **Bpev01.c0088.g0098** | **Bpev01.c0017.g0006** | **-0.012661556** |
| **Bpev01.c0088.g0098** | **Bpev01.c0536.g0006** | **-0.010832222** |
| **Bpev01.c0281.g0074** | **Bpev01.c0015.g0096** | **-0.007177667** |
| **Bpev01.c0920.g0014** | **Bpev01.c0320.g0016** | **0.010309111** |
| **Bpev01.c0950.g0006** | **Bpev01.c0017.g0006** | **0.008001444** |
| **Bpev01.c0950.g0006** | **Bpev01.c0115.g0062** | **0.008406556** |
| **Bpev01.c0950.g0006** | **Bpev01.c0891.g0008** | **-0.007565778** |
| **Bpev01.c1503.g0003** | **Bpev01.c0088.g0110** | **0.011443556** |
| **Bpev01.c0281.g0074** | **Bpev01.c0062.g0066** | **0.007668** |
| **Bpev01.c0407.g0009** | **Bpev01.c0320.g0016** | **0.011738333** |
| **Bpev01.c0950.g0006** | **Bpev01.c0320.g0016** | **0.010292667** |
| **Bpev01.c0407.g0009** | **Bpev01.c0088.g0110** | **0.009828667** |
| **Bpev01.c0088.g0098** | **Bpev01.c0088.g0110** | **-0.009431333** |
| **Bpev01.c0001.g0024** | **Bpev01.c2165.g0004** | **0.010789556** |
| **Bpev01.c0920.g0014** | **Bpev01.c0017.g0006** | **0.007800667** |
| **Bpev01.c0950.g0006** | **Bpev01.c0577.g0019** | **0.010302333** |
| **Bpev01.c0469.g0012** | **Bpev01.c0320.g0016** | **0.006899** |
| **Bpev01.c0469.g0012** | **Bpev01.c2165.g0004** | **0.007716444** |
| **Bpev01.c0469.g0012** | **Bpev01.c0661.g0011** | **0.008021222** |
| **Bpev01.c0162.g0014** | **Bpev01.c0577.g0019** | **0.008885444** |
| **Bpev01.c0407.g0009** | **Bpev01.c0536.g0006** | **0.008246111** |
| **Bpev01.c0088.g0098** | **Bpev01.c0022.g0051** | **-0.012949556** |
