## Supplementary material for "Transcriptome analysis provides insights into candidate genetic regulators of birch response to salt stress": Quantitative related information

1、Quantitative experimental design：

| Sample | 0 h | 24 h |
| --- | --- | --- |
| leaf | 3 biological replicates | 3 biological replicates |
| Root | 3 biological replicates | 3 biological replicates |

2、Samples

The plant materials were from the National Key Laboratory of Forest Genetics and Breeding of Northeast Forestry University.

3、Nucleic acid extraction

The method is CTAB method, and the concentration and integrity have no information.

4、Reverse transcription

The first-strand cDNA products of reverse transcription were achieved from 1 μg of DNA-free RNA using the ReverTreAce^®^ qPCR RT Kit (Toyobo, Osaka, Japan).

5、Target gene information

| Gene ID (Gene name) | Contig ID | Start | End |
| --- | --- | --- | --- |
| Bpev01.c0162.g0014 (RD26) | Contig162 | 180205 | 182259 |
| Bpev01.c0088.g0098 (ANAC2) | Contig88 | 1068650 | 1070728 |
| Bpev01.c0278.g0038 (HSFB2B) | Contig278 | 473250 | 475674 |
| Bpev01.c1292.g0002 (EDL3) | Contig1292 | 19814 | 21140 |
| Bpev01.c0480.g0073 (GALS1) | Contig480 | 526349 | 528682 |
| Bpev01.c1938.g0005 (KEA6) | Contig1938 | 44704 | 56731 |
| Bpev01.c1798.g0004 (MKK9) | Contig1798 | 82517 | 83665 |
| Bpev01.c1966.g0003 (UXE1/MUR4) | Contig1966 | 20008 | 27851 |
| Bpev01.c0022.g0007 (Histone H1-3) | Contig22 | 74218 | 75365 |

6、The primer sequence is in supplementary material 1. No other information。

7、qPCR method

Quantitative real-time PCR (RT-qPCR) assay was carried out using the quantitative SYBR green PCR Master Mix (Toyobo, Osaka, Japan). The RT-qPCR assay mixtures, consisting of 1 μL cDNA, 5 μL SYBR Green PCR Master Mix (Toyobo, Osaka, Japan), 1 μL of forward primer, 1 μL of reverse primer, and ultrapure water, were made. qRT-PCR reaction was performed under the condition of 95 ℃ for 30 s, 45 cycles of 95 ℃ for 30 s, 95 ℃ for 15 s, and 58 ℃ for 40 s on an ABI 7500 Real-Time PCR system (USA).

8、There is no information in the result validation section

9、Data analysis

The expression fold-change of each gene relative to a housekeeping gene of *18S rRNA* (GeneBank Accession #: FJ228477.1) was calculated using 2^－△△CT^ method (Schmittgen and Livak 2008).
